# Rapid screening and detection of inter-type viral recombinants using phylo-*k*-mers

**DOI:** 10.1101/2020.06.22.161422

**Authors:** Guillaume E. Scholz, Benjamin Linard, Nikolai Romashchenko, Eric Rivals, Fabio Pardi

## Abstract

**Motivation:** Novel recombinant viruses may have important medical and evolutionary significance, as they sometimes display new traits not present in the parental strains. This is particularly concerning when the new viruses combine fragments coming from phylogenetically-distinct viral types. Here, we consider the task of screening large collections of sequences for such novel recombinants. A number of methods already exist for this task. However, these methods rely on complex models and heavy computations that are not always practical for a quick scan of a large number of sequences.

**Results:** We have developed SHERPAS, a new program to detect novel recombinants and provide a first estimate of their parental composition. Our approach is based on the precomputation of a large database of “phylogenetically-informed *k*-mers”, an idea recently introduced in the context of phylogenetic placement in metagenomics. Our experiments show that SHERPAS is hundreds to thousands of times faster than existing software, and enables the analysis of thousands of whole genomes, or long sequencing reads, within minutes or seconds, and with limited loss of accuracy.

**Availability and Implementation:** The source code is freely available for download at https://github.com/phylo42/sherpas

**Supplementary information:** Supplementary Materials are available online.

## 1 Introduction

A fundamental task in viral bioinformatics is to recognize when a newly sequenced virus genome or genome fragment is a recombinant —that is, it carries regions from two or more genetically distinct parental strains. Detecting novel recombinant forms has important biological and medical implications, as the new recombinants are sometimes associated with drug resistance (Moutouh *et al.*, 1996), increased virulence (Liu *et al.*, 2002; Suarez *et al.*, 2004), the ability to infect new hosts (Kuiken *et al.*, 2006) or to evade the host’s immune system (Streeck *et al.*, 2008). Moreover, for many viral species, recombination is common: for example in HIV the rate of within-host recombination appears to be at least as high as that of point mutations (Neher and Leitner, 2010; Batorsky *et al.*, 2011). Interestingly, a number of artefacts (e.g. caused by PCR amplification or sequence assembly errors) can also result in recombinant sequences, which however never really existed in vivo (Martin *et al.*, 2011; Pérez-Losada *et al.*, 2015). Detecting such artificial recombinants is also important prior to any further sequence analysis.

A virus species is often subdivided into phylogenetically-distinct strains, sometimes called *groups, types* or *subtypes* (the nomenclature varies depending on the virus), representing the diversity of the genomes from that virus. For example, HIV-1 is divided into 4 groups (M, N, O and P) and the M group, responsible for the HIV pandemic, is further classified into at least 9 subtypes (A, B, C, D, F, G, H, J, K), some of which have sub-subtypes (Foley *et al.*, 2018). Here, we use the word *strain* to designate any subset of interest for the virus under consideration. Different strains are sometimes associated to important differences, for example in resistance to antiviral drugs (Wainberg and Brenner, 2010) or in disease progression (Kiguoya *et al.*, 2017).

In this paper, we focus on the computational task of recognizing novel recombinants composed of genomic regions coming from different strains (for example from different subtypes in the case of HIV-1). Given a collection of query sequences, we wish to identify inter-strain recombinants, and for each putative recombinant: (1) recognize which strains originated it; (2) partition it into the regions coming from different strains. Fig. 1 shows an example of the type of information that we intend to recover from a query.

**Fig. 1.**
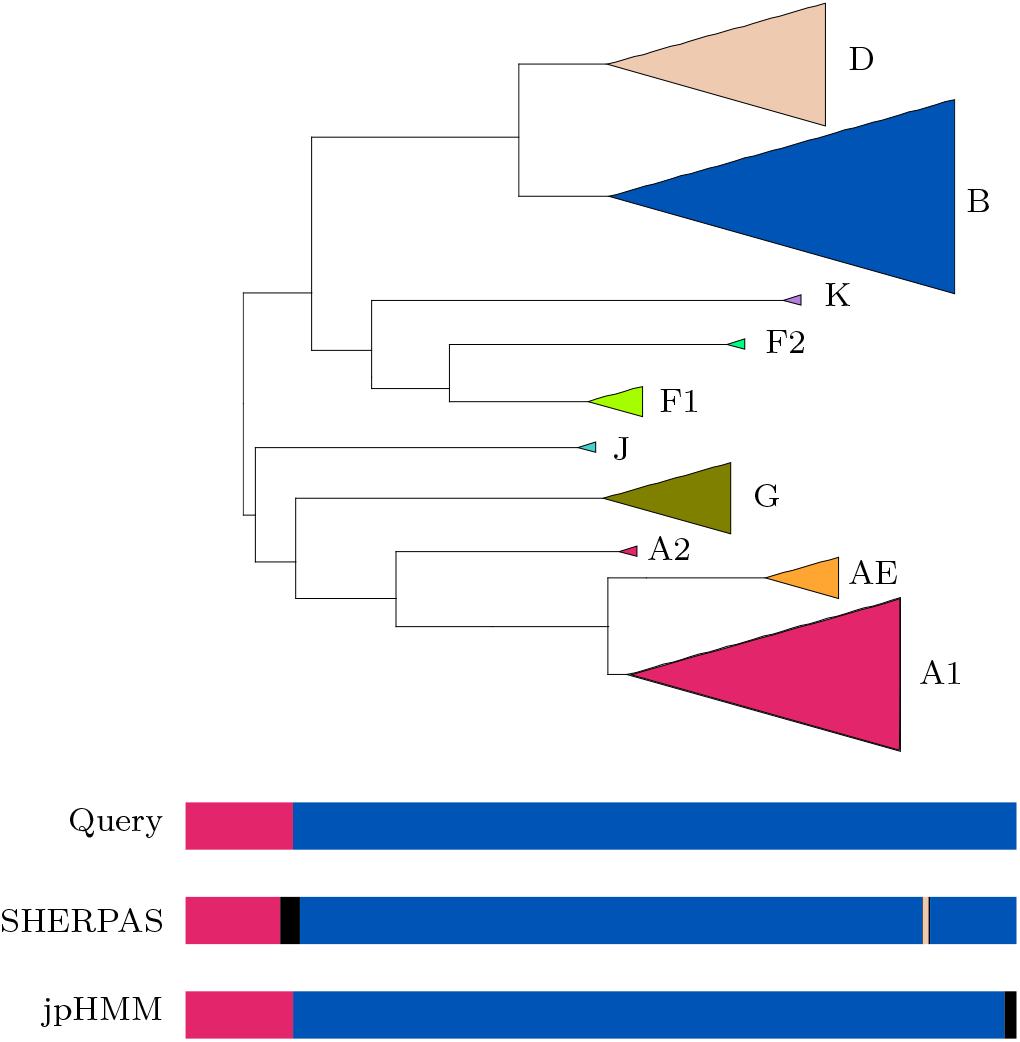
Illustration of the task of inter-strain recombination detection. Top: Example of what strains may look like in a realistic phylogeny (adapted from part of the reference tree for the HIV-pol dataset). Bottom: Illustration of the composition of a query and of the outputs of two programs. The query combines a small segment of a sequence annotated as A1, and a larger segment of a sequence annotated as B. (Neither of these two sequences were part of the reference alignment used to construct the reference tree.) SHERPAS and jpHMM (both run with default parameters) return the partitions represented by the other two bars. Black segments represent unassigned regions.

A number of tools can already be used precisely for this task. For example, jpHMM (Schultz *et al.*, 2006, 2009) —which partitions each query by “jumping” between profile HMMs constructed for the different strains—, SCUEAL (Kosakovsky Pond *et al.*, 2009) —a likelihood-based genetic algorithm— and the REGA subtyping tool (de Oliveira *et al.*, 2005) —which implements a sliding-window-based phylogenetic bootstrap analysis (bootscanning) for HIV-1. All these approaches use a reference alignment containing several representative sequences from each strain. They either need to align the queries to the reference alignment prior to the analysis (SCUEAL and REGA) or they implicitly construct an alignment during their execution (jpHMM). Sometimes the query alignment phase is followed by a phylogenetic analysis step (SCUEAL and REGA), which may have to be repeated over many different portions of the alignment. Because of the complexity of the computations involved, the execution of these tools may become tricky when the datasets to analyse contain more than a few thousands queries.

Because rapidly evolving sequencing technologies enable researchers and clinicians to routinely produce increasingly large sequence datasets —potentially containing millions of viral reads— we have developed a fast alignment-free method to detect inter-strain recombinants within large collections of queries, based on the use of phylo-*k*-mers (Linard *et al.*, 2019) (see Sec. 2.2). The new tool, called SHERPAS (*Screening Historical Events of Recombination in a Phylogeny via Ancestral Sequences*) is able to process thousands of long queries (potentially covering whole viral genomes) within minutes or seconds. It can be used as a tool to screen large sequence datasets for novel recombinants. If necessary, the putative recombinants found by SHERPAS can be subsequently re-analysed with more precise methods such as REGA, SCUEAL or jpHMM.

Beside being orders of magnitude faster than available tools for the discovery of novel recombinants, SHERPAS presents other points of interest. Unlike some popular web interfaces, the code of SHERPAS is distributed freely, which may be an advantage when, for privacy reasons, it is important to process the data in-house (e.g. in a clinical setting). This also makes SHERPAS very flexible: users can choose their own reference alignments, update them as new high-quality sequences become available, and most importantly adapt SHERPAS to any virus for which a reference alignment of sufficient quality can be obtained. Moreover, SHERPAS appears to be relatively robust to the high error rates that characterize Oxford Nanopore sequencers. For these reasons we believe that SHERPAS is appropriate for recombination detection even in the most challenging scenarios, such as in-situ outbreak monitoring, where computational resources and network accessibility may be limited (Quick *et al.*, 2016).

## 2 Algorithm

### 2.1 Preprocessing and overview

At a preprocessing stage, SHERPAS needs a collection of aligned reference sequences for the virus of interest and a phylogenetic tree built from this alignment. Each reference sequence must be annotated as belonging to exactly one strain, via a .csv file. In the Suppl. Materials (Sec. 3), we discuss a number of properties that we would ideally expect the references (alignment, tree and strains) to satisfy, such as the monophyly of strains and the absence of widespread recombination within the reference alignment. From the reference alignment and tree, a database of phylo-*k*-mers (the *pkDB*) is then constructed using the pkDB construction step currently implemented in the RAPPAS software (Linard *et al.*, 2019) (see next section). The pkDB construction is a heavy computational step, but it only needs to be executed when a new reference alignment is employed, or when it is updated.

Once these preprocessing steps have been carried out, large datasets of unaligned DNA sequences can be analyzed with the pkDB, as they become available. These sequences —which we refer to as *queries*— can be genomic fragments of moderate size (a few hundreds bp at least) up to entire genomes, including error-prone long reads generated by third-generation sequencing technologies.

The output of SHERPAS is a text file classifying continuous regions within the queries as either unassigned (“N/A”) or as belonging to one of the strains. The same format used by jpHMM is adopted.

### 2.2 The phylo-*k*-mers

Informally, phylo-*k*-mers can be described as phylogenetically-informed *k*-mers (subsequences of length *k*) that are present with non-negligible probability in unknown/unsampled relatives of the sequences contained in the reference alignment (Linard *et al.*, 2019). Importantly, phylo-*k*-mers are *inferred* from the reference data (alignment and tree), but not necessarily observed in any of the reference sequences. Typical values for *k* are currently in the range from 8 to 10. While a detailed mathematical treatment is deferred to the Suppl. Materials (Sec. 1), here we provide an overview.

The inference of phylo-*k*-mers relies on standard techniques that can calculate the posterior probability of the nucleotide state at (i) any site defined by a column of the reference alignment, and at (ii) any node with a well-defined location with respect to the reference tree. While in traditional applications, such as ancestral sequence reconstruction, the focus is on the internal nodes of the tree, here we are interested in the probabilities at new nodes that are added to the reference tree. These nodes, called *ghost nodes*, represent sequences that have diverged from a given branch, and lie at pre-defined distances from their branch of origin.

Posterior probability calculations are implemented in many programs for likelihood-based phylogenetics (e.g. Guindon and Gascuel (2003); Kozlov *et al.* (2019)), one of which is executed automatically at launch of the phylo-*k*-mer construction step. For each ghost node *u*, this step produces a table containing the posterior distribution of the nucleotide at *u*, at any site of the reference alignment.

The probability of a *k*-mer *w*, at a specific ghost node *u* and at a specific set of *k* consecutive sites, is then obtained as the product of the posterior probabilities of its constituent nucleotides at their respective sites, in the table for node *u*. This simple calculation relies on the assumption of statistical independence among sites, which is standard in phylogenetics (e.g. Felsenstein (2004); Yang (2006)).

A *k*-mer *w* is called a *phylo-k-mer* for branch *x* of the reference tree, if there exists at least one position in the reference alignment and one ghost node associated to *x*, where the probability of *w* exceeds a given threshold (controlled by a parameter of the phylo-*k*-mer construction process). When multiple such positions and ghost nodes exist for a given pair (*w,x*), the highest probability is the *probability score* of *k*-mer *w* at branch *x*. A *k*-mer’s probability score at *x* can be interpreted as a measure of how likely *x* is to be the *k*-mer’s “phylogenetic origin” —that is, the branch from which the *k*-mer diverged from the rest of the reference tree.

Finally, note that a *k*-mer *w* can be a phylo-*k*-mer for several branches, although with potentially very different probability scores. All such information is stored in the pkDB, which is a look-up table allowing, for a given phylo-*k*-mer *w*, the rapid retrieval of all branches and probability scores associated to *w*.

### 2.3 Full and reduced pkDBs

Prior to applying the algorithm for recombination detection, outlined below, each branch of the reference tree is assigned at most one strain from the user-specified set of strains, in the following way: Recall that each reference sequence belongs to exactly one of these strains. If all the sequences that descend from a branch belong to the same strain, then this branch gets assigned a label corresponding to that strain, otherwise the branch remains unassigned. Moreover, we call a branch *x* a *root branch* of strain *X* if (1) *x* is assigned to *X*, and (2) no branch ancestral to *x* is assigned to *X*. Note that if a strain *X* is monophyletic (which we expect to be usually the case), then *X* has exactly one root branch, the one lying at the root of the clade containing all sequences in *X*.

From there, two distinct versions of the pkDB can be constructed. The *full* pkDB is the one constructed by the phylo-*k*-mer inference step currently implemented in RAPPAS, without modification. The *reduced* pkDB is constructed by SHERPAS from the full pkDB, by only keeping the information relative to the branches that are root branches of some strain *X*. See the Suppl. Materials (Sec. 1.5) for more details. We call SHERPAS-full and SHERPAS-reduced the two variants of SHERPAS using the full pkDB (default) and the reduced pkDB, respectively.

### 2.4 The sliding window approach

The recombination detection phase in SHERPAS adopts a sliding window approach. Here, a *window* is defined as a contiguous subsequence of the query of a given length. For each window, instead of performing complex phylogenetic analyses, SHERPAS only looks for matches between the *k*-mers contained in the window and the selected pkDB (full or reduced), and dynamically updates a table of scores associated to the branches encountered in this process. We refer to the Suppl. Materials (Sec. 2) for a detailed description and analysis of the algorithm, and provide the main ideas below.

For each window, the score assigned to a branch is computed using the same weighted vote approach as in RAPPAS’s placement algorithm (Linard *et al.*, 2019). This score is a function of the probability scores at that branch, of the *k*-mers in the window. The scores for the first (leftmost) window are used to initialize a table of scores. For each subsequent window, the table of scores is updated efficiently on the basis of the *k*-mers that are added to it, and those that are removed from it. The number of *k*-mers that are added to the new window does not need to coincide with the number of *k*-mers that are removed from it. This is used to improve the behavior of SHERPAS at the ends of the query: while the leftmost and the rightmost window are relatively small (100 *k*-mers by default), the window gradually grows as it gets further from the ends of the query, until it reaches its maximum size (300 *k*-mers by default). By default, the coordinates of two consecutive windows of maximum size only differ by 1 bp.

SHERPAS is also able to process circular queries, which may arise for viruses with circular genomes. In this case, the variable-size approach described above is not executed. Instead, the sliding window retains the same size everywhere. When the sliding window reaches the end of the query, it will extend to the other end of the query, until the sliding window is back to the leftmost window in the query.

Assuming that the signal for classification is strong enough (see the next section for details), the midpoint in each window is classified into the strain that is associated to the highest-scoring branch for the window. This allows SHERPAS to partition the query into segments, each one associated with a strain identified as its origin.

### 2.5 Signal evaluation and unassigned regions

SHERPAS may leave some parts of a query unassigned, whenever the evidence for the classification into any particular strain is deemed to be too weak. In order to evaluate this, SHERPAS converts the score of a branch into a likelihood score (details of this conversion are provided in the Suppl. Materials, Sec. 2.4). The way this is used depends on the version of the pkDB (full/reduced).

In its full version, the pkDB contains all the branches of the reference phylogeny, including some branches that are not assigned to any strain. If the best scoring branch in a window is one of these unassigned branches, then SHERPAS classifies the window midpoint as unassigned (or “N/A”). If instead the best and second-best scoring branch belong to the same strain, SHERPAS classifies the midpoint in that strain. In all remaining cases, SHERPAS computes the ratio *l*_1_/*l*_2_, where *l*_1_ and *l*_2_ are the likelihoods for the best and second-best branch, respectively. If that ratio is smaller than a user-defined parameter *θ_F_*, SHERPAS classifies the window midpoint as unassigned, otherwise it classifies it in the strain of the best scoring branch.

In the reduced version, all branches recorded in the pkDB belong to some strain (usually just one branch per strain). In that case, SHERPAS computes the ratio 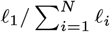, where *l*_1_ is the likelihood for the best scoring branch/strain and *l_i_, i* = 2,…,*N* are the likelihoods of all other branches/strains in the pkDB. Again, if that ratio is smaller than a user-defined parameter *θ_R_* ∈ [0,1), SHERPAS returns the window midpoint as unassigned (or “N/A”).

In both SHERPAS-full and SHERPAS-reduced, setting the control parameter *θ_F_* (or *θ_R_*) to a small value is expected to result in a liberal classification, potentially resulting in false positive breakpoints, while setting it to a high value corresponds to a more conservative classification, potentially missing some evidence of recombination. A last optional step that is applied by SHERPAS is the removal of N/A stretches between two segments classified in the same strain (by default, these regions are classified as belonging to that strain).

An interesting observation is that, since by default two consecutive windows only differ by two *k*-mers, it is very unlikely that their midpoints are both confidently assigned to different strains. Because of this, two genomic regions classified into different strains *X* and *Y* are usually separated by a N/A fragment, which can be interpreted as expressing uncertainty about the precise location of the breakpoint between *X* and *Y*. In other words, we expect the breakpoint *X/Y* to lie somewhere within this N/A fragment.

## 3 Materials and methods

### 3.1 Experimental protocol overview

#### Dataset construction

We evaluated the performance of SHERPAS on four datasets of synthetic recombinants, that is, query sequences that are constructed by concatenating fragments of real-world viral sequences. The first three datasets were obtained following the same general procedure: Each dataset is constructed from a different pair of alignments containing real-world sequences reliably annotated as belonging to known strains of a virus of interest (details in Sec. 3.3 to 3.5). One of these alignments is used as the reference alignment for SHERPAS. The sequences in the other alignment are called *pre-queries*. We ensure the two alignments contain no sequence in common. The pre-queries are used to build a large collection of queries by (1) drawing random recombination breakpoints in the alignment containing the pre-queries, (2) cutting the pre-queries at those breakpoints and (3) concatenating the resulting fragments. The fourth dataset was obtained by simulating long-read sequencing errors over the queries of one of the other datasets (Sec. 3.6). For each of the queries, we record the positions of the breakpoints, and the strain of origin of the fragments that are separated by those breakpoints. This recorded information is used as “ground truth” to evaluate the accuracy of the tested methods (see Sec. 3.2).

#### Software comparison

We compare the performance (accuracy and running times) of SHERPAS over these datasets against that of jpHMM (Schultz *et al.*, 2006, 2009), a natural choice because (1) it is the only tool whose main stated goal is the same as that of SHERPAS (detect *inter*-strain recombinants and partition them according to the strain of origin). Moreover, (2) jpHMM is not specialized for any single virus species, and is distributed with its own reference alignments for a number of viruses, which allows us to compare it to SHERPAS using the same reference alignments. Note that using the same reference alignment (essentially a training set) puts two tools on an equal ground for benchmarking purposes, allowing us to evaluate the relative merits of the algorithms alone —and exclude the influence of the reference data, which is potentially crucial (Pineda-Peña *et al.*, 2013). Also note that, when run with the-Q blat option to speed up its execution, jpHMM appears to be at least as fast as SCUEAL and REGA (Pineda-Peña *et al.*, 2013), thus providing a good comparison for running times. Those alternatives to jpHMM were excluded for the following reasons: SCUEAL (Kosakovsky Pond *et al.*, 2009) is specialized for the detection of HIV-1 recombinants, including *intra*subtype recombinants, and is only distributed with a single reference alignment (for the *pol* gene). The REGA tool (de Oliveira *et al.*, 2005) has only been developed for HIV-1, and does not give access to its code. Since it cannot be run on a local machine, it is not possible to perform fair running-time comparisons with it. All these exclusion criteria also apply to COMET (Struck *et al.*, 2014), a web-based subtyping tool for HIV-1, whose main goal is not recombination analysis.

### 3.2 Measures of accuracy

To measure the accuracy of SHERPAS and of the other methods, we used two approaches: a site-wise and a mosaic approach.

#### Site-wise approach

Since the composition of synthetic recombinant queries is known, we can see such composition as a site-wise assignment. It is then possible to compare the assignment of a site by a recombination-detection software with the correct assignment of that site. We use two different measures of the accuracy of a software: we compute the proportion of sites that are assigned to the correct strain, either out of all sites —the **site-wise sensitivity**— or out of all sites that are not assigned to N/A —the **site-wise precision**. We note that this is a slight abuse of vocabulary, as in multi-class classification, precision and sensitivity are class-specific measures. (See the Suppl. Materials, Sec. 4 for a mathematical reconciliation between these definitions.) In the absence of N/A regions, our definitions of site-wise precision and sensitivity give the same value.

#### Mosaic approach

This is the same approach used by the authors of SCUEAL (Kosakovsky Pond *et al.*, 2009). Any partition of the query into strains is translated into the sequence of strains that appear in it, ignoring the position of the breakpoints and of unassigned regions, when these are present. We call such sequence of strains a *mosaic*. For example the mosaic of the query in Fig. 1 is A1, B. The mosaic of each query is compared to the mosaic reconstructed by the software on that query. Each of these reconstructed mosaics is then classified into one of the following four categories, where the word *subsequence* is defined in the standard way, not implying contiguity (Wikipedia contributors, 2019; Gusfield, 1997). **Match**: the mosaic returned by the software coincides with the correct mosaic. **Superset:** the correct mosaic is a subsequence of the mosaic returned by the software. **Subset:** the mosaic returned by the software is a subsequence of the correct mosaic. **Mismatch:** none of the above. For example, the second mosaic in Figure 1 (returned by SHERPAS) is a superset compared to the correct mosaic (note the presence of the light brown bar towards the right), whereas the third (returned by jpHMM) is a match. For circular queries, the definitions above are modified accordingly.

### 3.3 HIV-pol dataset

To evaluate the performances of SCUEAL, Kosakovsky Pond *et al.* (2009) generated 10,000 synthetic recombinant queries, combining fragments from 863 pre-queries from the HIV-1 *pol* gene. We used this dataset without modification. The queries are about 1.6 kbp long.

To run SHERPAS on these queries, we built the pkDB using the same reference alignment as SCUEAL. This alignment contains 167 HIV *pol* sequences distributed into 17 strains, which correspond to groups, types, subtypes, chimpanzee SIV sequences, and the circulating recombinant form CRF01_AE. These strains are named A, A1, A2, A3, AE, B, C, D, F1, F2, G, H, J, K, N, O, CPZ. (The inclusion of CPZ and AE is discussed in the Suppl. Materials, Secs. 3.3 and 3.4, respectively.)

The output of SCUEAL on these queries is distributed along with the software, so we did not re-run SCUEAL on this dataset. (Also because SCUEAL is a non-deterministic algorithm.) The queries include intra-strain recombinants and SCUEAL’s output includes the detection of intra-strain recombination. In order to make this information comparable to the output of SHERPAS, we ignored intrastrain recombination, and only retained inter-strain recombination information. As a consequence, the mosaic-based accuracy measures that we obtain for SCUEAL (Table 2) are much better than those reported by Kosakovsky Pond *et al.* (2009) (e.g. 93.2% matches vs. 46.6%). In order to interpret the results for jpHMM on this dataset, we note that strains A and N cannot be recognized by jpHMM, which negatively impacts its accuracy measures on this dataset. The impact, however, is limited. (See the Suppl. Materials, Sec. 5.2 for more detail.)

### 3.4 HBV-genome dataset

Both SHERPAS and the latest version of jpHMM are able to analyze data from viruses with circular genomes, such as the hepatitis B virus (HBV) (Schultz *et al.*, 2012). To experiment with HBV data, we used the reference alignment that is distributed with jpHMM. It contains 339 whole-genome sequences classified into strains A, B, C, D, E, F, G, H (known as *genotypes*). Prior to the construction of the pkDB for SHERPAS, we extended this reference alignment by copying the first 9 columns of the alignment to the end of the alignment. This allows the construction of phylo-*k*-mers (with *k* = 10) from positions that overlap with the artificial end of the alignment.

To build a collection of queries, we started with a collection of pre-queries extracted from the database of aligned whole-genome HBV sequences available at the HBVdb website (HBVdb contributors, 2019; Hayer *et al.*, 2013). To construct a query, *2X* recombination breakpoints are chosen at random, where *X* ≥ 1 is geometrically distributed with parameter 0.8, while making sure that no two breakpoints are less than 100 bp apart (as in Kosakovsky Pond *et al.* (2009)). 2000 queries combine fragments from two pre-queries, and 1000 queries are based on three pre-queries. (See the Suppl. Materials, Sec. 5.3, for full details on this procedure.) The parameters used in this procedure were chosen so that the queries loosely reflect the characteristics of inter-genotype HBV recombinants presented in a recent overview (Araujo, 2015). The queries are about 3.2 kbp long.

### 3.5 HIV-genome dataset

This dataset consist of whole-genome sequences from HIV. Again, we used the reference alignment of jpHMM for HIV to build the pkDB database for SHERPAS. This alignment contains 881 whole-genome sequences, classified in the following 14 strains: A1, A2, AE, B, C, D, F1, F2, G, H, J, K, O, CPZ.

To construct a collection of 3000 synthetic queries, we used pre-queries extracted from Los Alamos HIV sequence database (the “complete Web alignment 2018”). In brief, the main difference with the procedure for the HBV-genome queries is that the number of parental pre-queries and the number of breakpoints are both drawn from (shifted) geometric distributions. Again, the construction procedure was designed to reflect the broad characteristics of known recombinant forms, those listed in the Los Alamos HIV sequence database. Full details of this procedure are described in the Suppl. Materials, Sec. 5.4. The average length of the resulting queries is 8.9 kbp.

### 3.6 Simulated Nanopore reads from the HIV-genome dataset

To test the robustness of SHERPAS to high error rates typical of long read sequencing technologies, we also built a dataset of reads generated with NanoSim-H, a simulator of Oxford Nanopore reads (Yang *et al.*, 2017; Břinda *et al.*, 2018). For each query in the HIV-genome dataset, we generated a single simulated read using NanoSim-H with minimum and maximum length set to 1000 and 9000, respectively, and rate of unaligned reads set to 0. All other parameters were left to their default values. A total of 3000 simulated reads, with average length about 5.9 kbp, were thus obtained. The reference alignment used for this dataset is the same as that for the HIV-genome dataset. (See the Suppl. Materials, Sec. 5.5 for details.)

### 3.7 Running the experiments

For each of the datasets described in Sections 3.3 to 3.6, a reference tree was constructed from the reference alignment with PhyML 3.3 (Guindon *et al.*, 2010) using GTR + Γ + I as substitution model. Alignment and tree were given as inputs to a customized version of RAPPAS that built a pkDB using parameters *k* = 10 and threshold parameter 1.5 (called “omega”).

We ran SHERPAS with 8 parameters combinations: SHERPAS-reduced for *θ_R_* ∈ {0.90,0.99} and window size in {300, 500}, and SHERPAS-full for *θ_F_* ∈ {1, 100} and window size again in {300, 500}. Using two values for each parameter allows us to gauge their impact on the accuracy of SHERPAS. We also ran jpHMM using its default behavior for HIV and HBV, with and without the option-Q blat to speed-up its execution. See Sec. 3.1 for motivation regarding the choice of jpHMM for comparisons. For the HIV-pol dataset (Sec. 3.3) the results of running SCUEAL are distributed together with the software (Kosakovsky Pond *et al.*, 2009), so we included them in our comparisons.

The commands used for all these operations and links to files used —including the pkDBs constructed by RAPPAS— are reported for reproducibility in the Suppl. Materials (Sec. 5). All experiments were run on the same PC with 32GB RAM and using a single core operating at 3.6GHz. Running times were measured using the Unix command time (recording user CPU time).

## 4 Results

### 4.1 Running times

Table 1 shows the running times of SHERPAS-full, SHERPAS-reduced and of two ways of executing jpHMM, that is, with and without the -Q blat option to speed-up its execution. We do not include the time necessary to construct the pkDBs with RAPPAS, as we assume that the pkDB has been obtained prior to the analysis. (To this end, SHERPAS is distributed with the 3 pkDBs used in the experiments reported here.) Moreover, the numerical parameters of SHERPAS (the *θ* thresholds and the window size) have very little impact on its running time. For this reason, we only report runtimes for default parameters. The running times for jpHMM could not be obtained in two cases for the following reasons: (1) for the HBV-genome dataset, we must run jpHMM with the -C option for circular queries, which automatically activates the -Q blat option; (2) for the simulated Nanopore HIV reads, the -Q blat option resulted in the program failing to execute, probably because of the difficulty of aligning error-rich reads.

**Table 1.** Running times of jpHMM and SHERPAS on the four datasets. Column “Mbp” reports the total size of the query dataset in Mbp. Column “#br.” reports the number of branches for which the full pkDB (reduced pkDB) stores information. “R” and “F” distinguish between SHERPAS-reduced and SHERPAS-full, respectively. “HBV-g” and “HIV-g” refer to the HBV-genome and HIV-genome datasets, respectively. “HIV-LR” refers to the dataset of simulated long reads. All times are measured in minutes (m) and seconds (s).

|  | Mbp | #br. | jpHMM |  | SHERPAS |  |
| --- | --- | --- | --- | --- | --- | --- |
|  |  |  | default | -Q blat | F | R |
| HIV-pol | 16.2 | 332(23) | 12964m 46s | 1533m 22s | 2m 40s | 32s |
| HBV-g | 9.6 | 676(8) | - | 673m 24s | 2m 35s | 11s |
| HIV-g | 26.7 | 1760(20) | 4997m 48s | 2367m 36s | 20m 44s | 51s |
| HIV-LR | 17.7 | 1760(20) | 7414m 17s | - | 12m 29s | 33s |

SHERPAS is orders of magnitude faster than jpHMM. Compared to jpHMM with the -Q blat option, SHERPAS-full is hundreds of times faster, while SHERPAS-reduced is thousands of times faster. Datasets that took days for jpHMM -Q blat to analyse, can be analyzed by SHERPAS in a matter of minutes, or even seconds.

The running time of SHERPAS essentially depends on two characteristics of the dataset. First, it scales linearly with the amount of data to analyse (number of queries and their lengths). Second, it is also related to the number of branches for which some information is stored in the pkDB. In the full version, this number is proportional to the size of the reference tree, while in the reduced version it is equal to the number of root branches. These numbers are reported in the first two columns of Table 1. See the Suppl. Materials (Sec. 2.6) for a detailed complexity analysis of the algorithms implemented in SHERPAS.

Consistent with the expectations above, the speed-up obtained with SHERPAS-reduced relative to SHERPAS-full is related to the strength of the reduction in the number of branches in the pkDB: the speed-up is moderate for HIV-pol (from 332 to 23 branches), but much more pronounced for HBV-genome (from 676 to 8 branches), and for the two whole-genome HIV datasets (from 1760 to 20 branches). As for the differences across different datasets, it is not surprising that the dataset that results in the longest running time for SHERPAS is HIV-genome: its set of queries has the largest aggregate size, and the number of branches in the full pkDB is by far the largest. Running times for HIV-LR (the simulated Nanopore reads dataset) are lower than those for HIV-genome because the simulated reads are in general shorter than the whole genome.

### 4.2 HIV-pol dataset

Table 2 compares the accuracy of inter-strain recombination detection methods (see Sec. 3.7) on the HIV-pol dataset. SCUEAL and jpHMM achieve high accuracies overall on this dataset. Here, SCUEAL and jpHMM use different reference alignments, and two strains (A and N) present in some of the queries cannot be recognized by jpHMM (see Sec. 3.3). We also observed that many of the pre-queries that Kosakovsky Pond *et al.* (2009) used to construct the queries in this dataset are in fact part of the reference alignment for HIV used by jpHMM. For these reasons, it is not a good idea to draw conclusions about the relative performance of SCUEAL and jpHMM here.

**Table 2.** Accuracies observed on the HIV-pol dataset. jpHMM-Qb stands for jpHMM with the -Q blat (fast) option. “R” and “F” distinguish between SHERPAS-reduced and SHERPAS-full, respectively. Columns “thr” and “w” report the threshold and window-size used by SHERPAS. Column “N/A” reports the percentage of sites that are not assigned to any strain. Columns “sens” and “prec” report site-wise sensitivity and precision (in percentage), respectively. Columns “m”, “sup”, “sub” and “mm” report the percentages of mosaic matches, supersets, subsets and mismatches, respectively. (See Sec. 3.2 for definitions.)

| Method | thr | w | site-wise |  |  | mosaic |  |  |  |
| --- | --- | --- | --- | --- | --- | --- | --- | --- | --- |
|  |  |  | N/A | sens | prec | m | sup | sub | mm |
| SCUEAL | - | - | 0.0 | 98.5 | 98.5 | 93.2 | 3.0 | 1.9 | 1.9 |
| jpHMM | - | - | 0.0 | 97.4 | 97.4 | 90.0 | 0.0 | 7.0 | 2.9 |
| jpHMM-Qb | - | - | 0.0 | 97.4 | 97.5 | 90.2 | 0 | 7.0 | 2.8 |
| SHERPAS R | 0.9 | 500 | 7.5 | 89.8 | 97.1 | 83.6 | 8.5 | 6.3 | 1.6 |
| SHERPAS R | 0.9 | 300 | 8.8 | 89.4 | 98.0 | 81.9 | 12.6 | 4.0 | 1.5 |
| SHERPAS R | 0.99 | 500 | 17.0 | 81.2 | 97.9 | 82.6 | 3.0 | 13.1 | 1.3 |
| SHERPAS R | 0.99 | 300 | 21.0 | 78.2 | 98.8 | 82.3 | 3.6 | 12.9 | 1.2 |
| SHERPAS F | 1 | 500 | 4.4 | 93.5 | 97.8 | 81.9 | 12.0 | 5.3 | 0.8 |
| SHERPAS F | 1 | 300 | 3.0 | 95.3 | 98.2 | 78.2 | 19.0 | 2.2 | 0.7 |
| SHERPAS F | 100 | 500 | 7.0 | 91.6 | 98.6 | 89.0 | 3.3 | 7.3 | 0.3 |
| SHERPAS F | 100 | 300 | 5.3 | 93.7 | 98.9 | 88.4 | 7.4 | 3.7 | 0.5 |

Overall, the accuracies displayed by SHERPAS on this dataset are not as good as those of the other methods, especially in terms of sitewise sensitivity and mosaic measures. The low sensitivity is due to the high incidence of unassigned regions, which is particularly pronounced for SHERPAS-reduced and high values of the thresholds. On the other hand, for SHERPAS-full, a high value of the threshold (*θ_F_* = 100) results in a better site-wise precision than SCUEAL and jpHMM, and in mosaic measures that are almost as good as those of SCUEAL and jpHMM (frequency of mosaic matches: 88.4%-89% vs. 90%-93.2%).

We also observe that on this dataset SHERPAS-full is generally more accurate than SHERPAS-reduced. This is not surprising, as SHERPAS-reduced uses far less pre-computed information (a much smaller pkDB) than SHERPAS-full. As for the effect of window size, smaller windows consistently result in higher site-wise precision, and lower frequencies of mosaic matches. This appears to be due to the fact that a smaller window “switches” more easily between different strains and therefore has a tendency to produce finer classifications, but more fragmented mosaics. This is corroborated by the observation that the frequency of superset mosaics is consistently higher for windows of size 300 than for windows of size 500.

#### 4.3 HBV-genome dataset

The results in Table 3 show that, again, jpHMM has a very high overall accuracy, which is rarely matched by SHERPAS. Some of the observations made for HIV-pol can be re-iterated here: again, the site-wise sensitivity of SHERPAS is markedly lower than that of jpHMM, and again, as expected, increasing the thresholds deteriorates sensitivity, and improves mosaic accuracy.

**Table 3.**
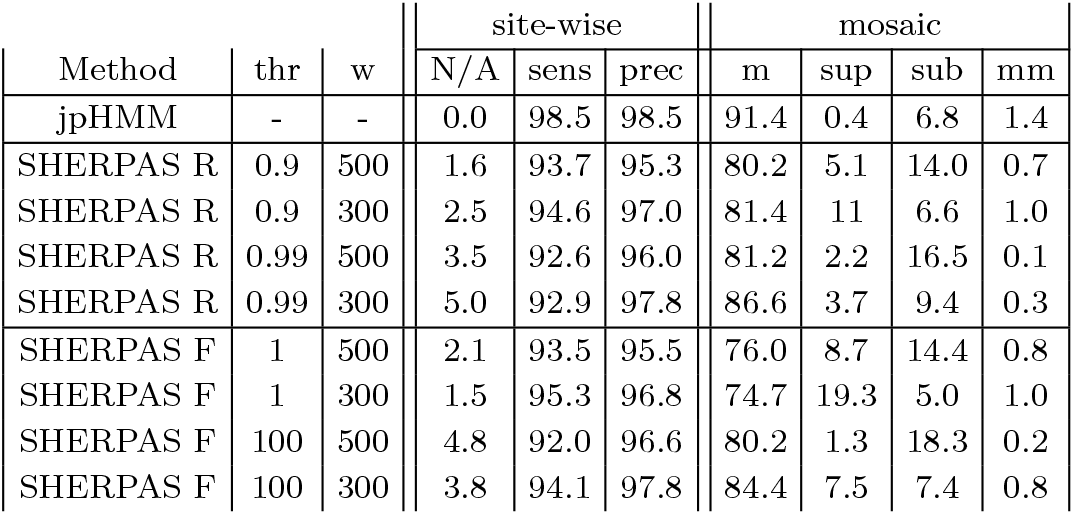
Accuracies observed on the HBV-genome dataset. jpHMM stands for jpHMM launched with the -C option for circular queries. Note that this option automatically activates the -Q blat (fast) option. All other abbreviations are as in Table 2.

Interestingly, on this dataset there does not seem to be any consistent difference between the accuracies of SHERPAS-full and SHERPAS-reduced. This may have something to do with the nature of the reference tree for HBV, where the 8 strains are monophyletic and well-delimited by relatively long root branches. (Which is not the case for all the strains in HIV-1.) This may imply that for HBV, the phylo-*k*-mers inferred for the root branches represent well their respective strains. It is also interesting to note that on this dataset, setting the window size to 300 usually leads to better results than 500, an observation that is not generally true for the other datasets.

#### 4.4 HIV-genome dataset

The results for the HIV-genome dataset, shown in Table 4, show a slightly different pattern from the other datasets. On the one hand, jpHMM and SHERPAS-full have similar site-wise accuracy measures. Unlike in the previous datasets, the sensitivity of SHERPAS-full is higher than that of jpHMM in 3 cases out of 4. On the other hand, the frequency of mosaic matches for SHERPAS is now substantially lower than that of jpHMM.

**Table 4.** Accuracies observed on the HIV-genome dataset. All abbreviations are as in Table 2

| Method | thr | w | site-wise |  |  | mosaic |  |  |  |
| --- | --- | --- | --- | --- | --- | --- | --- | --- | --- |
|  |  |  | N/A | sens | prec | m | sup | sub | mm |
| jpHMM | - | - | 3.6 | 95.6 | 99.2 | 77.8 | 4.2 | 16.6 | 1.4 |
| jpHMM-Qb | - | - | 4.0 | 95.4 | 99.3 | 78.2 | 4.0 | 17.0 | 0.8 |
| SHERPAS R | 0.9 | 500 | 3.4 | 94.5 | 97.9 | 48.2 | 37.3 | 6.1 | 8.4 |
| SHERPAS R | 0.9 | 300 | 5.5 | 92.5 | 97.9 | 27.4 | 63.0 | 1.9 | 7.7 |
| SHERPAS R | 0.99 | 500 | 6.0 | 92.7 | 98.6 | 65.8 | 15.3 | 13.4 | 5.5 |
| SHERPAS R | 0.99 | 300 | 8.6 | 90.5 | 99.0 | 56.7 | 27.0 | 9.8 | 6.5 |
| SHERPAS F | 1 | 500 | 2.1 | 96.1 | 98.2 | 46.3 | 43.6 | 4.3 | 5.8 |
| SHERPAS F | 1 | 300 | 1.6 | 96.6 | 98.2 | 24.2 | 71.5 | 0.7 | 3.6 |
| SHERPAS F | 100 | 500 | 3.5 | 95.3 | 98.8 | 67.8 | 19.2 | 9.5 | 3.5 |
| SHERPAS F | 100 | 300 | 2.7 | 96.4 | 99.1 | 54.3 | 39.6 | 2.9 | 3.2 |

These seemingly contradictory observations can be explained by inspecting the outputs of SHERPAS and jpHMM on the queries in this dataset. The Suppl. Materials (Annex C) contain an illustration of the outputs of SHERPAS-full and jpHMM on the first 100 queries out of the 3000 in this dataset. An important observation is that the partition produced by SHERPAS often includes short erroneous fragments (that is, that were not present in the correct partition of the query). For example, among the first 10 queries shown in the Suppl. Materials, 5 queries present such short erroneous fragments (queries 2, 3, 4, 6, 9; in some cases the erroneous fragment is so short that it is difficult to observe without zooming). The output of SHERPAS in Fig. 1 (corresponding to query 56) is also an example of this phenomenon: note the short erroneous fragment from strain D.

A consequence of this behavior of SHERPAS is that, although its output is usually close to the correct partition, the mosaics it produces are often supersets of the correct mosaics. This phenomenon was also present in the other datasets, as can be seen in the frequencies of supersets, which are always higher than in the other methods (see again Tables 2 and 3). However, here this becomes more visible because the queries are about 3 to 5 times longer than in the other datasets, meaning that the probability of observing such erroneous short fragments in one query increases significantly. As we discuss in Sec. 5.1, when using SHERPAS to screen for recombinants, supersets should be regarded as far less serious errors than subsets or mismatches. From Table 4, it is easy to check that here the aggregate frequency of subsets and mismatches is higher for jpHMM (about 18%) than in all 4 runs of SHERPAS-full.

The lower sensitivity of jpHMM relatively to the other datasets is due to its behavior at the two ends of queries spanning a whole HIV genome. As can be seen in the Suppl. Materials (Annex C), jpHMM often leaves the ends of a HIV-genome query as unassigned (N/A). This is likely due to the difficulty of alignment and of profile-based modeling in those peripheral regions.

Like in the HIV-pol dataset, we note that SHERPAS-full tends to be slightly more accurate than SHERPAS-reduced, in terms of sitewise measures. Using a window of size 300 instead of 500, strongly reduces the frequency of mosaic matches, which is again due to a higher frequency of short erroneous fragments. However, it consistently reduces the aggregate frequency of subsets and mismatches (not shown) which may be important for screening purposes (Sec. 5.1).

#### 4.5 Simulated Nanopore reads from the HIV-genome dataset

The results in Table 5 show that the simulated Nanopore reads pose a significant challenge to jpHMM and SHERPAS. This is not surprising, given the high error rates that characterize these reads. We refer to Yang *et al.* (2017) and its Suppl. Materials for an in-depth analysis of these error rates.

**Table 5.**
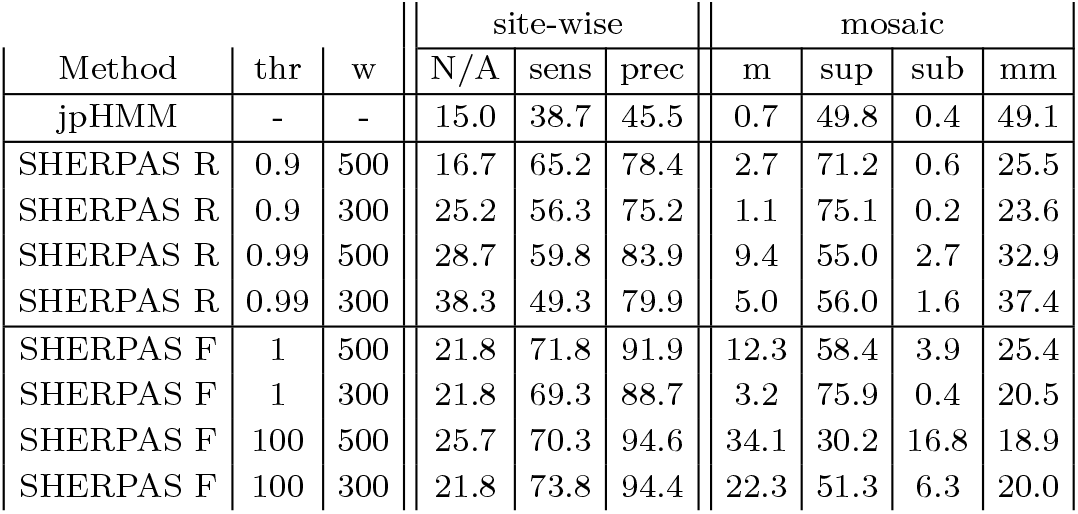
Accuracies observed on the dataset of simulated Nanopore HIV reads. For jpHMM only the results of launching it with its default options for HIV are reported, as the use of the -Q blat (fast) option resulted in the program failing to execute. All abbreviations are as in Table 2.

| Method | thr | w | site-wise |  |  | mosaic |  |  |  |
| --- | --- | --- | --- | --- | --- | --- | --- | --- | --- |
|  |  |  | N/A | sens | prec | m | sup | sub | mm |
| jpHMM | - | - | 15.0 | 38.7 | 45.5 | 0.7 | 49.8 | 0.4 | 49.1 |
| SHERPAS R | 0.9 | 500 | 16.7 | 65.2 | 78.4 | 2.7 | 71.2 | 0.6 | 25.5 |
| SHERPAS R | 0.9 | 300 | 25.2 | 56.3 | 75.2 | 1.1 | 75.1 | 0.2 | 23.6 |
| SHERPAS R | 0.99 | 500 | 28.7 | 59.8 | 83.9 | 9.4 | 55.0 | 2.7 | 32.9 |
| SHERPAS R | 0.99 | 300 | 38.3 | 49.3 | 79.9 | 5.0 | 56.0 | 1.6 | 37.4 |
| SHERPAS F | 1 | 500 | 21.8 | 71.8 | 91.9 | 12.3 | 58.4 | 3.9 | 25.4 |
| SHERPAS F | 1 | 300 | 21.8 | 69.3 | 88.7 | 3.2 | 75.9 | 0.4 | 20.5 |
| SHERPAS F | 100 | 500 | 25.7 | 70.3 | 94.6 | 34.1 | 30.2 | 16.8 | 18.9 |
| SHERPAS F | 100 | 300 | 21.8 | 73.8 | 94.4 | 22.3 | 51.3 | 6.3 | 20.0 |

Strikingly, however, jpHMM is much more negatively affected by the simulated Nanopore sequencing errors than SHERPAS. Note that if a classifier was to choose randomly one of the 14 strains at every site, its sensitivity and precision would be 1/14 = 7.1%, but if it was to classify every query as a non-recombinant sequence belonging to the most frequent strain (B) its sensitivity and precision would be 41.9% (this is the proportion of sites from strain B in the queries). Thus, jpHMM’s site-wise accuracy measures are only partially better than those of random classifiers. On the other hand the site-wise precision of SHERPAS, especially in the full version, is only marginally affected (cf. Table 4). The site-wise sensitivity is lowered, which is due to the fact that error-rich regions are often unassigned. SHERPAS is also more accurate than jpHMM in terms of mosaic measures (cf. the frequencies of matches and mismatches).

Finally, once again SHERPAS-full is consistently more accurate than SHERPAS-reduced. Interestingly, on this dataset, using a smaller window generally results in a deterioration of accuracy. However, this is not true if the goal is to minimize the aggregate frequency of subsets and mismatches (see Sec. 5.1).

## 5 Discussion

### 5.1 Uses of SHERPAS

SHERPAS is a tool for the detection and analysis of inter-strain recombinants in a large collection of query sequences. It relies on the availability of a reference multiple sequence alignment, which is used to “learn” to recognize sequences from the different strains. It accomplishes a bioinformatics task considerably different from that of detecting the presence of recombinant sequences within a multiple sequence alignment — a task that can be tackled with other methods, such as those implemented in the RDP software (Martin *et al.*, 2015, 2017). An important difference between the two tasks is that here we make a clear distinction between reference sequences (known in advance and well-characterized) and the novel sequences to analyse, the queries. The latter do not need to be aligned, which opens the possibility of treating a much larger amount of sequence data. SHERPAS can be used for any dataset of viral sequences for which a reference alignment of sufficient quality and size can be obtained. In fact SHERPAS, like SCUEAL, could also be used to detect recombinant bacterial sequences (Kosakovsky Pond *et al.*, 2009), although we have not experimented with such data.

By default, SHERPAS uses the full pkDB, with *θ_F_* = 100 and window size 300. If the user chooses to run SHERPAS-reduced, the default parameters are *θ_R_* = 0.99 and again window size 300. These default settings were chosen while trying to achieve a good balance among all accuracy measures, and assuming a volume of data that is not prohibitively large. However, users should be aware that the choice of settings will depend on the nature of the data and the goal of the analysis.

For example, SHERPAS may be used as a first screen to detect potential recombinants in a large set of sequences. The putative recombinants can then be analysed further with more accurate but slower software, such as REGA (de Oliveira *et al.*, 2005; Pineda-Peña *et al.*, 2013), SCUEAL (Kosakovsky Pond *et al.*, 2009) or jpHMM (Schultz *et al.*, 2006, 2009). In this case, the primary goal is not high accuracy, but rather to lower the odds of missing evidence of recombination in a query. In terms of inferred mosaics, this means lowering the frequencies of subsets and mismatches. In Tables 2 – 5, the parameter combinations that minimize the occurrence of subsets and mismatches for SHERPAS-full and SHERPAS-reduced are the ones with the lowest tested thresholds and window size 300. Note that these combinations consistently produce fewer aggregate subsets and mismatches than jpHMM.

To provide further insight into the ability of detecting evidence of recombination, we re-analyzed the queries in the HIV-pol dataset, which include a substantial number of sequences that are not interstrain recombinants. We then considered SHERPAS as a binary classifier for inter-strain recombination, classifying a sequence as a “positive” if at least one breakpoint is detected, and a “negative” otherwise. This allows us to observe how changing settings in SHERPAS (full vs. reduced database, threshold and window size) affects its ability of recovering true positives (known as the *recall* of the classifier), and how much specificity has to be traded to improve this ability. The results of this experiment, shown in Fig. 2, confirm that SHERPAS can indeed achieve high recall with the use of low thresholds (not the default one), and small windows, although users must be aware that this entails a loss of specificity. A full description and discussion about this experiment can be found in the Suppl. Materials (Sec. 5.6).

**Fig. 2.**
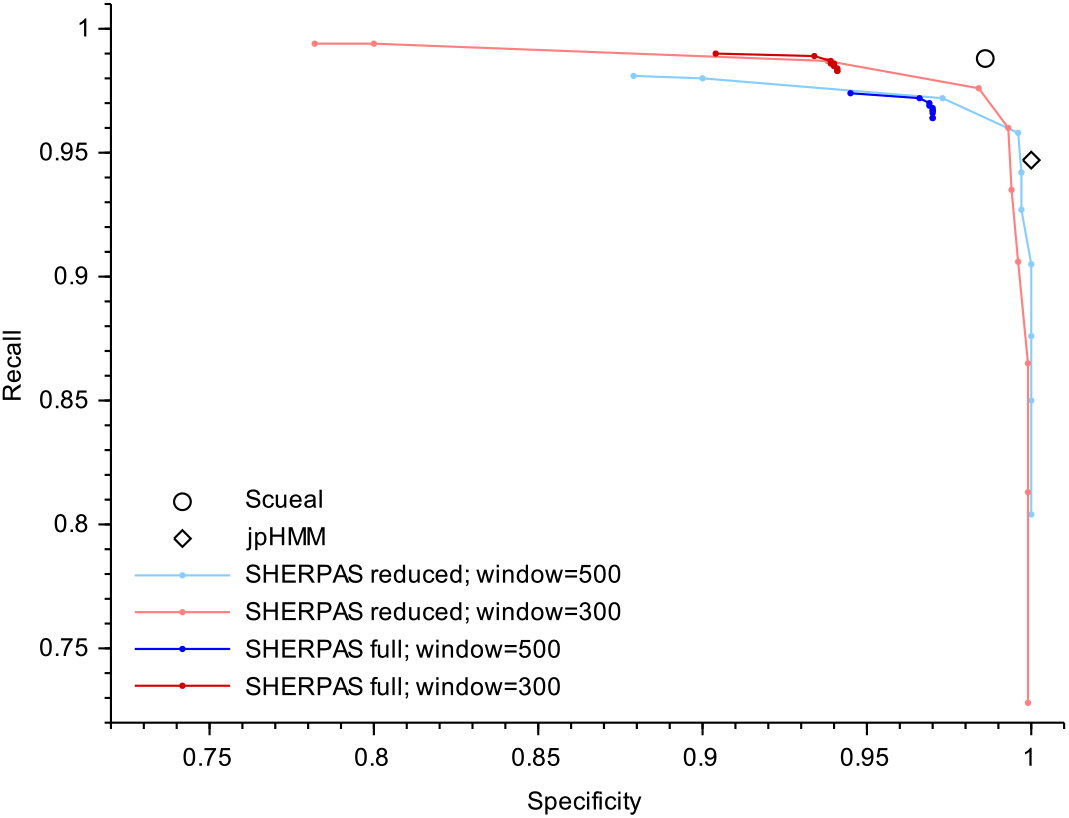
Trade-off between recall and specificity for the binary classification of HIV-pol queries. Recall and specificity are plotted for SCUEAL (circle), jpHMM (diamond) and SHERPAS (colored lines). The four colored lines correspond to the different combinations of a pkDB version (full/reduced) and window size (500,300) for SHERPAS. Each point in a colored line corresponds to a different value of the threshold, with the lowest values of the threshold (1 for SHERPAS-full and 0 for SHERPAS-reduced) resulting in the leftmost points. See the Suppl. Materials (Sec. 5.6) for full details. Note that all rates fall in the interval [0-72,1], which is why the curves are not depicted in the full [0,1] range.

### 5.2 Scaling-up

Another important factor influencing how SHERPAS should be run is the amount of data to analyse. The query datasets that we used here were of relatively manageable sizes, to facilitate comparisons with slower software. Should the data to analyse be substantially more abundant (e.g. millions of reads), running SHERPAS in reduced mode may become more appealing, or even necessary in some cases. This is especially true if the reference tree is large, as in this case the speedup for SHERPAS-reduced is more pronounced (see Sec. 4.1). Moreover, for some datasets or applications, the accuracy of SHERPAS-reduced may be comparable to that of SHERPAS-full (see, e.g., Sec. 4.3 and Fig. 2).

Because the running times of SHERPAS scale linearly with the amount of data to analyse, running SHERPAS on few millions queries is feasible in a matter of days (using SHERPAS-full) or in a matter of hours (using SHERPAS-reduced; see Table 1). To the best of our knowledge, none of the recombination detection tools currently available are scalable to datasets of that size. Although our experiments focused on comparing SHERPAS to jpHMM —for the reasons detailed in Sec. 3.1— previous comparisons of running times between jpHMM and phylogeny-based tools for recombination detection (namely REGA and SCUEAL) showed that jpHMM was at least as fast as those tools (see Pineda-Peña *et al.* (2013), Table 4), when run with the fast -Q blat option, meaning that the running time advantage of SHERPAS likely extends to the other available tools.

### 5.3 Accuracy

Consistent with previous literature (Schultz *et al.*, 2006; Kosakovsky Pond *et al.*, 2009; Schultz *et al.*, 2009), we evaluated the predictive accuracy of SHERPAS using large datasets of semi-artificial recombinant sequences, which combine fragments of real HIV and HBV sequences (the *pre-queries*) via artificially-introduced breakpoints. In the absence of large datasets of real sequences for which the true recombinant structure is known with certainty, this is a good way to evaluate a new method for recombination detection. (Note that we ensure that none of the pre-queries belongs to the reference alignment for the tested methods.) Using sequences that are fully simulated using a fixed evolutionary model (Kosakovsky Pond *et al.*, 2009) is also a viable option, but the choice of the simulation parameters can have an important impact on the results, and the advantage over using semi-artificial sequences is unclear.

The experiments were designed to assess accuracy loss in SHERPAS compared to jpHMM (Schultz *et al.*, 2006, 2009). The choice of jpHMM is motivated in Sec. 3.1. The other goal of our experiments was to explore the influence of the parameters of SHERPAS, including the use of full/reduced pkDB. Using two possible values for all parameters allows us to gauge their impact. The two window sizes (300 and 500 bp) were chosen on the basis of common practices of sliding window approaches (e.g. the REGA tool employs a window of 400 bp).

Possibly the most interesting result here is the inferior performance of jpHMM compared to SHERPAS on the simulated Nanopore reads dataset. We suspect that the reason for this is that profile HMMs may be strongly affected by large indels (which are common in these reads) and by errors that do not correspond well to their emission probabilities (which were estimated on error-free datasets by Schultz *et al.* (2006)). SHERPAS, on the other hand, appears to be able to exploit the information coming from the error-free stretches of sequences that lie between errors in the reads. Further work, beyond the scope of the present paper, would be needed to investigate these hypotheses.

### 5.4 Future work and limitations

SHERPAS-full implicitly computes the most probable branch of origin of any window within a query. This means that it can be used for precise phylogenetic placement of the segments composing the query (Matsen *et al.*, 2010; Berger *et al.*, 2011; Barbera *et al.*, 2019; Linard *et al.*, 2019), or even to detect *intra*-strain recombinants. We plan to add these functionalities in future versions of SHERPAS.

Second, the use of a sliding window has a few well-known disadvantages (Kosakovsky Pond *et al.*, 2009). Specifically, it makes it hard to precisely locate breakpoints (Schultz *et al.*, 2006), and choosing its size involves a trade-off between resolution and informativeness. In the future, we plan to implement algorithms that are not window-based in SHERPAS (using, e.g., dynamic programming). However, this will potentially entail a cost in terms of computational efficiency.

Third, here we focused on the problem of recognizing cases of *homologous* recombination, which occurs when the new sequence combines parental fragments with different origins, but joined at homologous sites. Non-homologous or *illegitimate* recombination is also known to occur in viruses, and results in genomes displaying structural changes (e.g. with large insertions, deletions, duplications etc.) (Crawford-Miksza and Schnurr, 1996; Scheel *et al.*, 2013; Galli and Bukh, 2014). Some preliminary experiments (not shown) suggest that SHERPAS is also able to recognize and correctly partition non-homologous recombinants. Note that phylogeny-based tools such as SCUEAL and REGA align the query to the reference sequences prior to the analysis, a problematic step when the query contains, for example, genomic duplications or translocations. In the future, we plan to conduct an in-depth study of this novel functionality of SHERPAS.

Fourth, SHERPAS was developed to detect novel recombinants, but not to recognize widespread and well-known recombinants — known as *circulating recombinant forms* (CRFs). If some of the query sequences are CRFs, SHERPAS should detect that they are interstrain recombinants, and partition them accordingly. Although we have done so for one CRF (CRF01_AE) in HIV-1, including CRFs in the reference alignment and defining one strain per CRF is risky, as the reference tree will not be an accurate description of the true history. This point is discussed in depth in the Suppl. Materials (Sections 3.2 and 3.3), where we also explain how users may solve this problem by modifying the reference alignment following an idea already exploited for example, by Kosakovsky Pond *et al.* (2009) and D. Martin (personal communication). Automatic treatment of CRFs is a possible extension that we plan to add to SHERPAS.

Finally, every step involved in SHERPAS’s analyses can be in principle parallelized, including the construction of the phylo-*k*-mer database. This would further improve the scalability of our approach.

### 5.5 Conclusion

SHERPAS achieves a reasonable accuracy compared to state-of-the-art inter-strain recombination detection tools for viruses, but is orders of magnitude more efficient. This advantage derives from the fact that SHERPAS does not need to align the query sequences, and from the relative simplicity of its classification algorithm. To the best of our knowledge, it is the first software that can estimate the recombinant structure of thousands of long sequences (up to whole genomes) within minutes or even seconds. It also appears to be relatively robust to high error rates typical of long read sequencing technologies. SHERPAS paves the way to systematic screening of recombinants in large datasets of long reads or assembled genome sequences.

## Acknowledgements

We thank Philippe Roumagnac for useful discussions and guidance, and the VIROGENESIS consortium (http://www.virogenesis.eu/) for stimulating this work — in particular Pieter Libin, Kristof Theys and Anne-Mieke Vandamme.

## Funding and conflicts of interest

This work was publicly funded through ANR (The French National Research Agency) under the *Investissements d’avenir* programme with the reference ANR-16-IDEX-0006. B.L. is employed by Spygen, a company specialising in aquatic and terrestrial biodiversity monitoring using environmental DNA. N.R. is supported by a doctoral fellowship from the French *Ministère de l’Enseignement supérieur, de la Recherche et de l’Innovation*. We also received support from *Institut Français de Bioinformatique* (ANR-11-INBS-0013).

## Notes

### Competing Interest Statement

The authors have declared no competing interest.

### Summary of Updates

Final version. Supplementary Materials are now included.

