## Supplementary Materials for "Rapid screening and detection of inter-type viral recombinants using phylo-*k*-mers"

**References** **25**

**A Annex: Marginal posterior distributions of ancestral states** **27**

**B Annex: Approximate relationship between branch scores and likelihoods** **28**

**C Annex: Some illustrated outputs** **30**

### 1 Phylo-$k$-mers, formalized

For completeness, here we describe in detail what phylo-$k$-mers are, and the main ideas behind their construction. The material presented in Section 1 not novel, as it was introduced informally in the original publication about RAPPAS [11]. The treatment here is somewhat more formal, which should help readers interested in the details of the technique, or as a first step towards understanding its theoretical foundations.

#### 1.1 The reference data and their preprocessing

The input data for the construction of phylo-$k$-mers are a *reference alignment* $A_0$, a *nucleotide substitution model* (e.g. GTR+Γ), and a rooted *reference tree* $T_0$ describing the evolutionary process that led to the sequences in $A_0$ (i.e. there is a one-to-one correspondence between the leaves of $T_0$ and the sequences aligned in $A_0$). By default, the branch lengths of $T_0$ and the parameters $\theta$ of the substitution model are re-estimated using $A_0$ at launch of the phylo-$k$-mer building process.

Both references (alignment and tree) are then preprocessed for the construction of phylo-$k$-mers. Improved ways to do this are the subject of ongoing research. Here we describe the way this is performed currently by default. We denote the set of nodes, leaves and branches (edges) of any tree $T$ by $V(T)$, $L(T)$ and $E(T)$, respectively. Note that $L(T) \subset V(T)$.

First, we remove from $A_0$ all columns that contain more than a certain fraction of gaps "–" (0.99 by default). The rationale for this step is that ancestral reconstruction techniques do not account for gaps, and highly gappy columns have been observed to be deleterious for phylo-$k$-mer construction. We call the resulting alignment the *reduced reference alignment*, and denote it by $A$.

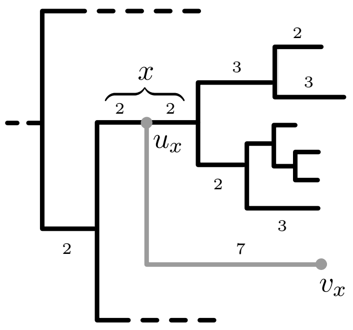

Figure 1: **Injection of two ghost nodes on a branch of the reference tree.** Depicted in black is the reference tree $T_0$. Branches are labelled by their lengths. Branches without labels have length 1. In gray, two ghost nodes $u_x, v_x$ injected over branch $x$: $u_x$ is placed on the midpoint of $x$, giving rise to two new branches of length 2; $v_x$ is a new leaf, connected to $u_x$ via the new branch $(u_x, v_x)$. Since the lengths of the paths from $u_x$ to its 6 descendants in $T_0$ are 7, 8, 6, 7, 7, 7, the length of the new branch is set to their average, i.e. 7. To get the extended reference tree, this process is repeated for each $x \in E(T_0)$.

As for the reference tree, a number of nodes and branches are added to $T_0$, leading to what we call the *extended reference tree* $T$. For each branch $x$ in $T_0$, two new nodes $u_x$ and $v_x$ are introduced, as shown in Figure 1: node $u_x$ is placed at the midpoint of branch $x$, and a new branch $(u_x, v_x)$ leading to a new leaf $v_x$ is also added. The length of this new branch is set to the average length among all paths from $u_x$ to the leaves in $L(T_0)$ that descend from $u_x$. This length is the only aspect that depends on the position of the root of $T_0$ in the phylo-$k$-mer building process. Note that $V(T_0) \subset V(T)$. Nodes that are in $V(T)$ but not $V(T_0)$ are called *ghost nodes*.

#### 1.2 Probability of a $k$-mer at a position in the reference tree and alignment

We consider the reduced reference alignment as a matrix $A = (a_{u,i})$, where $a_{u,i} \in \{\mathtt{A}, \mathtt{C}, \mathtt{G}, \mathtt{T}, -\}$ is the nucleotide or gap in the sequence at leaf $u \in L(T_0)$ aligned to site $i \in \{1, \ldots, m\}$, where $m$ is the number of columns in $A$. Note that gaps are usually interpreted and treated as missing data in phylogenetics. Aside from gaps, the contents of $A$ represent all sequence data that have been actually observed from the reference tree.

Here, we wish to model nucleotides at the nodes in $V(T) \setminus L(T_0)$, i.e. the nodes in the extended reference tree from which no data has been observed. In order to represent these unobserved nucleotides, we define, for every $u \in V(T)$ and every site $i$, the random variable

$$A'_{u,i} = \text{nucleotide at site } i \text{ of the sequence at node } u.$$

Note that the only possible realizations of this random variable are in $\{\mathtt{A}, \mathtt{C}, \mathtt{G}, \mathtt{T}\}$ and that whenever $u \in L(T_0)$ and $a_{u,i} \neq -$, we must have $A'_{u,i} = a_{u,i}$. Thus, the $A'$ variables specify a random "extension" of alignment $A$. For the calculations that follow, the only relevant $A'_{u,i}$ variables are those where $u$ is a ghost node.

The observed data in $A$ and the substitution model specified by $\theta$ determine the marginal posterior distribution of $A'_{u,i}$, i.e. the probabilities, for every $a \in \{\mathtt{A}, \mathtt{C}, \mathtt{G}, \mathtt{T}\}$

$$\mathbb{P}[A'_{u,i} = a \mid A, \theta]. \tag{1}$$

Calculating these probabilities can be performed with standard techniques of ancestral state reconstruction, which for completeness we briefly summarize in Annex A.

Because the probabilities above depend on the particular nucleotide substitution model that the user wishes to adopt, and because substitution models are a subject of active research, their calculation is best performed by well-maintained and up-to-date software packages for likelihood-based phylogenetic calculations, such as RAxML-NG [9] and PhyML [6]. The choice of the software to use as well as the nucleotide substitution model and its parameters $\theta$, can be controlled with suitable options of the phylo-$k$-mer construction software (currently a module within RAPPAS). The result of this step is a large collection of tables containing all the marginal posterior probabilities of $A'_{u,i}$ that are relevant for the calculations that follow. See Figure 2 for an example of such a table, and its use.

Now let $S_{u,i}$ be the random sequence $A'_{u,i}A'_{u,i+1} \ldots A'_{u,i+k-1}$, that is the $k$-mer at sites $i, i + 1, \ldots, i + k - 1$ in the sequence at node $u$. Using the assumption of independent evolution at different sites, we calculate the probability distribution of $S_{u,i}$ as follows:

$$\mathbb{P}[S_{u,i} = a_1 a_2 \ldots a_k \mid A, \theta] = \prod_{j=1}^{k} \mathbb{P}[A'_{u,i+j-1} = a_j \mid A, \theta]. \tag{2}$$

The assumption of independence between sites, although unrealistic, is standard practice in phylogenetics, and is also needed for computing the probabilities in (1).

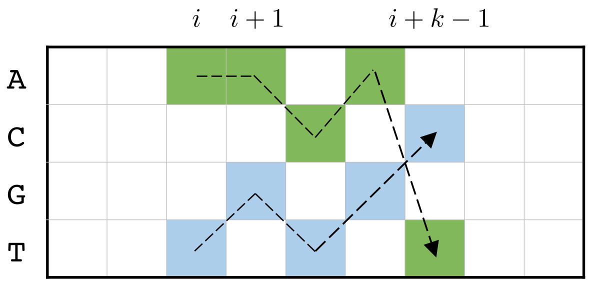

Figure 2: **Computing the probability of a $k$-mer at $k$ consecutive sites in $A$.** Software for likelihood-based phylogenetics can be used to produce, for any given ghost node $u$, a table of marginal posterior probabilities of the four nucleotides at $u$ and at all sites in $A$, i.e. the probabilities in (1). The table for a specific node $u$ is depicted here with nucleotides corresponding to rows and sites corresponding to columns. Given this table, the probability of any $k$-mer at $u$ and any $k$ consecutive sites in $A$ can be computed using (2). For example, the probability of 5-mer `AACAT` (resp. `TGTGC`) at $u$ and sites $i$ to $i+4$ is obtained by multiplying the table entries highlighted in green (resp. blue).

##### 1.3 Phylo-$k$-mers and their probability score

In the previous section, we have explained the simple ideas behind calculating the probability $\mathbb{P}[S_{u,i} = w \mid A, \theta]$ for any $k$-mer $w$, any node $u \in V(T)$, and any site $i \in \{1, \ldots, m-k+1\}$. Also recall that each branch $x \in E(T_0)$ is associated to a subset of ghost nodes in $V(T)$ (by default $\{u_x, v_x\}$, see Section 1.1). Let $G_x$ denote this subset.

For any $k$-mer $w$ and any branch of the original reference tree $x \in E(T_0)$, we then define the *probability score* of $w$ at $x$ as

$$\widetilde{\mathbb{P}}[w|x] = \max_{u \in G_x} \max_{i \in \{1,\ldots,m-k+1\}} \mathbb{P}[\, S_{u,i} = w \mid A, \theta \,]. \tag{3}$$

In other words, $\widetilde{\mathbb{P}}[w|x]$ is probability of $w$ at the best placement for $w$ within $A$ and among the nodes in $G_x$. The $\widetilde{\mathbb{P}}[w|x]$ scores play a central role to determine the phylogenetic origin of a sequence in both SHERPAS and RAPPAS [11]. Because of this, it is helpful to pause here and offer a few informal interpretations of what they mean.

1. The simplest way to interpret $\widetilde{\mathbb{P}}[w|x]$ is as an approximation of the probability of $k$-mer $w$ being present in a sequence that has diverged from the reference tree $T_0$ at branch $x$. Note, however, that if we were to compute such probability, it would make more sense to take a weighted average over $u \in G_x$ instead of the first maximum, and to take $\mathbb{P}[\,$there exists $i$ such that $S_{u,i} = w \mid A, \theta\,]$ instead of the second maximum.

2. We can also view $\widetilde{\mathbb{P}}[w|x]$ as a quantity proportional to the maximum likelihood score of the phylogenetic tree that is obtained from $T_0$ by inserting $w$ at the end of a new branch attached to $x$ (as in Figure 1). Taking the maximum over $i \in \{1, \ldots, m-k+1\}$ corresponds to finding a maximum likelihood alignment of $k$-mer $w$ to $A$, constrained to having no indel breaking up the $k$-mer. Taking the maximum over $u \in G_x$ corresponds to finding a maximum likelihood configuration within a small set of possible configurations for the branch lengths surrounding the attachment point of $w$ to $x$. (E.g. when $G_x = \{u_x, v_x\}$, as in Figure 1, placing $w$ at $u_x$ corresponds to setting the new pendant branch length to 0, while placing it at $v_x$ corresponds to setting it to 7.)

In short, all that matters for SHERPAS and other algorithms using the $\widetilde{\mathbb{P}}[w|x]$ scores is that they represent a probabilistic measure of how likely $x$ is to be the phylogenetic origin of $k$-mer $w$. Note that this implies that scores coming from different $k$-mers must be combined using products, or equivalently by using sums of their logarithms.

A *phylo-k-mer* is defined as a $k$-mer $w$ for which there exists at least one branch $x \in E(T_0)$ with $\widetilde{\mathbb{P}}[w|x]$ larger than a given threshold, currently set to $(\omega/4)^k$, where $\omega$ is a user-set parameter (1.5 by default).

The phylo-$k$-mer database construction step relies on algorithms that generate all such phylo-$k$-mers, and stores them together with a list of (1) all the branches in $E(T_0)$ for which $\widetilde{\mathbb{P}}[w|x]$ is larger than the threshold, and (2) their respective probability scores $\widetilde{\mathbb{P}}[w|x]$ (see next section for more details). A number of alternative algorithms for this step have been the subject of recent research which will be described in an upcoming publication (N. Romashchenko et al., pers. comm.).

#### 1.4 The full pkDB

The *full phylo-k-mer database* (*pkDB*) is a data structure containing all phylo-$k$-mers and their associated information. It is a look-up table $B$ that, given a phylo-$k$-mer $w$, returns the list of pairs

$$B(w) = \Big[(x_1, s_1), (x_2, s_2), \ldots, (x_{|B(w)|}, s_{|B(w)|})\Big]$$

where the $x_i$ are branches of the reference tree (hence the use of the letter $B$) and

$$s_i = \log \widetilde{\mathbb{P}}[w|x_i].$$

Unless otherwise stated, all logarithms are base 10.

The pkDB also contains all the relevant information regarding the parameters that were used to generate the phylo-$k$-mers, most notably the threshold on their scores. In the following we call $\mathsf{Thr}_B$ the threshold for log-probability scores, meaning that all $s_i$ in a $B(w)$ list are such that $s_i > \mathsf{Thr}_B$.

The value of the threshold is taken into account in the calculations performed by SHERPAS: anytime a branch $x$ does not appear in $B(w)$, SHERPAS assumes $\log \widetilde{\mathbb{P}}[w|x] = \mathsf{Thr}_B$.

##### 1.5 The reduced pkDB

The *root branches* of the reference tree are those that lie at the root of a maximal subtree whose leaves all belong to the same strain. (See the main text for an equivalent definition.)

The *reduced pkDB* $B'$ is obtained from the full pkDB $B$ by removing all pairs $(x_i, s_i)$ where $x_i$ is not a root branch. That is, if $B(w) = \left[(x_1, s_1), (x_2, s_2), \ldots, (x_{|B(w)|}, s_{|B(w)|})\right]$ and $x_{i_1}, x_{i_2}, \ldots, x_{i_l}$ are the root branches among $x_1, x_2, \ldots, x_{|B(w)|}$, then

$$B'(w) = [(x_{i_1}, s_{i_1}), (x_{i_2}, s_{i_2}), \ldots, (x_{i_l}, s_{i_l})] .$$

In the following, for simplicity we denote by $B$ the pkDB used, regardless of whether the full or reduced version is used.

#### 2 The algorithm, formalized

##### 2.1 The algorithm at a glance

Below, we provide a detailed description of the algorithm underlying SHERPAS, complementing the high-level description in the main text. In order to understand the pseudocode that follows, here we introduce the main important ideas, namely the mathematical scores that SHERPAS computes.

At any point during the execution of SHERPAS, only a subset of the branches of the reference tree are "active". These are the branches that are associated by the pkDB to at least one $k$-mer in the current window. We will denote this subset of branches by $L$, for consistency with the algorithm used by RAPPAS for placement [11].

As the sliding window moves along, SHERPAS updates $L$, and for every branch $x$ currently in $L$, it also keeps a score $S[x]$ equal to the sum of all the log-probabilities associated to $x$ by the $k$-mers in the current window. To express this mathematically, let $W$ denote the sequence of $k$-mers in the current window. That is, $W = (w_1, w_2, \ldots, w_{|W|})$, where each $w_i$ is a $k$-mer that overlaps by $k-1$ characters with $w_{i-1}$ and $w_{i+1}$, for $i \in \{2, \ldots, |W|-1\}$. Then,

$$S[x] = \sum_{i=1}^{|W|} \log \widetilde{\mathbb{P}}[w_i|x] \tag{4}$$

Recall that if $x$ does not appear in any pair in $B(w_i)$, SHERPAS assumes $\log \widetilde{\mathbb{P}}[w_i|x] = \mathsf{Thr}_B$ where $\mathsf{Thr}_B$ is the threshold for log-probability scores, used for the construction of the pkDB (see Section 1.4).

To determine the recombinant structure of the query sequence, for every window SHERPAS inspects the branch or branches in $L$ that have the highest score $S[x]$. In order to find these branches efficiently, $L$ is implemented as a binary max-heap [2], a data structure that partially orders the branches it contains on the basis of their scores $S[x]$, and that is efficient to update. Every time a branch changes score or is added or removed from $L$, the max-heap is updated using a well-known procedure that for completeness we report in Sec. 2.3.

#### 2.2 The main procedure

Let $W_0, W_1, \ldots, W_N$ denote the windows considered by SHERPAS during its execution. We set $W_0 = \emptyset$ for mathematical convenience. The way the other windows are defined depends on whether the query is circular or not. If the sequence is not circular, the first window $W_1$ contains the first 100 $k$-mers of the sequence. Then two $k$-mers are added from one window to the next, until the size of a window coincides with the standard size (300 by default). As that size is attained, at each step the first $k$-mer of the current window $W_i$ is removed, and the $k$-mer following the last $k$-mer in $W_i$ is added, to get the next window $W_{i+1}$. Finally, when the end of the query is reached, the first two $k$-mers of the current window are removed at each step, until we get a final window $W_N$ of size 100.

If the query sequence represents a full circular genome, the size of all the windows are the same, and each window is obtained from the previous one by removing the first $k$-mer and adding the next one. However, a pre-processing step modifies the query, to reflect its circularity, as follows: The last $(|W| + k - 1)/2$ base pairs of the sequence are added to its beginning, and the first $(|W| + k - 1)/2$ base pairs are added to its end. Note that $|W| + k - 1$ equals the length of the sliding window in base pairs.

The pseudocode below describes the algorithm employed by SHERPAS to treat one query. The following notation and terminology is adopted:

- The set of *candidate branches* $E$ is the set of all branches for which some information is stored in the pkDB. For SHERPAS-full, $E = E(T_0)$, i.e. $E$ is the set of all branches in the reference tree, whereas for SHERPAS-reduced, $E$ is the set of root branches.
- $+=$ denotes increment of its left-hand side by its right-hand side.
- $W_i \setminus W_{i-1}$ denotes the sequence of $k$-mers that are newly added to $W_i$. Note that, unless $i = 1$, there are at most two $k$-mers in $W_i \setminus W_{i-1}$, but usually (when $|W_i| = |W_{i-1}|$) there is exactly one $k$-mer here.
- $W_{i-1} \setminus W_i$ denotes the sequence of $k$-mers that are removed when transitioning from $W_{i-1}$ to $W_i$. Again there are at most two $k$-mers here (usually exactly one).
- $\biguplus B(w), w \in W$ denotes the concatenation of the lists $B(w)$, for all $k$-mers $w$ in $W$.
- $L$ is the data structure containing the branches that are currently active, that is, associated by the pkDB to at least one $k$-mer in the current window.

- $C[\,]$ is a table of counts, where $C[x]$ contains the number of $k$-mers in the current window that are associated by the pkDB to branch $x$.

**Algorithm 1:** SHERPAS, main algorithm (treatment of one query)

**Input:** A query with (ordered) windows $W_1, \ldots, W_N$, a pkDB $B$, a set of candidate branches $E$, a threshold $\theta$

**Output:** For each window $W_i$, a strain $P_i$ associated to $W_i$

```
set C[x] = 0 for all branches x ∈ E
set L = ∅, W_0 = ∅
for i = 1 to N do
    set B_a = ⊎ B(w_a), w_a ∈ W_i \ W_{i-1}          // B_a = branches added
    set B_r = ⊎ B(w_r), w_r ∈ W_{i-1} \ W_i          // B_r = branches removed

    for all pairs (x, s) in B_a do
        if C[x] = 0 then
            add x at the end of L
            set S[x] = |W_{i-1}| · Thr_B             //Set S[x] to its minimum possible value
        C[x] += 1
        S[x] += s − Thr_B
        Update(L, x)

    for all pairs (x, s) in B_r do
        C[x] += (−1)
        S[x] += Thr_B − s
        if C[x] = 0 then
            set S[x] = −∞                            // This causes the removal of x from L
        Update(L, x)

    if |W_i| ≠ |W_{i-1}| then
        for all branches x ∈ L do
            S[x] += Thr_B · (|W_i| − |W_{i-1}|)      // Minimum possible value correction

    set P_i = Get_strain(L, θ)
return P_1, ..., P_N
```

Sub-procedures Update and Get_strain are described in subsections 2.3 and 2.5, respectively.

#### 2.3 Heap property and update

As mentioned in Sec. 2.1, $L$ is a binary max-heap. We adopt here the standard practice to implement $L$ as an array whose first element is $L[1]$ and whose last element is $L[|L|]$ [2]. Each of its elements $L[j]$ is a branch in $E$, and we ensure that $L$ satisfies the *heap property*, that is, for all $j \in \{2, \ldots, |L|\}$, we have $S[L[\lfloor j/2 \rfloor]] \geq S[L[j]]$. We use this structure because it is computationally fast to update it when the score of an element of $L$ changes (Algorithm 2), and because it ensures that at any time the following two properties are satisfied:

(i) The best-scoring branch is $L[1]$.

(ii) The second best-scoring branch is either $L[2]$ or $L[3]$, whichever has the greatest score.

In terms of computational complexity, each update is carried out in $O(\log |L|)$ time, and finding the two best scoring branches (which is what SHERPAS-full does) takes $O(1)$ time.

**Algorithm 2:** UPDATE$(L, x)$ (subprocess of Algorithm 1)

**Input:** An array $L$ satisfying the heap property for a score vector $S$. An element $x$ of $L$ whose score $S[x]$ was modified.

**Output:** The list $L$ that satisfies the heap property for the new value of $S[x]$.

```
set j as the position of x in L
if S[x] = −∞ then
    set L[j] = L[|L|]
    remove the last element of L from L
set hp = 0
while hp = 0 do
    set p = ⌊j/2⌋
    set l = 2j, r = 2j + 1
    if S[L[j]] > S[L[p]] then
        swap the contents of L[j] and L[p]
        set j = p
    else if (l ≤ |L| and S[L[l]] > S[L[j]]) or (r ≤ |L| and S[L[r]] > S[L[j]]) then
        if l = |L| or S[L[l]] ≥ S[L[r]] then
            set b = l
        else
            set b = r
        swap the contents of L[j] and L[b]
        set j = b
    else
        set hp = 1
return L
```

#### 2.4 Converting scores into likelihoods

To compare the scores of different branches, these scores must first be converted into numbers that can be interpreted as likelihoods or probabilities. Recall the definition of $S[x]$:

$$S[x] = \sum_{i=1}^{|W|} \log \widetilde{\mathbb{P}}[w_i|x]$$

Because $\widetilde{\mathbb{P}}[w_i|x]$ is defined as the product of the probabilities of the nucleotides in $w_i$ at a given placement, the log-probability of each nucleotide in the sum above is usually counted $k$ times (the only exception being the nucleotides near the ends of the window). Thus it makes sense to: (i) divide $S[x]$ by $k$, and then (ii) convert the resulting log-probability into a probability. This leads to the following definition:

$$\ell_x = b^{S[x]/k}, \tag{5}$$

where $b$ is the base of the logarithm (10 in current databases). We interpret $\ell_x$ as an approximation of the likelihood of $x$ being the phylogenetic origin of the sequence in $W$.

A more detailed justification for definition (5) is developed in Annex B, where the assumptions behind the argument above are made explicit.

##### 2.5 Producing the output

For any given window, procedure GET_STRAIN$(L, \theta)$ associates a classification (either a strain or N/A, which stands for *not assigned*) to that window on the basis of the contents of $L$. This involves converting the scores of the two best-scoring branches (SHERPAS-full) or all branches (SHERPAS-reduced) in $L$ into likelihoods $\ell_x$, in the way described in Sec. 2.4.

**Algorithm 3:** GET_STRAIN$(L, \theta)$ (subprocess of Algorithm 1)

**Input:** A max-heap $L$ of branches for window $W_i$ and a threshold $\theta$
**Output:** A classification for window $W_i$

```
if full pkDB in use then
    set x_1 and x_2 to the best-scoring and the second best-scoring branch in L, resp.
    if x_1 and x_2 are assigned to the same strain or ℓ_{x_1}/ℓ_{x_2} ≥ θ then
        return the strain assigned to x_1
    else
        return N/A
else // reduced pkDB in use
    set r = ℓ_{L[1]} / Σ_{i=1}^{|L|} ℓ_{L[i]}
    if r ≥ θ then return the strain assigned to L[1]
    else return N/A
```

Given the output $P_1, \ldots, P_N$ of Algorithm 1, SHERPAS infers the breakpoints as follows. If for some $1 \leq i \leq N-1$, we have $P_i \neq P_{i+1}$, a breakpoint between a segment of origin $P_i$ and a segment of origin $P_{i+1}$ is placed between the middle point of the window $W_i$ and the middle point of window $W_{i+1}$. Note that by construction of the windows, these two positions are always consecutive.

By default, if an unassigned (N/A) segment is inferred between two segments associated to the same strain $X$, the segment is then reassigned to strain $X$, and the breakpoints at its ends are removed. Note that this option can be deactivated by the user (see SHERPAS GitHub page).

##### 2.6 Complexity analysis

We start by analyzing the computational complexity of processing a single query.

Each $k$-mer in the query is processed at most twice: once when it is added for the first time to the sliding window, and once when it is removed from it. Processing a $k$-mer means first retrieving $B(w)$ from the pkDB, and then, for each pair $(x, s)$ in $B(w)$, updating $S[x]$, $C[x]$ and $L$ (see again Algorithm 1 for notation). Assuming that $k$ is a constant, the retrieval and inspection of each element in $B(w)$ takes $O(|B(w)|)$ time. The same holds for the update of $S[\cdot]$ and $C[\cdot]$, as each element of $B(w)$ involves $O(1)$ operations of update. As for the update of $L$ following the change of one $S[x]$, very often this will involve no change, as the change in $S[x]$ is usually very small and thus it does not require any swap within the max-heap. However, the worst-case time complexity of updating $L$ is $O(\log |L|)$. Since this needs to be repeated for many branches appearing in $B(w)$, the worst-case time complexity of processing a single $k$-mer $w$ is $O(|B(w)| \log |L|)$. Now note that $L$ can only contain candidate branches taken from $E$, and therefore $|L| \leq |E|$. The worst-case time complexity of processing a single $k$-mer $w$ is therefore

$$O(|B(w)| \log |E|)$$

The above analysis covers the time spent executing most of the code within the **for** $i = 1$ to $N$ loop in Algorithm 1, but it does not cover the part of the code within **if** $|W_i| \neq |W_{i-1}|$, nor the time spent executing GET_STRAIN. Whenever $|W_i| \neq |W_{i-1}|$ is true, SHERPAS updates the score of $O(|L|) = O(|E|)$ branches. But $|W_i| \neq |W_{i-1}|$ is only true for a constant number of windows (about 200 with default parameters), which means that the overall contribution to the running time of SHERPAS of this **if** clause is $O(|E|)$.

Finally, GET_STRAIN runs in $O(1)$ time for SHERPAS-full and $O(|E|)$ time for SHERPAS-reduced, because the calculation of the likelihood ratio requires reading the scores of 3 branches and $|L|$ branches for SHERPAS-full and SHERPAS-reduced, respectively. Now note that GET_STRAIN is called once for each window and that the number of windows is $O(|Q|)$, where $|Q|$ is the number of $k$-mers in the query $Q$. Therefore, the total contribution of GET_STRAIN to the running time of SHERPAS is $O(|Q|)$ and $O(|Q||E|)$ for SHERPAS-full and SHERPAS-reduced, respectively.

If we let $w_1, w_2, \ldots, w_{|Q|}$ be the sequence of $k$-mers that make up $Q$, then we can put together the observations of the last three paragraphs, and obtain an overall time complexity of

$$O(|E| + \sum_{i=1}^{|Q|} |B(w_i)| \log |E|) \quad \text{for SHERPAS-full}$$
$$O(|Q||E| + \sum_{i=1}^{|Q|} |B(w_i)| \log |E|) \quad \text{for SHERPAS-reduced}$$

These expressions can be simplified: let $\bar{B}_Q$ denote the average size of $B(w)$ across all $k$-mers in $Q$. In other words, we have $|Q|\bar{B}_Q = \sum_{i=1}^{|Q|} |B(w_i)|$. This allows us to rewrite the complexities above:

$$O(|E| + |Q|\bar{B}_Q \log |E|) \quad \text{for SHERPAS-full}$$
$$O(|Q||E| + |Q|\bar{B}_Q \log |E|) \quad \text{for SHERPAS-reduced}$$

Now note that while $\bar{B}_Q$ is usually very close to $|E|$ in SHERPAS-reduced (they are both small numbers), the same cannot be said in general for SHERPAS-full, where depending on the pkDB (and to a lesser extent on the query $Q$), we may have $\bar{B}_Q \ll |E|$ (in which case we say that the pkDB is *sparse*) or $\bar{B}_Q \approx |E|$ (the pkDB is *dense*). Moreover, the number of candidate branches $|E|$ is very different for SHERPAS-full and SHERPAS-reduced. For SHERPAS-full, $|E| = O(|T_0|)$, where $|T_0|$ is any measure of the size of the reference tree (e.g. $|T_0| = |E(T_0)|$). For SHERPAS-reduced, $|E| = |R_0|$, where $R_0$ denotes the set of root branches in $T_0$. Because of these observations, we conclude the following worst-case time complexities for a single query $Q$:

$$\begin{aligned} O(|T_0|+|Q|\bar{B}_Q \log |T_0|) &\quad \text{for SHERPAS-full and a sparse pkDB} \\ O(|Q||T_0| \log |T_0|) &\quad \text{for SHERPAS-full and a dense pkDB} \\ O(|Q||R_0| \log |R_0|) &\quad \text{for SHERPAS-reduced, with } |R_0| \ll |T_0|. \end{aligned}$$

As for memory complexity, SHERPAS uses $O(|E|)$ space to store all the information on candidate branches, including $S[\cdot], C[\cdot]$ and $L$. In practice, most of the memory employed by SHERPAS is used to store the pkDB, which is proportional to $\sum_w |B(w)|$ and therefore also $O(|E|)$, with a very large multiplicative constant proportional to the number of phylo-$k$-mers. Although in theory $O(|Q|)$ space is used to store the query and the partial results $P_1, \ldots, P_N$, in practice this is dominated by the $O(|E|)$ term. Therefore the space complexity for SHERPAS is $O(|E|)$.

To extend the analyses above for multiple queries, for running times it suffices to add together the complexities for the single queries. Memory complexity remains $O(|E|)$.

### 3 Assumptions on the reference data

Below we discuss a number of assumptions that, ideally, the reference data should satisfy. As we will illustrate in the subsections below, many of these assumptions were violated to some extent in the experiments we report in the paper. This means that some negative effects that these violations may have on the accuracy of SHERPAS may already be accounted by the results that we reported. This makes us optimistic about the robustness of SHERPAS to such violations.

#### 3.1 Accuracy of the reference tree

Like any other method based on a phylogenetic model, SHERPAS was developed assuming that the input phylogenetic tree is an accurate reflection of the evolutionary history of the sequences in the reference alignment. Violations of this assumption will result in phylo-$k$-mers having scores that do not reflect optimally their phylogenetic origin.

Although we have not investigated it extensively, we believe that SHERPAS should exhibit some robustness to the use of a reference tree containing a few errors. In fact, we believe that the trees that we used in our experiments are likely to contain a few differences with the reality, most notably because some ancestors of the reference sequences have probably undergone recombination. In this case *no* reference tree is an accurate description of the reality. We discuss the issue of hidden recombination in the next section.

The position of the root in the reference tree also has an influence on the pkDB construction step and on the scores of the phylo-$k$-mers. We refer the reader to the Supplementary Materials of the RAPPAS paper [11], where this point was discussed at length. For the purpose of recombination detection, the user should make sure that the root is placed outside the clades that are monophyletic with respect to the strains (see Sec. 3.4). Aside from that, it is definitely a good idea to place the root of the reference tree in a realistic position, but we do not think this will have a major influence on the accuracy of SHERPAS. In our experiments for HIV, we placed the root of the reference tree on the branch directly ancestral to strain O, which is probably not the correct position of the root for the reference tree.

#### 3.2 Absence of recombination

A standard assumption of phylogeny-based methods for recombination detection is that the reference alignment is recombination-free, that is, none of the sequences in the alignment, nor their ancestors, have undergone recombination events. See for example the discussion on this point in the paper about SCUEAL [8]. However, other tools are inherently robust to the inclusion of some types of recombinants, as their models do not depend on a particular phylogenetic tree. For example this is the case for jpHMM. Being phylogeny-based, SHERPAS is closer to SCUEAL in this respect. However, we can expect that minor violations to the assumption that the alignment is recombination-free can be tolerated. We discuss the possible effect of a number of possible violations below.

First, it is likely that even in the reference alignments used by many phylogeny-based methods, sequences that are “pure” for one strain are in fact *intra*-strain recombinants. These recombinants —which are difficult to detect, as they involve recombination events between similar sequences— are believed to be relatively frequent, for example in HIV [7] and HBV [12]. Including intra-strain recombinants in the reference alignment is probably not only inevitable, but also relatively innocuous for SHERPAS. This is because the errors it causes in the reference phylogeny will presumably only involve branches that are internal to one strain. These errors are likely to have a limited effect on the scores of some phylo-$k$-mers, and they should not cause the misclassification of a window in an incorrect strain. Note that most of our experiments with SHERPAS were done using the same reference alignments as jpHMM, which to the best of our knowledge were not screened to exclude intra-subtype recombinants.

Second, sometimes entire strains may be composed of recombinant sequences. This occurs when a recombination event was ancestral to a common ancestor of all the sequences in the strain. This is believed to be the case for *circulating recombinant forms* (CRFs), which are recombinant forms that have been observed in several unrelated individuals. A naïve approach to include sequences of a particular CRF in a reference alignment is to define a strain coinciding with that CRF. This is what we have done in the two HIV datasets for CRF01_AE, which is believed to be a recombinant between subtype A and a now extinct (or unsampled) subtype E. Setting a strain to a CRF allows users to recognize novel recombinants between that CRF and other strains.

However, the naïve approach above comes at a risk: the true evolutionary history of the reference sequences involves at least one reticulation ancestral to the CRF, meaning that the reference tree —whatever it is— will definitely not be correct in the part ancestral to that CRF. We do not know how strong the effect of using such an incorrect tree is, but it is probably more important than the effect of including intra-strain recombinants. For this reason, with the exception of CRF01_AE, we have avoided the inclusion of CRFs in the reference alignment, and we advise users to do the same. In the HIV-genome dataset we have noticed that SHERPAS appears to have some difficulties distinguishing CRF01_AE from strain A1 (this can be observed for example in queries 18, 51, 65 in Section C below), which is related to the fact that the relative position of CRF01_AE and the A strains within the reference tree is probably erroneous for at least part of the reference alignment. In the next subsection we discuss how an advanced user may include CRFs in the reference alignment in a clever way, which addresses in a satisfactory way the problems above.

Third, some sequences may be erroneously annotated as belonging to one strain, but in fact they are *inter*-strain recombinants. Compared to the other cases discussed above, these are definitely the recombinants that may cause the most serious disruption to the accuracy of SHERPAS —and of other methods too, including jpHMM and SCUEAL. **Users should always make sure that the reference alignment only contains sequences that are "pure" with respect to the input strains.** In our experiments, we have always used the reference alignments that were distributed with either jpHMM or SCUEAL.

##### 3.3 Dealing with circulating recombinant forms

Including CRFs in the reference alignment and assigning them to strains labelled by the CRF identifier is attractive from a practical point of view because it may allow users of SHERPAS to (1) recognize if a sequence is a CRF and (2) detect novel recombinants between a CRF and another strain. However, as explained above, the naïve way of doing so is probably not a good idea, as the reference tree will contain some large-scale differences with the reality.

If tasks (1) and (2) are important to a user, there is a relatively simple way of solving this problem, which we explain below.

This idea was already explained, among others, by Sergei Kosakovsky Pond and coauthors [8] and Darren Martin (personal communication). It exploits the fact that, because CRFs have been the object of several analyses, their recombinant structure is well-known, including the (sometimes approximate) position of breakpoints. This means that any CRF sequence can be decomposed into subsequences that are "pure" with respect to their strains. For example in HIV-1, the sequences tagged as CRF03_AB have mosaic structure ABA with the central B segment starting at site 2688 and ending at site 8649 [10]. In this case, every sequence $x$ from CRF03_AB should be decomposed into the following two sequences prior to their inclusion in the reference alignment: $x_A$, which is identical to $x$, except for positions 2688-8649 which are replaced by a long stretch of gaps; and $x_B$, a sequence that has gaps up to position 2687, then coincides with $x$ from position 2688 to 8649, and finally has gaps up to the last position of $x$. These two sequences can then be assigned to strain CRF03_AB. Note that $x_A$ and $x_B$ have different phylogenetic origins within the reference tree, with $x_A$ likely to be found near subtype A and $x_B$ likely to be found near subtype B. In fact all the sequences obtained as $x_A$ from some sequence $x$ from CRF03_AB should form a clade, and the same holds for all sequences obtained as $x_B$.

In general, if we subdivide each reference sequence from a CRF into its non-recombinant components, then it is not a problem anymore to represent the evolution of the reference sequences with a phylogenetic tree. The strain corresponding to a CRF will be partitioned in as many sub-strains as there are parental sequences for that CRF, and generally each of these sub-strains will form a separate clade in the reference tree. Since SHERPAS is able to deal with strains that are polyphyletic (see next section), it can be used on the pkDB constructed above without modification.

##### 3.4 Monophyly of the strains

Since strains are usually defined either using phylogenetic criteria or by adopting appropriate thresholds for intra-strain similarity, we expect them to be usually monophyletic, meaning that each strain should correspond to a separate clade in the reference tree. In fact it is always a good idea for users of SHERPAS to check that most strains are monophyletic after they reconstructed, or downloaded, the reference tree. A few exceptions to this rule are acceptable, for example when (1) an "artificial" strain is designed to contain all reference sequences that are not in a strain of interest, or (2) the user knows that the strain is not monophyletic, yet it makes sense to define it as a single strain. As an example of (2), in our experiments on HIV, we used a strain named CPZ to include all SIV (simian immunodeficiency virus) sequences from chimpanzees, which are expected to be polyphyletic. For another example of why (2) may be useful, see Sec. 3.3 above.

Because of the possible exceptions to monophyly, SHERPAS was designed to tolerate the inclusion of polyphyletic strains. However, this will have some subtle effects on the behavior of SHERPAS, which may differ depending on the version (full/reduced) of the pkDB used. Specifically, the presence of polyphyletic strains increases the number of branches that are unassigned (not assigned to any strain) in SHERPAS-full and increases the number of root branches in SHERPAS-reduced. Although our experiments did include a few polyphyletic strains (3 in HIV-pol and 2 in HIV-genome, including CPZ), we do not have enough perspective to describe their effect on the analysis. Because of this, we believe that they should be used with caution.

### 4 Site-wise measures of accuracy

Here we show some simple mathematical relationships between the site-wise sensitivity and precision defined in the Materials and Methods of the paper, and strain-specific definitions of sensitivity and precision, which are standard in multi-class classification literature.

Let $n$ be the number of strains. For ease of notation, we associate each strain to one integer from 1 to $n$, whereas we associate integer $n+1$ to the "unassigned" status.

We define the *confusion matrix* as an $n \times (n+1)$ matrix $M = (m_{ij})$, where $m_{ij}$ is the number of sites whose correct strain is $i$ and that were classified as $j$.

The following two definitions are standard in multi-class classification.

- The *sensitivity for strain* $i$ $(1 \leq i \leq n)$ is the proportion of sites that are classified as $i$, out of all sites whose correct strain is $i$. That is, in terms of the confusion matrix:

$$\text{sensitivity}_i = \frac{m_{ii}}{\sum_{j=1}^{n+1} m_{ij}}.$$

- The *precision for strain* $i$ $(1 \leq i \leq n)$ is the proportion of sites whose correct strain is $i$, out of all sites that are classified as $i$. That is, in terms of the confusion matrix:

$$\text{precision}_i = \frac{m_{ii}}{\sum_{j=1}^{n} m_{ji}}.$$

The following two definitions are the ones that we gave in the paper.

- The *site-wise sensitivity* is the proportion of sites that are classified in their correct strain, out of all sites. That is, in terms of the confusion matrix:

$$\text{site-wise sensitivity} = \frac{\sum_{i=1}^{n} m_{ii}}{\sum_{i=1}^{n} \sum_{j=1}^{n+1} m_{ij}}.$$

- The *site-wise precision* is the proportion of sites that are classified in their correct strain, out of all sites that are classified in some strain other than $n+1$. That is, in terms of the confusion matrix:

$$\text{site-wise precision} = \frac{\sum_{i=1}^{n} m_{ii}}{\sum_{i=1}^{n} \sum_{j=1}^{n} m_{ij}}.$$

The two pairs of definitions above are linked in an intuitive way by the following two propositions.

**Proposition 1.** *The site-wise sensitivity is the weighted average of the strain-specific sensitivities, where the weight for strain* $i$ *is proportional to the number of sites whose correct strain is* $i$*.*

*Proof.* The weight $w_i$ for class $i$ is proportional to $\sum_{j=1}^{n+1} m_{ij}$. Since the sum of weights in a weighted average is 1, we have:

$$w_i = \frac{1}{S}\sum_{j=1}^{n+1} m_{ij},$$

where

$$S = \sum_{i=1}^{n}\sum_{j=1}^{n+1} m_{ij}.$$

Then, the weighted average in the statement is given by:

$$\begin{aligned}\sum_{i=1}^{n} w_i \cdot \text{sensitivity}_i &= \sum_{i=1}^{n}\left(\frac{1}{S}\sum_{j=1}^{n+1} m_{ij}\right)\cdot\frac{m_{ii}}{\sum_{j=1}^{n+1} m_{ij}}\\ &= \frac{1}{S}\sum_{i=1}^{n} m_{ii}\\ &= \text{site-wise sensitivity.}\end{aligned}$$

□

**Proposition 2.** *The site-wise precision is the weighted average of the strain-specific precisions, where the weight for strain i is proportional to the number of sites that are classified as i.*

*Proof.* The weight $w'_i$ for class $i$ is proportional to $\sum_{j=1}^{n} m_{ji}$. Since the sum of weights in a weighted average is 1, we have:

$$w'_i = \frac{1}{S'}\sum_{j=1}^{n} m_{ji},$$

where

$$S' = \sum_{i=1}^{n}\sum_{j=1}^{n} m_{ji}.$$

Then, the weighted average in the statement is given by:

$$\begin{aligned}\sum_{i=1}^{n} w'_i \cdot \text{precision}_i &= \sum_{i=1}^{n}\left(\frac{1}{S'}\sum_{j=1}^{n} m_{ji}\right)\frac{m_{ii}}{\sum_{j=1}^{n} m_{ji}}\\ &= \frac{1}{S'}\sum_{i=1}^{n} m_{ii}\\ &= \text{site-wise precision.}\end{aligned}$$

□

### 5 Dataset construction, data availability, and reproducibility

Here we describe a number of details about the construction of the datasets behind the experiments presented in the main text. We also provide the information necessary for their reproducibility. All relevant files that do not belong to a third party can either be downloaded from the SHERPAS GitHub repository at https://github.com/phylo42/sherpas or from the Dryad dataset distributed with this paper (link provided on the GitHub README page).

#### 5.1 Preprocessing

The operations necessary prior to the execution of SHERPAS —i.e. the construction of the reference tree, of the phylo-$k$-mer database, and of the sequences-to-strains mapping— were the same for all the datasets. We briefly describe them here.

Reference trees were constructed from the reference alignments (dataset-specific, see below) using PhyML 3.3.20180214 [5] using GTR+Γ+I as nucleotide substitution model. A discrete Γ distribution with 4 rate categories (the default in PhyML) was used to model rate variation across sites. The proportion of invariant sites was estimated from the alignment, as well as the shape parameter $\alpha$ (the latter is done by default by PhyML). The resulting reference trees are included in the Dryad dataset. PhyML was run with the following command:

```
phyml -i <reference_alignment> -m GTR -v e
```

The pkDBs were constructed using RAPPAS2 v0.1.3a, using parameter $k = 10$ (which is not the default value) and threshold parameter $\omega = 1.5$ (the default). (See also Sec. 1.3 above for the meaning of the $\omega$ parameter.) The resulting pkDBs are included in the Dryad dataset (`.rps` files). RAPPAS2 was run with the following command:

```
python3 rappas2.py build -s nucl -b <path_to_phyml> -w <output_directory>
        -r <reference_alignment> -t <reference_tree> -m GTR -k 10
```

Finally, the `.csv` files mapping reference sequences to strains were constructed manually, using the information contained in the name of each sequence, to infer the strain it belongs to. These files are also included in the Dryad dataset.

#### 5.2 HIV-pol

**References.** The reference alignment that we used for SHERPAS is the one provided with SCUEAL (167 sequences), and is available at https://github.com/spond/SCUEAL/blob/master/data/pol2009.nex. (Accessed October 2019.)

**Queries.** The 10,000 synthetic queries and the output of SCUEAL for these queries are accessible here: http://www.hyphy.org/pubs/SCUEAL/SCUEAL%20Files_files/Shuffled.zip. (Accessed October 2019.) Note that none of the queries contains fragments from strains A3, AE, O, CPZ, although these strains are present in the reference alignment. All other strains in the reference alignment are present in some queries.

**Running SHERPAS.** The reference tree reconstructed from the reference alignment, the .csv file recording the classification of the sequences in the reference alignment, and the .rps file encoding the pkDB for this dataset are available in the `pkDB-HIV-pol` directory within the Dryad dataset.

SHERPAS was executed with the following command:

```
SHERPAS -o out/ -d pkDB-HIV-pol/DB_k10_o1.5.rps
        -q HIV_pol/queries.fasta -g pkDB-HIV-pol/ref-groups.csv
```

**Running jpHMM.** jpHMM was executed with default parameters for HIV sequences, with the following command:

```
./jpHMM -s HIV_pol/queries.fasta -v HIV
```

It was also executed with the `-Q blat` option for speed-up. Out of the several files produced by jpHMM, the output file that was used to read the predictions made by jpHMM is `recombination.txt`.

Note that on this dataset the reference alignment used by jpHMM does not coincide with that used by the other methods. Moreover, as discussed in the main text, strains A and N cannot be recognized by jpHMM, which has a slightly negative impact on jpHMM's accuracy measures on this dataset.

In order to understand how strong this impact is, we also analyzed the results of jpHMM under the assumptions that returning A1 or A2 is correct when A is the true strain, and that CPZ is correct when N is the true strain. In this case, jpHMM returns 92.9% matches, 0% supersets and 6.9% subsets, and its sensitivity and precision equal 99.3%.

#### 5.3 HBV-genome

**References.** We used the reference alignment that is distributed with jpHMM. It can be downloaded at `http://jphmm.gobics.de/download.html`, and is located at `jpHMM/input/HBV_alignment.fas`. (Accessed December 2019.) This alignment contains 339 whole-genome sequences classified into strains A, B, C, D, E, F, G, H. Prior to the construction of the pkDB for SHERPAS, we extended this reference alignment by copying the first $k - 1 = 9$ columns of the alignment to the end of the alignment. This allows the construction of phylo-$k$-mers (with $k = 10$) from positions that overlap with the artificial end of the alignment.

**Queries.** We downloaded the dataset for nucleotide sequence genomes of "genotype all" from the HBVdb website at `https://hbvdb.ibcp.fr/HBVdb/HBVdbDataset?view=/data/nucleic/alignments/all_Genomes.clu&seqtype=0`. (Accessed December 2019.) This alignment contained 7273 sequences, from which we removed 794 sequences marked as recombinant and 273 sequences that belonged to the jpHMM reference alignment. We also removed 1664 sequences that were estimated to be recombinant by both jpHMM and SHERPAS (in fact some of these turn out to be known recombinants [12]).

The remaining 4542 sequences are the pre-queries that were used as a basis to build 3000 recombinant queries. Out of these 3000 queries, 2000 combine fragments from two pre-queries, and 1000 are based on three pre-queries. To generate each query, we proceeded as follows. First, the 2 or 3 pre-queries are chosen at random from the 4542, making sure that they belong to different strains. Second, we picked $2X$ random coordinates within the alignment of pre-queries, where $X \geq 1$ is chosen following a geometric distribution with parameter $p = 0.8$. The coordinates are chosen uniformly at random, making sure that no two coordinates are less that 100 positions apart (circularity-wise). This is the same minimal distance that was applied to build the dataset of queries for HIV-pol [8]. Third, each query is built by combining segments extracted from the selected pre-queries and delimited by the random coordinates.

The parameters that we used in the construction procedure above were chosen so as to loosely reflect the characteristics of inter-genotype HBV recombinants presented in a recent overview [1]. For example, about 80% of the recombinants involving 2 genotypes in Fig. 2 of that review only have 2 breakpoints, hence the choice of $p = 0.8$.

The queries are available here: `https://github.com/phylo42/sherpas/blob/master/examples/HBV_all/queries-3000.fasta`.

**Running SHERPAS.** The reference tree reconstructed from the reference alignment, the .csv file recording the classification of the sequences in the reference alignment, and the .rps file encoding the pkDB for this dataset are available in the `pkDB-HBV-full` directory within the Dryad dataset (link above).

SHERPAS was executed using the option `-c` (for circular queries) with the following command:

```
SHERPAS -o out/ -d pkDB-HBV-full/DB_k10_o1.5.rps
        -q HBV_all/queries-3000.fasta -g pkDB-HBV-full/ref-groups.csv -c
```

**Running jpHMM.** jpHMM was executed using the option `-C` (for circular queries):

```
./jpHMM -s HBV_all/queries-3000.fasta -v HBV -C
```

Note that the `-C` option automatically activates the `-Q blat` option. Out of the several files produced by jpHMM, the output file that was used to read the predictions made by jpHMM is `recombination.txt`.

##### 5.4 HIV-genome

**References.** The reference alignment was obtained as part of the jpHMM package which can be downloaded here: `http://jphmm.gobics.de/download.html`, and is located at `jpHMM/input/HIV_alignment.fas`. (Accessed December 2019.) This alignment contains 881 whole-genome sequences, classified in the following 14 strains: A1, A2, AE, B, C, D, F1, F2, G, H, J, K, O, CPZ.

**Queries.** We downloaded the Los Alamos Web alignment 2018 available at https://www.hiv.lanl.gov/content/sequence/NEWALIGN/align.html (accessed January 2020), using the following parameters. *Alignment type*: Web alignment (all complete sequences). *Year*: 2018. *Organism*: HIV-1/SIVcpz. *DNA/Protein*: DNA. *Region*: genome. *Subtype*: ALL. *Format*: FASTA. *Alignment ID*: 118AG1. This alignment consisted of 4004 sequences, from which we removed 1328 sequences that do not belong to any of the 14 strains above (CRFs other than CRF01_AE, see Sec. 3.3 above), and also removed 803 sequences that are present in the jpHMM reference alignment. Finally, we removed 27 sequences that were identified as recombinant by both jpHMM and SHERPAS.

We constructed 3000 synthetic recombinant sequences from the remaining 1846 pre-queries. For each of the 3000 sequences, we started by drawing $X \geq 1$ and $Y \geq 1$ from geometric distributions with parameters 0.2 and 0.8 respectively, discarding any pair with $X < Y$. $X$ and $Y + 1$ represent the number of breakpoints in the query and the number of pre-queries involved, respectively. We picked $X$ coordinates within the alignment of pre-queries using a uniform distribution, making sure that no two coordinates are less that 100 sites apart and no coordinate is less than 100 sites away from one end of the query. We also drew $Y + 1$ pre-queries at random, making sure that at least two of these pre-queries are from different strains. Finally, we built a query sequence by inserting, between each pair of consecutive coordinates, the corresponding segment in one of the selected pre-queries, making sure that the pre-queries chosen for any two consecutive intervals are different. The length of the resulting queries is on average 8.9 kbp, and it ranges from 5.5 kbp to 9.9 kbp. The longest queries are obtained from pre-queries whose length is close to 1 Mbp (which are common e.g. in group O or when sampled from chimpanzees).

The queries are available at https://github.com/phylo42/sherpas/blob/master/examples/HIV_all/queries-C.fasta.

**Running SHERPAS.** The reference tree reconstructed from the reference alignment, the .csv file recording the classification of the sequences in the reference alignment, and the .rps file encoding the pkDB for this dataset are available in the `pkDB-HIV-full` directory within the Dryad dataset (link above).

SHERPAS was executed with the following command:

```
SHERPAS -o out/ -d pkDB-HIV-full/DB_k10_o1.5.rps
        -q HIV_all/queries-C.fasta -g pkDB-HIV-full/ref-groups.csv
```

**Running jpHMM.** jpHMM was executed with default parameters for HIV sequences:

```
./jpHMM -s HIV_all/queries-C.fasta -v HIV
```

It was also executed with the `-Q blat` option for speed-up. Out of the several files produced by jpHMM, the output file that was used to read the predictions made by jpHMM is `recombination.txt`.

##### 5.5 HIV Nanopore reads dataset

**References.** The reference alignment, the reference tree and the pkDB are the same as in the HIV-genome dataset.

**Queries.** We ran NanoSim-H (v1.1.0.4) on the queries from the HIV-genome dataset using the following command:

```
nanosim-h HIV_all/queries-C.fasta -n 100000 -o reads-C.fasta -u 0.0 --max-
    len 9000 --min-len 1000
```

This generates 100,000 reads, each one from a random query of the HIV-genome dataset. For each one of these sequences, we picked the first (in order of appearance) read in the forward strand that comes from that sequence. The resulting queries are available at https://github.com/phylo42/sherpas/blob/master/examples/HIV_all/reads-C.fasta.

**Running SHERPAS and jpHMM.** The same commands as for the HIV-genome dataset were used here (replace the name of the file containing the queries). The execution of jpHMM with the `-Q blat` option aborted.

##### 5.6 Experiment on the specificity/recall trade-off

**Measures of accuracy.** Given a binary classifier $\mathcal{C}$ for the detection of inter-strain recombinant sequences, and a collection of query sequences, we define:

**TP** (*true positives*): the number of queries that are both inter-strain recombinants and classified as such by $\mathcal{C}$.

**FP** (*false positives*): the number of queries that are erroneously classified by $\mathcal{C}$ as inter-strain recombinants.

**FN** (*false negatives*): the number of queries that are inter-strain recombinants, but not classified as such by $\mathcal{C}$.

**TN** (*true negatives*): the number of queries that are neither inter-strain recombinants, nor classified as such by $\mathcal{C}$.

**Recall** (also known as *true positive rate* or *sensitivity*): out of all inter-strain recombinant queries, the proportion that are recognized as such by $\mathcal{C}$. That is, Recall $= \frac{\text{TP}}{\text{TP}+\text{FN}}$.

**Specificity** (also known as *true negative rate*): out of all queries that are *not* inter-strain recombinants, the proportion that are not classified as inter-strain recombinants. That is, Specificity $= \frac{\text{TN}}{\text{TN}+\text{FP}}$.

**Dataset.** Among the 10,000 queries in the HIV-pol dataset, there are 2374 queries that are not inter-strain recombinants. Thus this dataset provides a way to compute not only the recall of a classifier, but also its specificity. Moreover, on this dataset, the results of running SCUEAL were made available by the authors [8], which allows us to compare the results of SHERPAS to those of jpHMM *and* SCUEAL. Since jpHMM is not designed to recognize sequence fragments labelled as A or N, for this experiment we removed all queries from the initial 10,000 that were obtained from sequences from these two strains. This ensures that all classifiers compete on an equal ground, and leaves us with 7652 queries, of which 5317 are inter-strain recombinants.

**Classifiers.** For all tested programs, a query is classified as an inter-strain recombinant if and only if the partition produced by the program contains regions from 2 or more different strains. jpHMM was run with and without the `-Q blat` option, but the results were virtually indiscernible. SHERPAS-full was run for all combinations of window size in $\{300, 500\}$ and threshold parameter $\theta_F \in \{1, 2, 5, 10, 20, 50, 100, 200, 500, 1000\}$. SHERPAS-reduced was run for all combinations of window size in $\{300, 500\}$ and threshold parameter $\theta_R \in \{0, 0.5, 0.75, 0.9, 0.95, 0.975, 0.99, 0.995, 0.9975, 0.999\}$.

**Discussion of the results.** Results of this experiment are reported in Fig. 2 of the main text. A number of observations can be made here. First, while SCUEAL and jpHMM result in good (Pareto-optimal) combinations of specificity and recall, it is interesting to note that on this dataset jpHMM has better (100%) specificity, while SCUEAL has better recall. Previously reported results on other datasets do not appear to confirm this observation [13], but this is not surprising because those results were on a different but related classification problem, namely typing whole sequences, i.e. assigning them to their strain of origin. Moreover SCUEAL implements a non-deterministic optimization heuristic, with better results expected if SCUEAL is left to run for longer times.

As for SHERPAS, window size has a clear effect, with small windows favoring recall, and larger windows favoring specificity. This is consistent with the results in Tables 2 – 5 in the main text, where smaller window sizes are generally associated with more fragmented mosaics. Interestingly the threshold $\theta_R$ has a very strong control over the trade-off between recall and specificity in SHERPAS-reduced, with small values of $\theta_R$ causing very high recall, and large $\theta_R$ causing very high specificity. This is not the case for SHERPAS-full, where $\theta_F$ only has a limited effect on specificity and recall.

This difference in the behaviors of SHERPAS-reduced and SHERPAS-full can be understood by considering again the way these algorithms use their respective thresholds (see Sec. 2.5 above, and Algorithm 3). Consider the case where a region of the query has weak evidence for belonging to strain $X$. While $\theta_R$ entirely controls whether $X$ is present in the partition produced by SHERPAS-reduced, $\theta_F$ only plays a role in determining the output of SHERPAS-full if the two best-scoring branches are from different strains. If the two best-scoring branches in SHERPAS-full are both from $X$, then $X$ will be present in the output partition, irrespective of the value of the threshold. This is why the specificity for SHERPAS-full cannot be increased up to 100% by increasing its threshold.

Coming back to the question that motivated this experiment —namely the use of SHERPAS as a first screening tool for inter-strain recombinants— finally note that very high recall can be achieved with both SHERPAS-reduced and SHERPAS-full by using small window sizes and low thresholds. On this dataset, these parameter settings result in recall that is substantially higher than jpHMM, and at least as good as SCUEAL (up to 99.0% for SHERPAS-full, and up to 99.4% for SHERPAS-reduced, vs. 98.8% for SCUEAL), for a fraction of their running times.

[14] Ziheng Yang. *Computational Molecular Evolution.* Oxford University Press, 2006.

#### A Annex: Marginal posterior distributions of ancestral states

Ancestral state reconstruction is a wide subject in phylogenetics. As explained in Section 1.2, constructing phylo-$k$-mers and calculating their probability scores relies on a specific technique for ancestral reconstruction: the calculation of the marginal posterior distribution of $A'_{u,i}$. (Recall that $A'_{u,i}$ represents the random nucleotide at node $u$ in the extended reference tree, homologous to site $i$ in alignment $A$.)

Here, we describe how this calculation is performed by standard software for phylogenetic inference (e.g. PhyML [6]) using the notation introduced in Section 1.2. More information can be found in many phylogenetics textbooks (e.g. [14, Chapter 4.4]).

Let $A_{*,i}$ denote the $i$th column of $A$. Because of the assumption of independent evolution at different sites, the probability that we seek to calculate only depends on the data in $A_{*,i}$. That is,

$$\mathbb{P}[A'_{u,i} = a \mid A, \theta] \quad = \quad \mathbb{P}[A'_{u,i} = a \mid A_{*,i}, \theta]. \tag{6}$$

The first ingredient to derive the probability above is the probability of the data in $A_{*,i}$, given $A'_{u,i} = a$, that is

$$\mathbb{P}[A_{*,i} \mid A'_{u,i} = a, \theta]. \tag{7}$$

This is sometimes referred to as a *partial likelihood* and can be obtained for every $a \in \{\mathtt{A}, \mathtt{C}, \mathtt{G}, \mathtt{T}\}$ by using Felsenstein's pruning algorithm [3, 4] on the extended reference tree $T$ rooted in $u$.

The other ingredient is the prior distribution of $A'_{u,i}$, which is simply given by

$$\mathbb{P}[A'_{u,i} = a \mid \theta] = \pi_a \tag{8}$$

where $(\pi_\mathtt{A}, \pi_\mathtt{C}, \pi_\mathtt{G}, \pi_\mathtt{T})$ is the stationary distribution implied by the substitution model. (E.g. for the Jukes-Cantor model all $\pi_a = 1/4$, while for many other models the $\pi_a$ are parameters included in $\theta$.)

Given the probabilities in (7) and (8), the probability in (6) can be obtained as

$$\mathbb{P}[A'_{u,i} = a \mid A_{*,i}, \theta] = \frac{\mathbb{P}[A'_{u,i} = a, A_{*,i} \mid \theta]}{\mathbb{P}[A_{*,i} \mid \theta]} = \frac{\mathbb{P}[A'_{u,i} = a \mid \theta] \cdot \mathbb{P}[A_{*,i} \mid A'_{u,i} = a, \theta]}{\sum_{a' \in \{\mathtt{A},\mathtt{C},\mathtt{G},\mathtt{T}\}} \mathbb{P}[A'_{u,i} = a' \mid \theta] \cdot \mathbb{P}[A_{*,i} \mid A'_{u,i} = a', \theta]}.$$

### B Annex: Approximate relationship between branch scores and likelihoods

Here we show that, under many simplifying assumptions, the quantity

$$\ell_x = b^{S[x]/k} \tag{9}$$

is approximately proportional to the likelihood of $x$ being the phylogenetic origin of the sequence in the current window $W$. Since the assumptions are very strong, we only expect this relationship to hold in the limit, as the assumptions get closer and closer to being verified. Even if the assumptions are never verified in practice, this result provides a justification for the way SHERPAS converts scores into likelihoods. The text below shows the details of the reasoning behind (9) and its assumptions.

Let $a_1 a_2 \ldots a_{m_W}$ be the nucleotide sequence in the current window $W$, where $m_W = |W|+k-1$. We use here the notation introduced in Section 2.1, where the $i$th $k$-mer in $W$ is $w_i = a_i a_{i+1} \ldots a_{i+k-1}$.

Our first assumption is that $a_1 a_2 \ldots a_{m_W}$ unequivocally aligns to a stretch of contiguous sites, going from site $s_W + 1$ to site $s_W + m_W$. In fact we also assume that $k$ is long enough for every $k$-mer $w_i$ to align unequivocally to the $k$ sites starting from $s_W + i$.

Now recall that $x \in E(T_0)$ denotes a branch of the reference tree and define tree $T_x$ as the tree that is obtained from $T_0$ by attaching a ghost branch to the midpoint of $x$, where $v_x$ is the new ghost leaf (see Figure 1). Finally, we place sequence $a_1 a_2 \ldots a_{m_W}$ at leaf $v_x$.

We can then define the likelihood of $x$ being the phylogenetic origin of $W$ as the likelihood of tree $T_x$, which we can then develop following standard phylogenetic calculations based on rooting $T_x$ at leaf $v_x$. Using the notation introduced in Section 1.2 and Annex A, we get:

$$\begin{aligned}
\mathrm{Lik}(T_x) &= \prod_{j=1}^{m_W} \mathbb{P}[\, A_{*,s_W+j} \mid A'_{v_x,s_W+j} = a_j \,,\, \theta \,] \cdot \pi_{a_j} \\
&= \prod_{j=1}^{m_W} \mathbb{P}[\, A'_{v_x,s_W+j} = a_j \mid A_{*,s_W+j} \,,\, \theta \,] \cdot \mathbb{P}[A_{*,s_W+j} \mid \theta] \\
&= \mathrm{const} \cdot \prod_{j=1}^{m_W} \mathbb{P}[A'_{v_x,s_W+j} = a_j \mid A_{*,s_W+j}, \theta]
\end{aligned}$$

where const is a constant that does not depend on $x$. The result above simply states that, as we vary $x$, the likelihood of $T_x$ is directly proportional to the posterior probability of having $a_1 a_2 \ldots a_{m_W}$ at leaf $v_x$ and at sites $s_W + 1$ to $s_W + m_W$.

Now define $p_j(x)$ as the posterior probability of having $a_j$ at leaf $v_x$ and site $s_W+j$, i.e. $p_j(x) = \mathbb{P}[A'_{v_x,s_W+j} = a_j \mid A_{*,s_W+j}, \theta]$. This allows us to express $\mathrm{Lik}(T_x)$ compactly:

$$\mathrm{Lik}(T_x) = \mathrm{const} \cdot \prod_{j=1}^{m_W} p_j(x) \tag{10}$$

Now consider $\widetilde{\mathbb{P}}[w_i|x]$, defined in (3) as $\widetilde{\mathbb{P}}[w_i|x] = \max_{u \in G_x} \max_j \mathbb{P}[\, S_{u,j} = w_i \mid A, \theta \,]$. Since $w_i$ aligns to the $k$ sites starting from $s_W + i$, and assuming that the probabilities associated to different nodes $u \in G_x$ are approximately equal, we have that

$$\widetilde{\mathbb{P}}[w_i|x] \approx \mathbb{P}[\, S_{v_x, s_W+i} = w_i \mid A, \theta\,] = \prod_{h=0}^{k-1} \mathbb{P}[A'_{v_x, s_W+i+h} = a_{i+h} \mid A, \theta].$$

Recall that, because sites are assumed to be independent, the posterior probabilities at a site only depend on the data at that same site. This implies

$$\widetilde{\mathbb{P}}[w_i|x] \approx \prod_{h=0}^{k-1} \mathbb{P}[A'_{v_x, s_W+i+h} = a_{i+h} \mid A_{*, s_W+i+h}, \theta] = \prod_{h=0}^{k-1} p_{i+h}(x).$$

Now combine the equation above to the definition of SHERPAS's scores (Eqn. (4), page 7):

$$S[x] = \log \prod_{i=1}^{|W|} \widetilde{\mathbb{P}}[w_i|x] \;\approx\; \log \prod_{i=1}^{|W|} \prod_{h=0}^{k-1} p_{i+h}(x)$$
$$= \log \Big( p_1(x)^1 \cdot p_2(x)^2 \cdots p_k(x)^k \cdot p_{k+1}(x)^k \cdots p_{|W|}(x)^k \cdot p_{|W|+1}(x)^{k-1} \cdots p_{|W|+k-2}(x)^2 \cdot p_{|W|+k-1}(x)^1 \Big)$$

By changing the first and last $k-1$ terms in the product above, so that all terms have exponent $k$ or 0, we obtain the following bounds:

$$\log \prod_{j=1}^{m_W} p_j(x) \lesssim \frac{S[x]}{k} \lesssim \log \prod_{j=k}^{|W|} p_j(x) \tag{11}$$

Compare this result to (10) and note that both the lower and the upper bound in (11) can be interpreted as log-likelihoods (albeit for slightly different sequences). By exponentiating we get the following approximate relationship:

$$b^{\frac{S[x]}{k}} \approx \text{const}' \cdot \text{Lik}(T_x),$$

which is what we set out to show.

#### C Annex: Some illustrated outputs

To provide readers with a feeling of how similar the results of SHERPAS and jpHMM are, here we show an illustration of the outputs of SHERPAS-full (with default parameters) and jpHMM on the first 100 queries out of the 3000 in the HIV-genome dataset.

For each query, three bars show respectively: (top) the true composition of the query, with different strains represented by different colors; (mid) the output of SHERPAS-full, (bottom) the output of jpHMM. Black regions are those that are left unassigned (N/A) by the recombination detection tool. Each colored fragment is drawn to scale. Below the three bars, we also report the color coding for the strains present in at least one of the three bars.

It is helpful to look at these results together with those reported in Table 4 in the main text.

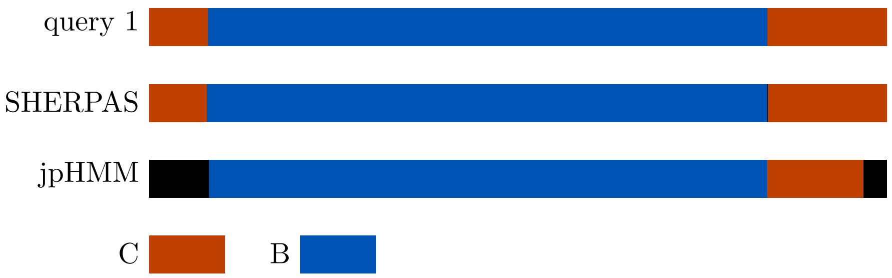

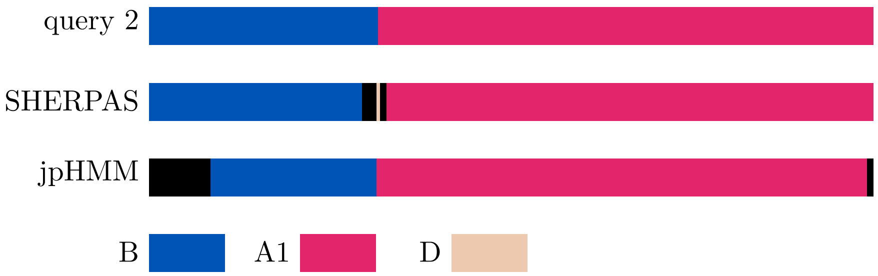

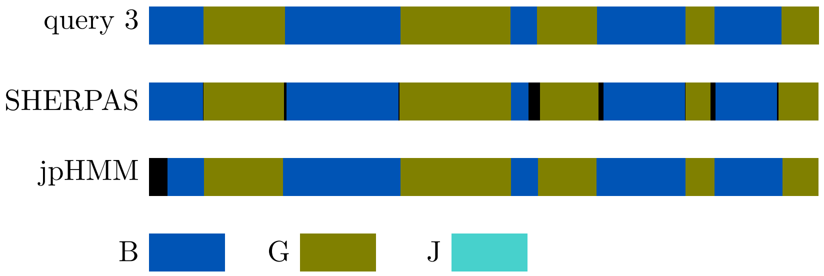

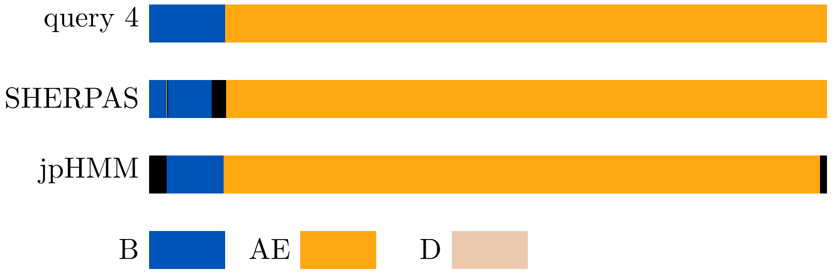
query 4
SHERPAS
jpHMM
B
AE
D

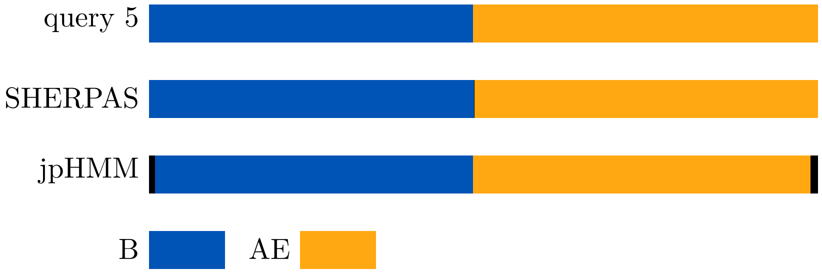
query 5
SHERPAS
jpHMM
B
AE

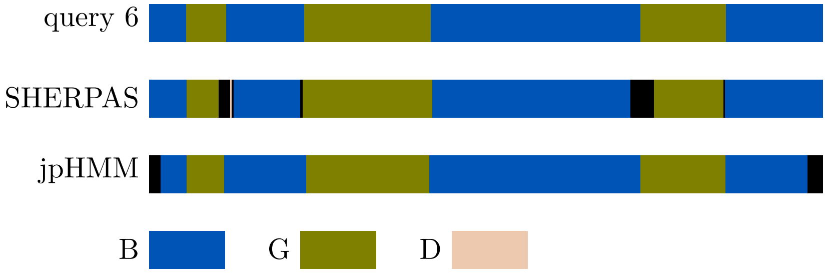
query 6
SHERPAS
jpHMM
B
G
D

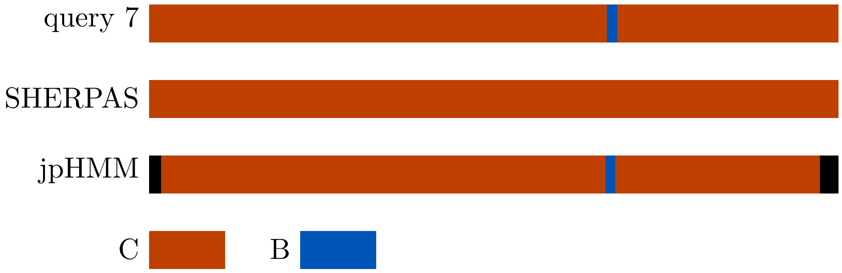
query 7
SHERPAS
jpHMM
C
B

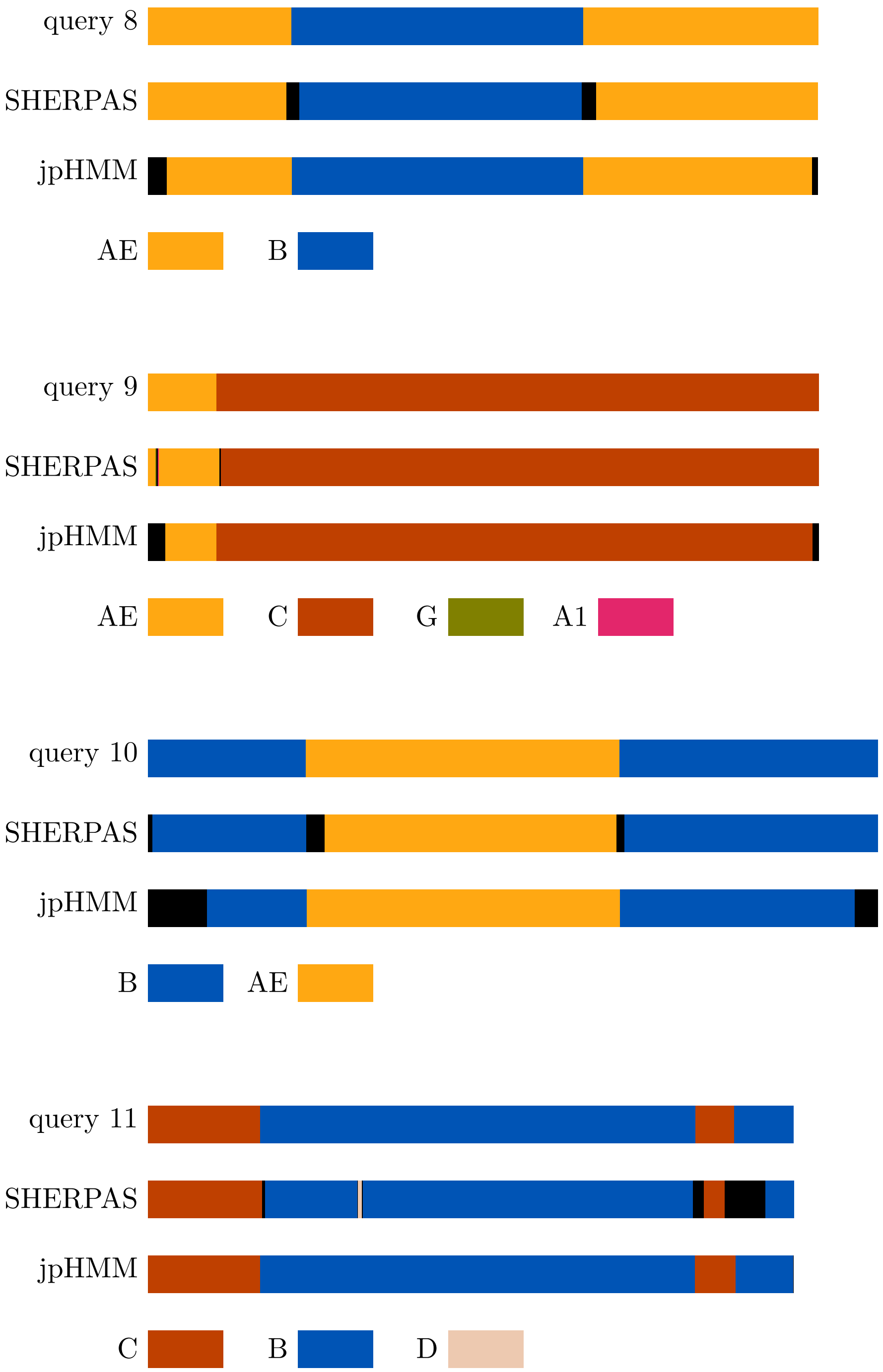
query 8
SHERPAS
jpHMM
AE
B
query 9
SHERPAS
jpHMM
AE
C
G
A1
query 10
SHERPAS
jpHMM
B
AE
query 11
SHERPAS
jpHMM
C
B
D

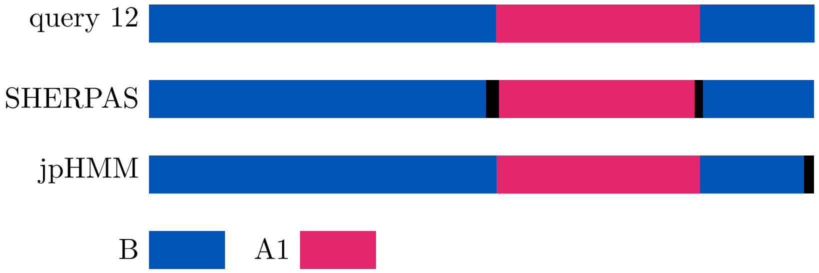
query 12
SHERPAS
jpHMM
B
A1

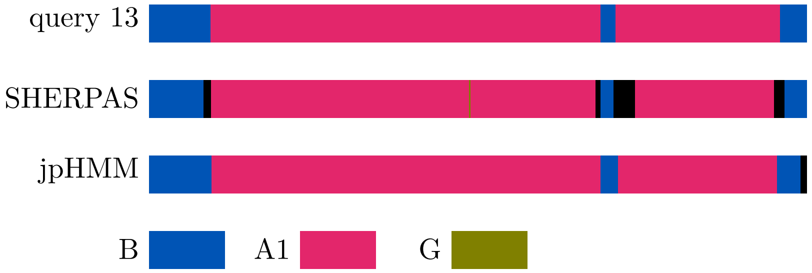
query 13
SHERPAS
jpHMM
B
A1
G

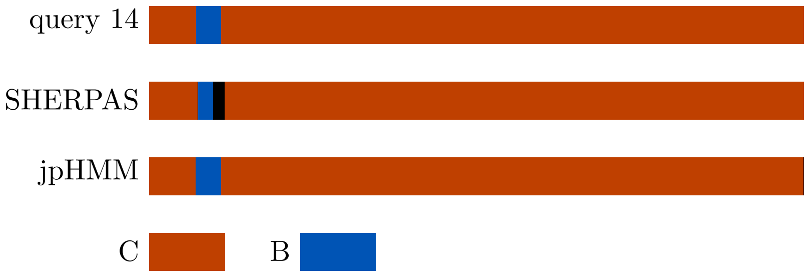
query 14
SHERPAS
jpHMM
C
B

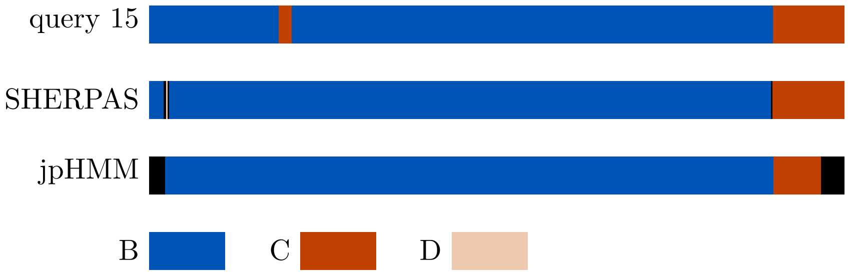
query 15
SHERPAS
jpHMM
B
C
D

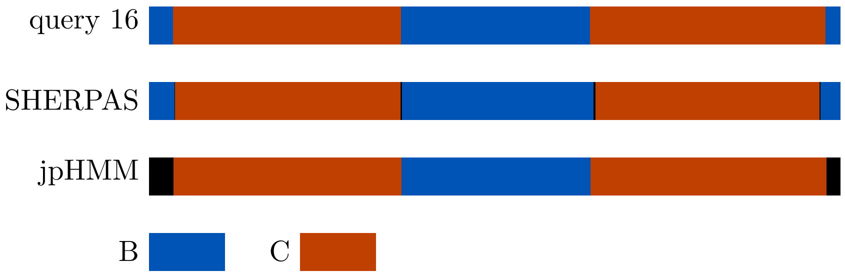
query 16
SHERPAS
jpHMM
B
C

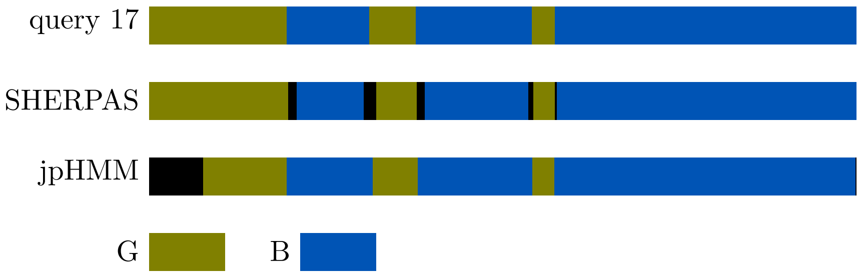
query 17
SHERPAS
jpHMM
G
B

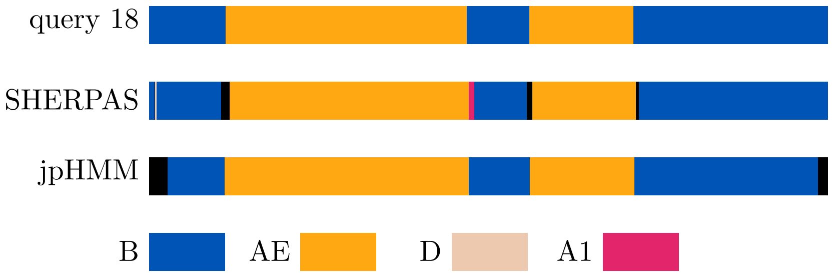
query 18
SHERPAS
jpHMM
B
AE
D
A1

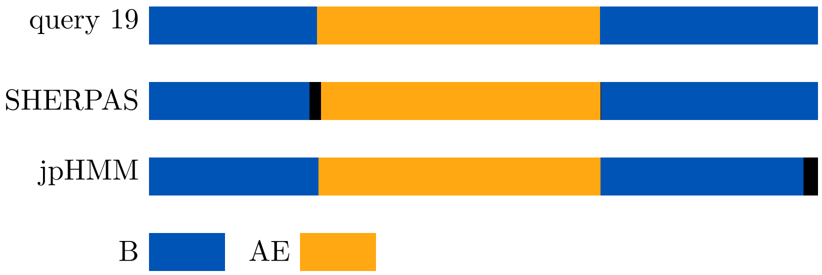
query 19
SHERPAS
jpHMM
B
AE

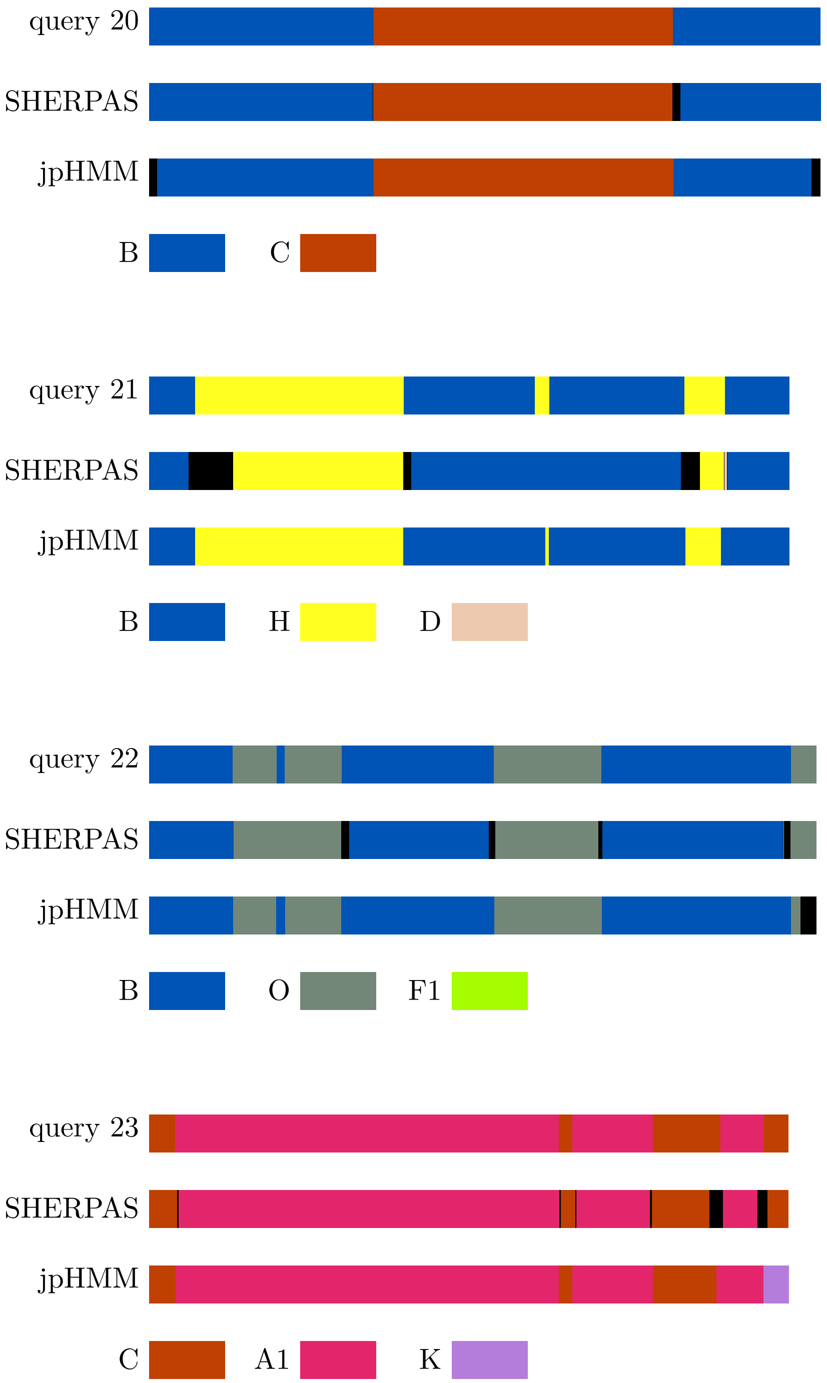
query 20
SHERPAS
jpHMM
B
C
query 21
SHERPAS
jpHMM
B
H
D
query 22
SHERPAS
jpHMM
B
O
F1
query 23
SHERPAS
jpHMM
C
A1
K

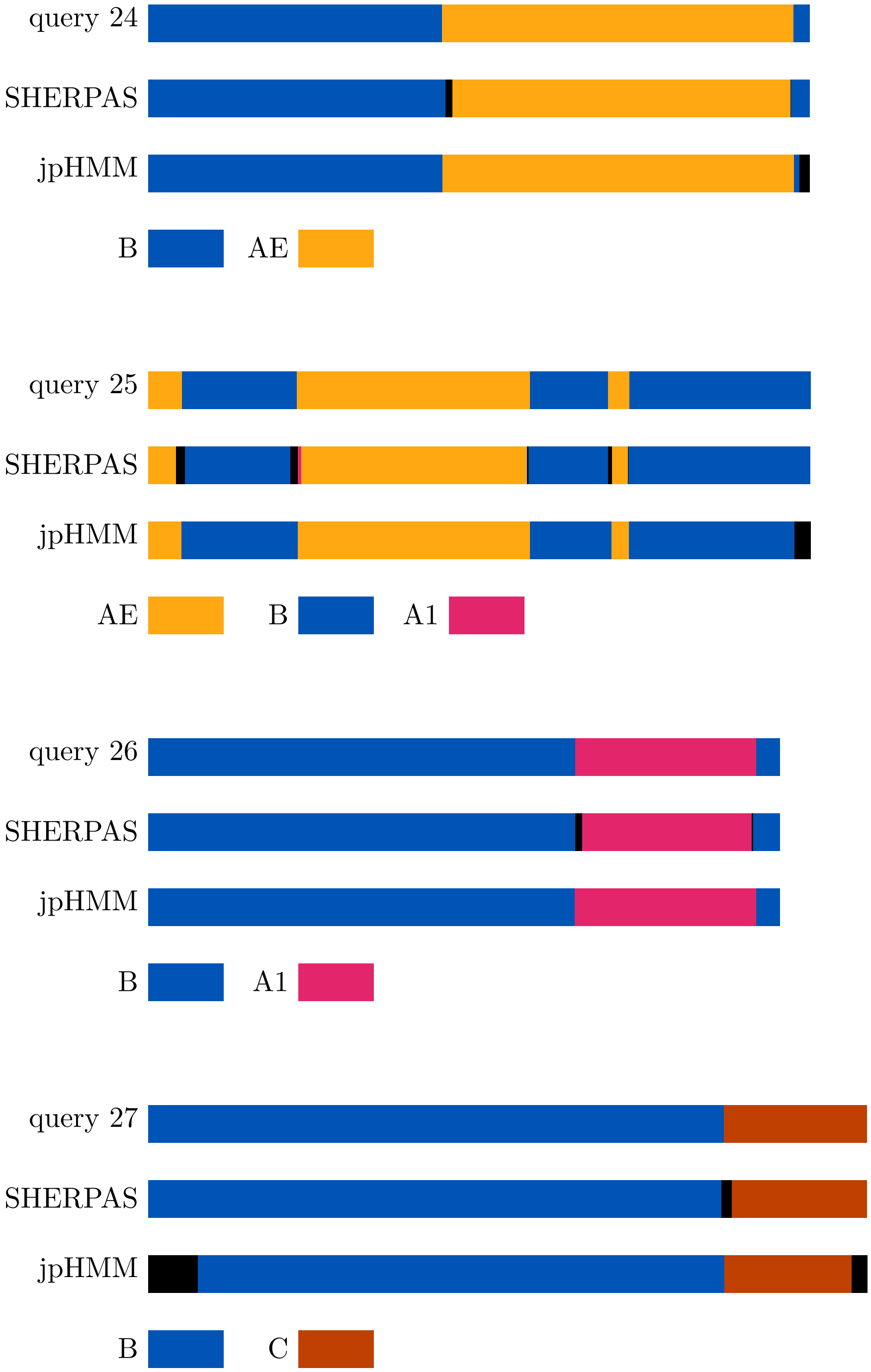
query 24
SHERPAS
jpHMM
B
AE
query 25
SHERPAS
jpHMM
AE
B
A1
query 26
SHERPAS
jpHMM
B
A1
query 27
SHERPAS
jpHMM
B
C

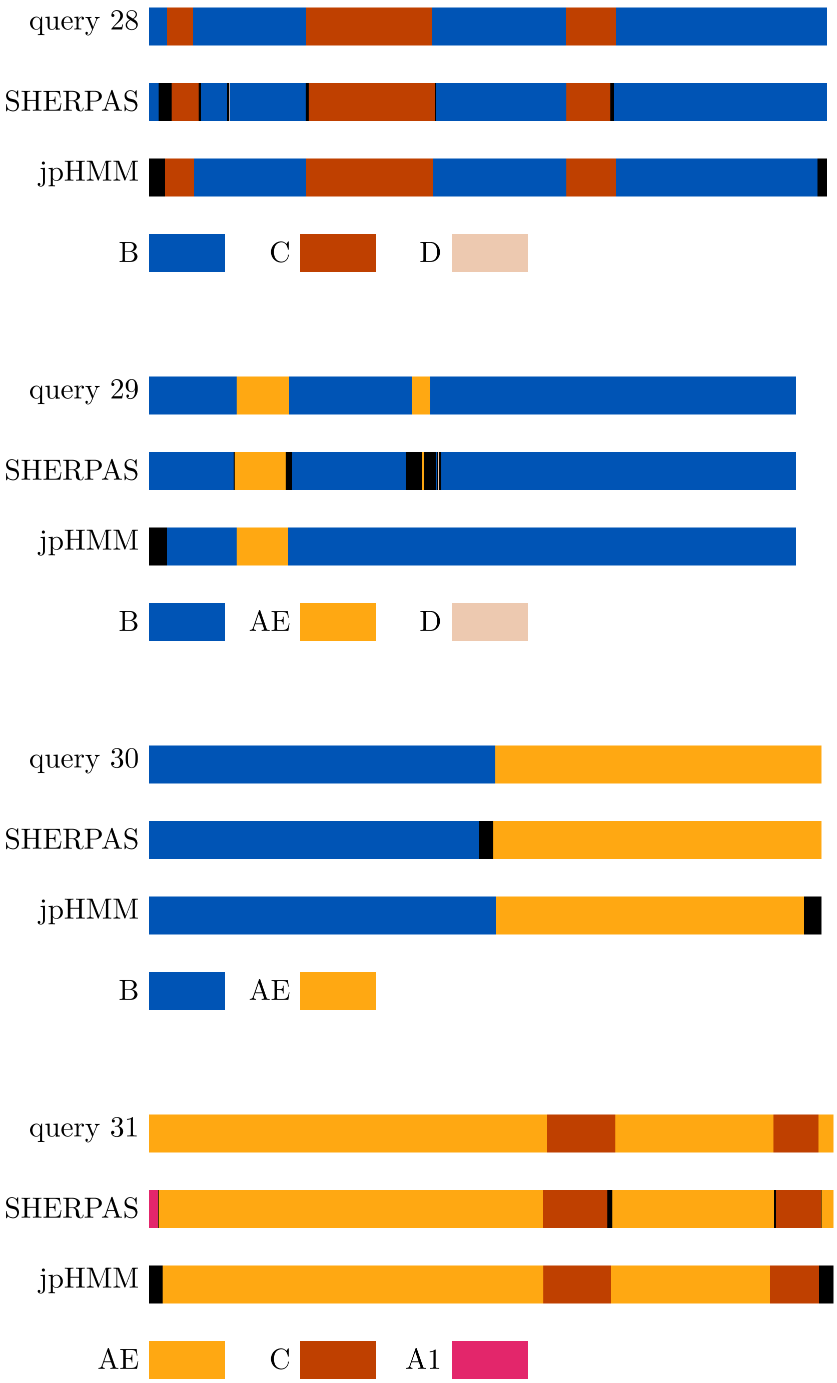
query 28
SHERPAS
jpHMM
B
C
D
query 29
SHERPAS
jpHMM
B
AE
D
query 30
SHERPAS
jpHMM
B
AE
query 31
SHERPAS
jpHMM
AE
C
A1

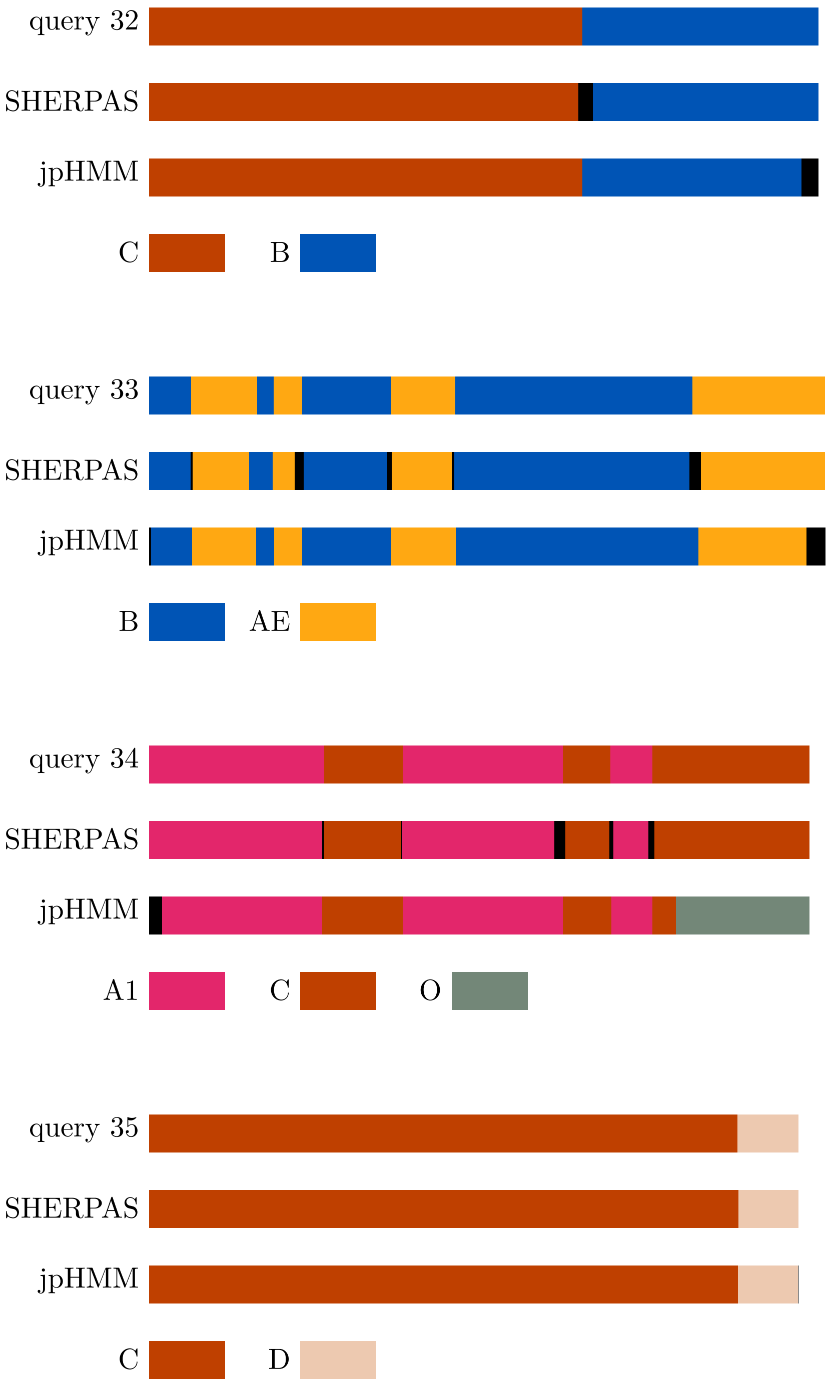
query 32
SHERPAS
jpHMM
C
B
query 33
SHERPAS
jpHMM
B
AE
query 34
SHERPAS
jpHMM
A1
C
O
query 35
SHERPAS
jpHMM
C
D

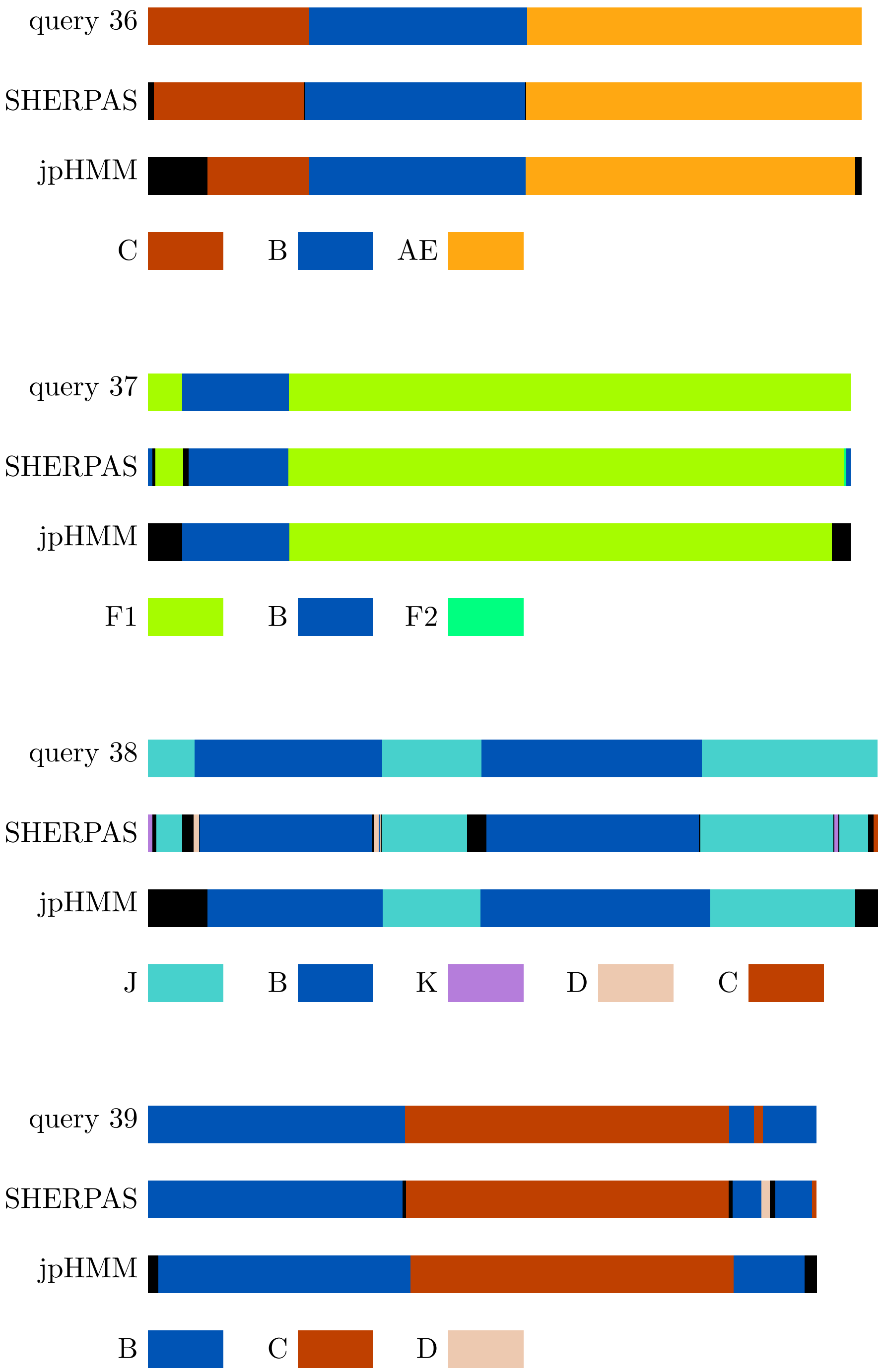
query 36
SHERPAS
jpHMM
C
B
AE
query 37
SHERPAS
jpHMM
F1
B
F2
query 38
SHERPAS
jpHMM
J
B
K
D
C
query 39
SHERPAS
jpHMM
B
C
D

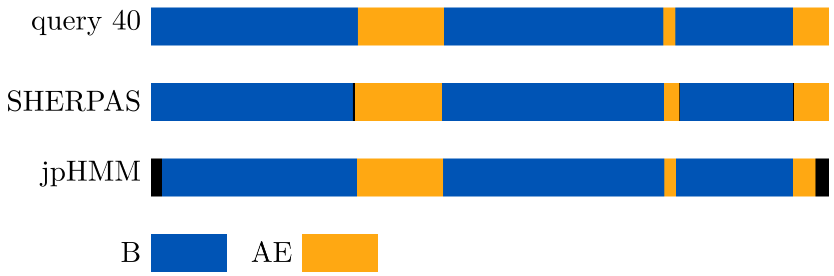
query 40
SHERPAS
jpHMM
B
AE

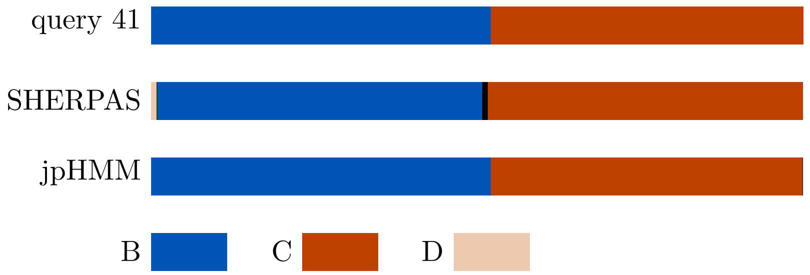
query 41
SHERPAS
jpHMM
B
C
D

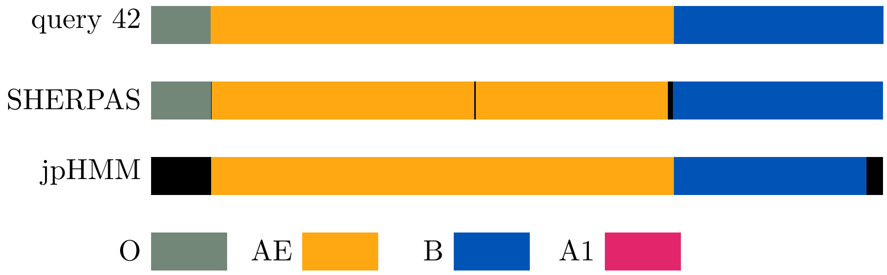
query 42
SHERPAS
jpHMM
O
AE
B
A1

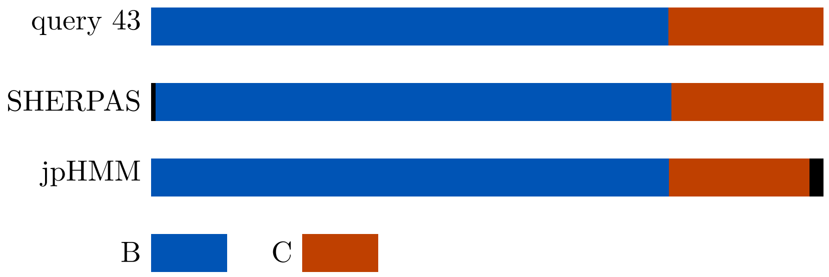
query 43
SHERPAS
jpHMM
B
C

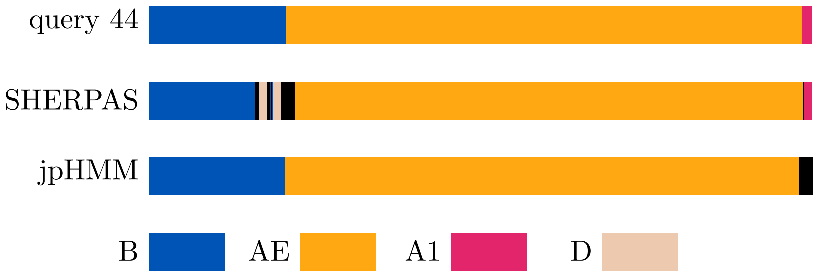
query 44
SHERPAS
jpHMM
B
AE
A1
D

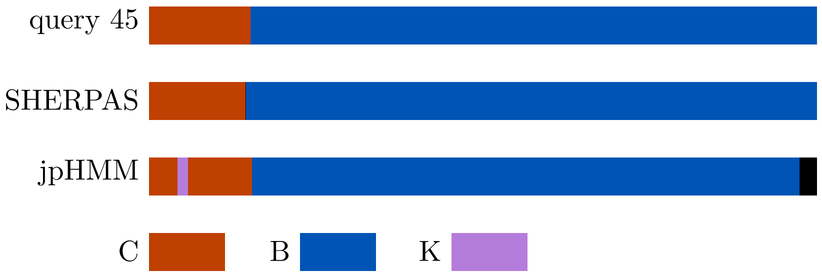
query 45
SHERPAS
jpHMM
C
B
K

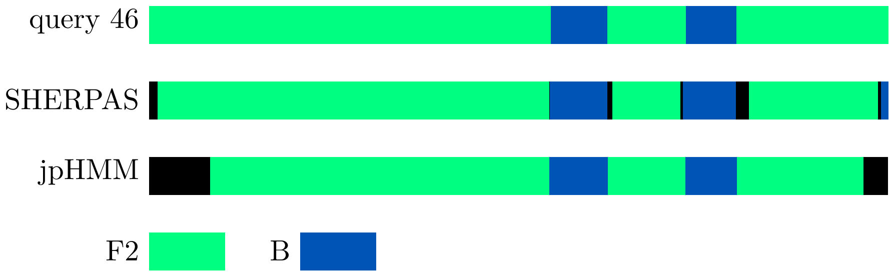
query 46
SHERPAS
jpHMM
F2
B

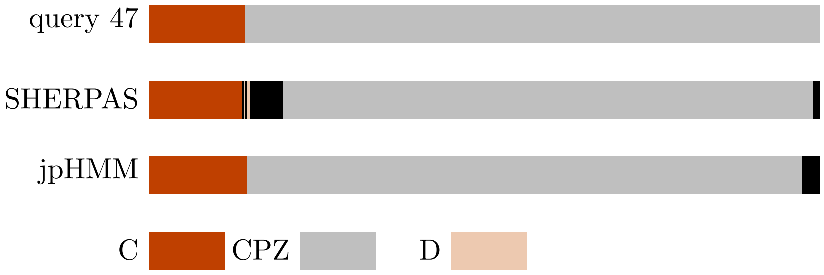
query 47
SHERPAS
jpHMM
C
CPZ
D

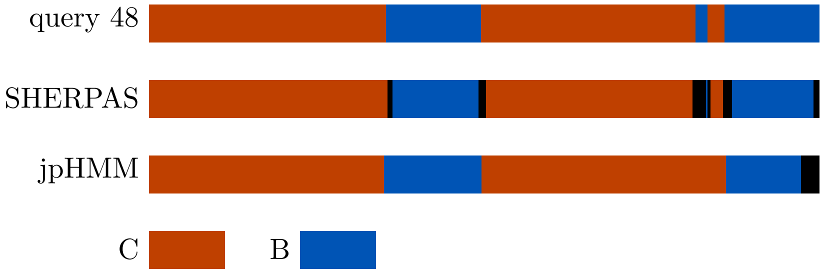
query 48
SHERPAS
jpHMM
C
B

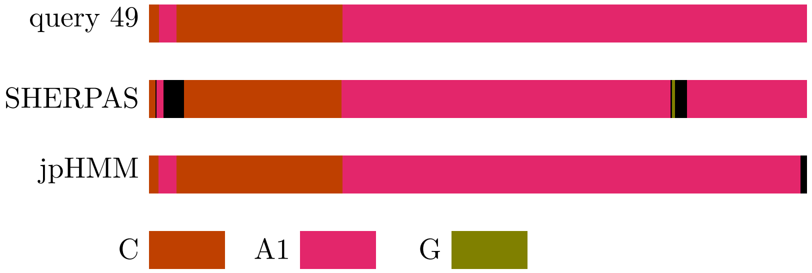
query 49
SHERPAS
jpHMM
C
A1
G

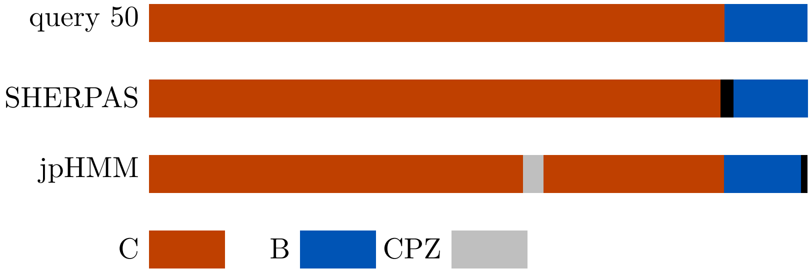
query 50
SHERPAS
jpHMM
C
B
CPZ

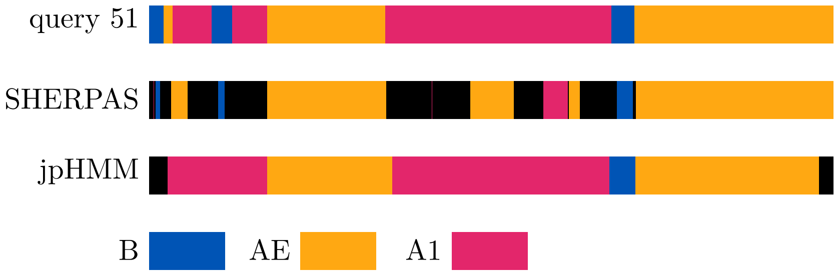
query 51
SHERPAS
jpHMM
B
AE
A1

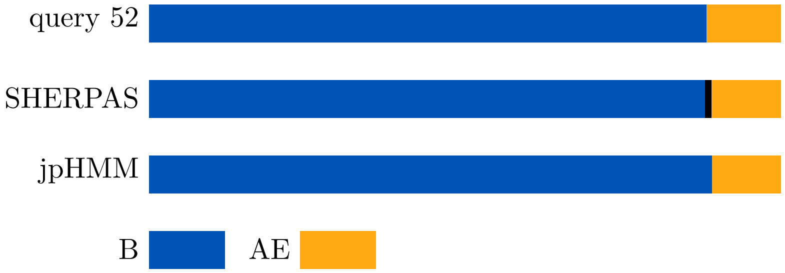
query 52
SHERPAS
jpHMM
B
AE

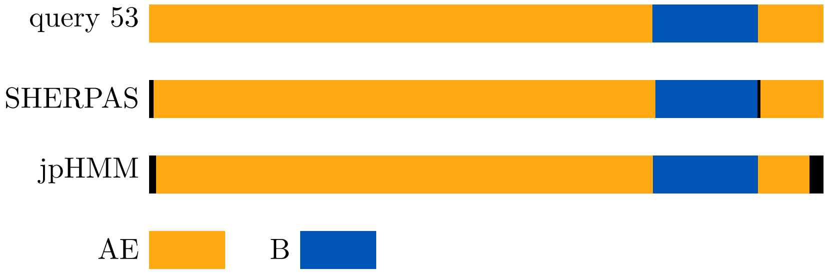
query 53
SHERPAS
jpHMM
AE
B

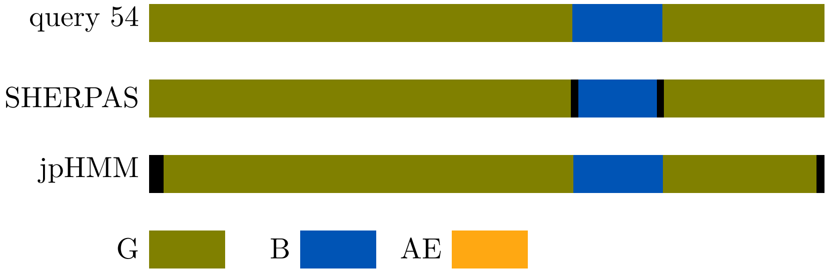
query 54
SHERPAS
jpHMM
G
B
AE

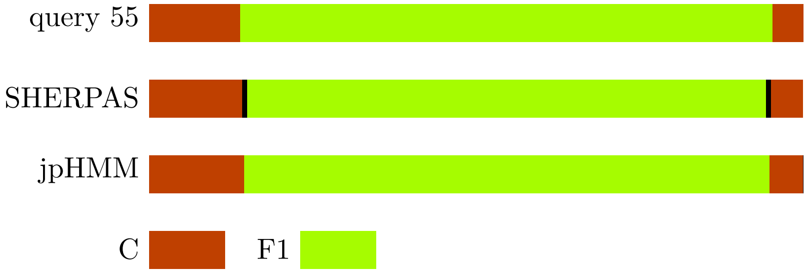
query 55
SHERPAS
jpHMM
C
F1

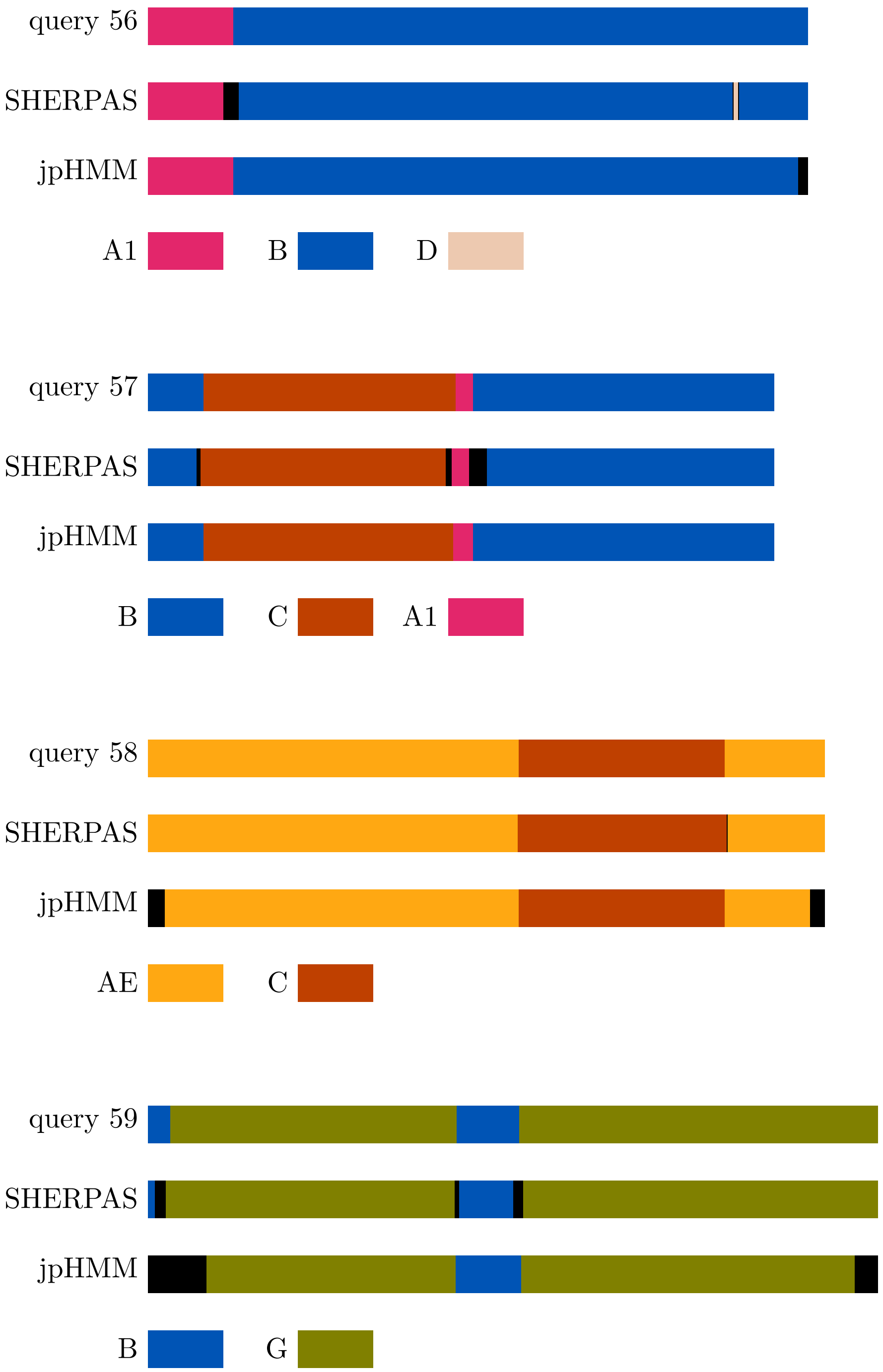
query 56
SHERPAS
jpHMM
A1
B
D
query 57
SHERPAS
jpHMM
B
C
A1
query 58
SHERPAS
jpHMM
AE
C
query 59
SHERPAS
jpHMM
B
G

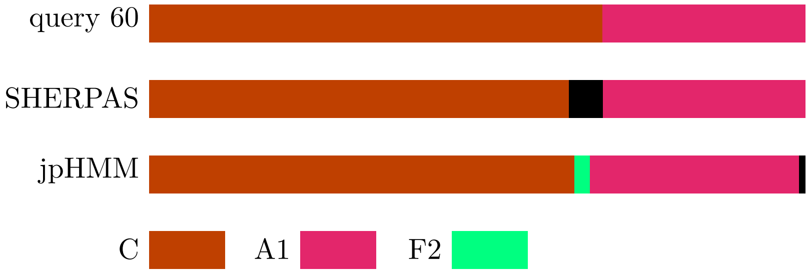
query 60
SHERPAS
jpHMM
C
A1
F2

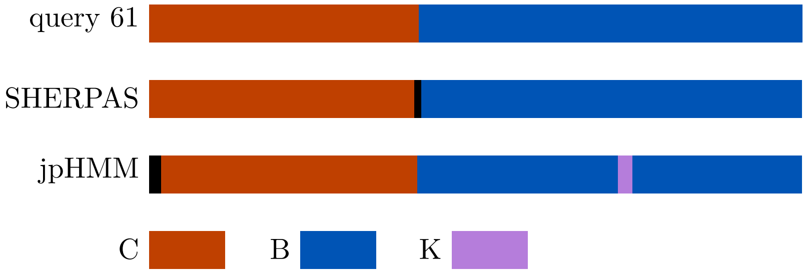
query 61
SHERPAS
jpHMM
C
B
K

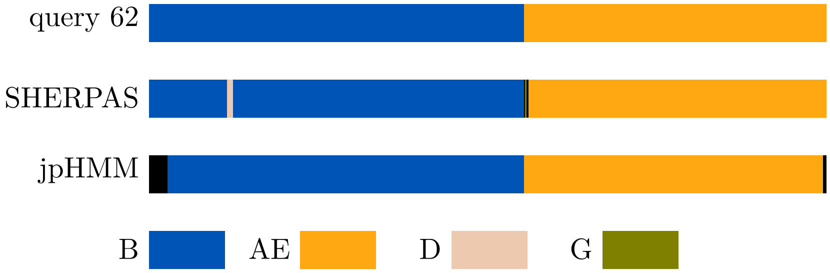
query 62
SHERPAS
jpHMM
B
AE
D
G

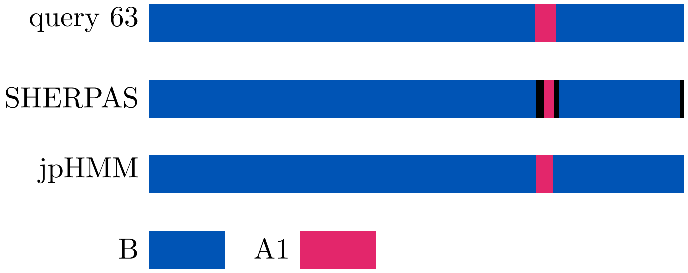
query 63
SHERPAS
jpHMM
B
A1

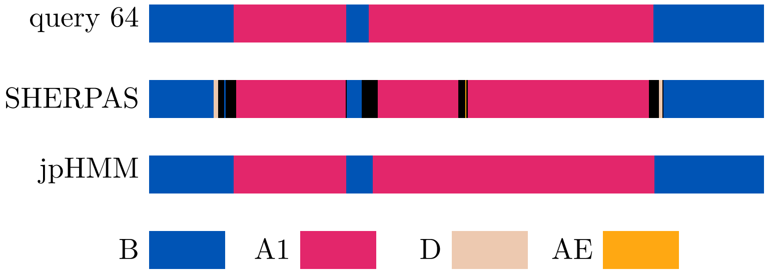
query 64
SHERPAS
jpHMM
B
A1
D
AE

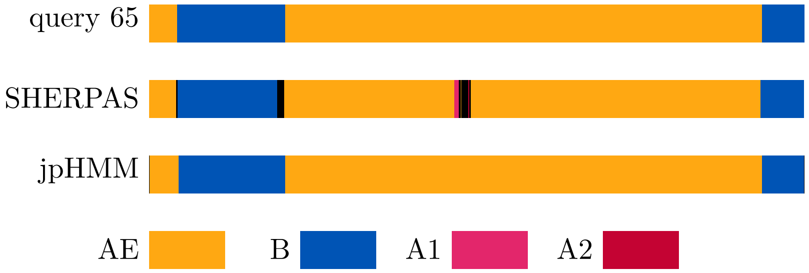
query 65
SHERPAS
jpHMM
AE
B
A1
A2

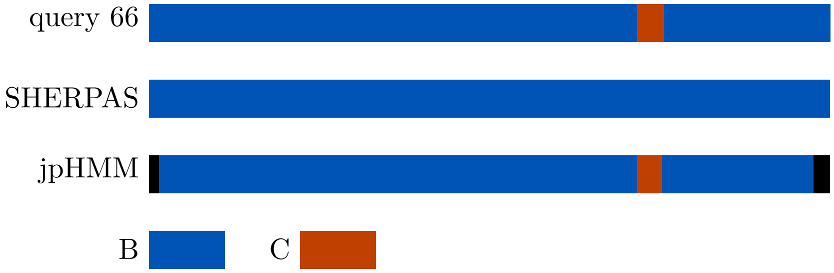
query 66
SHERPAS
jpHMM
B
C

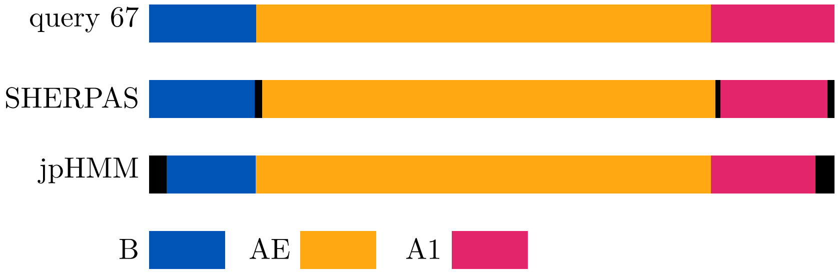
query 67
SHERPAS
jpHMM
B
AE
A1

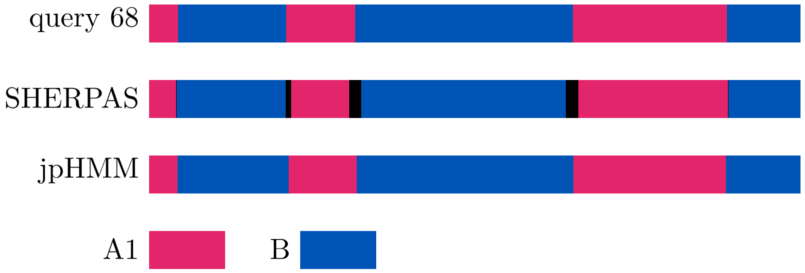
query 68
SHERPAS
jpHMM
A1
B

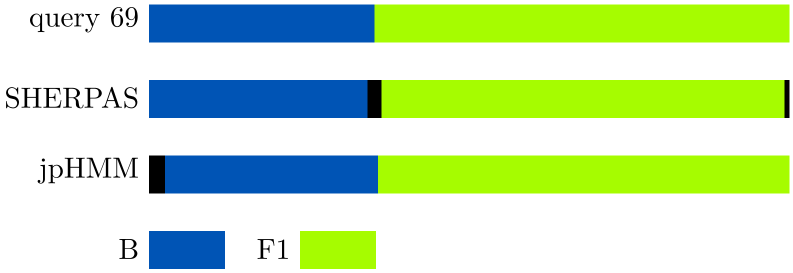
query 69
SHERPAS
jpHMM
B
F1

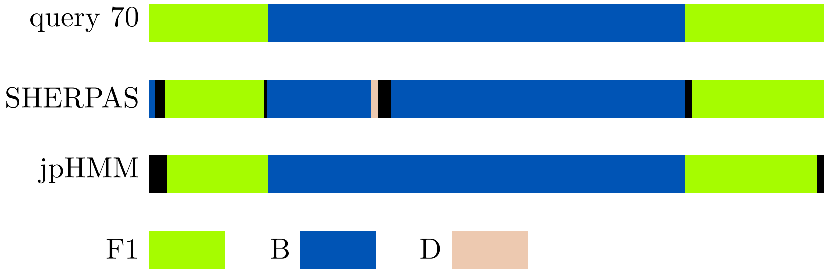
query 70
SHERPAS
jpHMM
F1
B
D

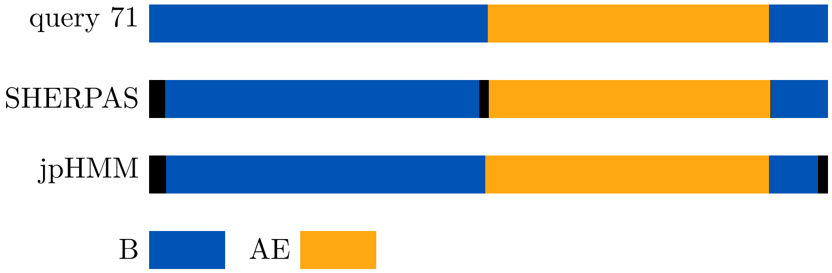
query 71
SHERPAS
jpHMM
B
AE

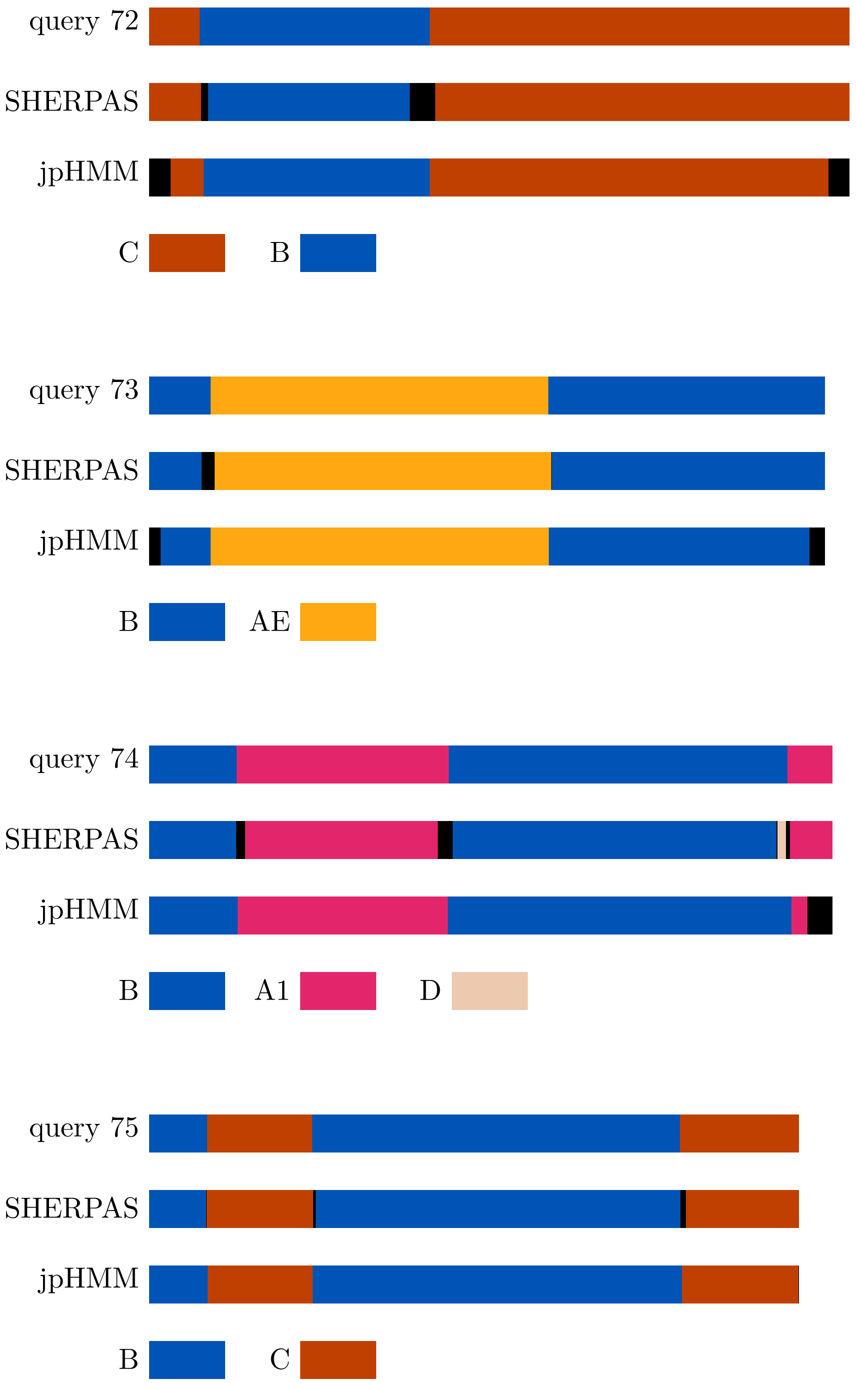
query 72
SHERPAS
jpHMM
C
B
query 73
SHERPAS
jpHMM
B
AE
query 74
SHERPAS
jpHMM
B
A1
D
query 75
SHERPAS
jpHMM
B
C

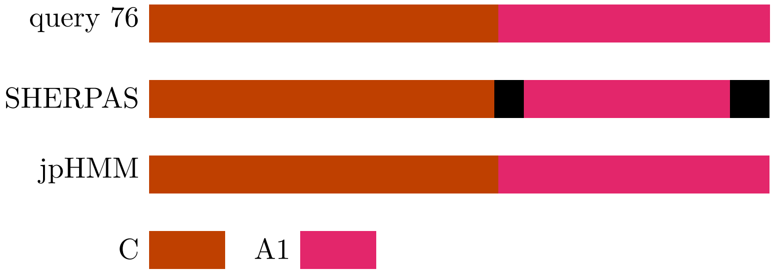
query 76
SHERPAS
jpHMM
C
A1

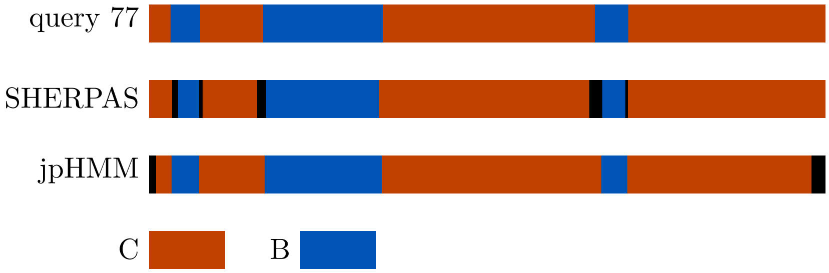
query 77
SHERPAS
jpHMM
C
B

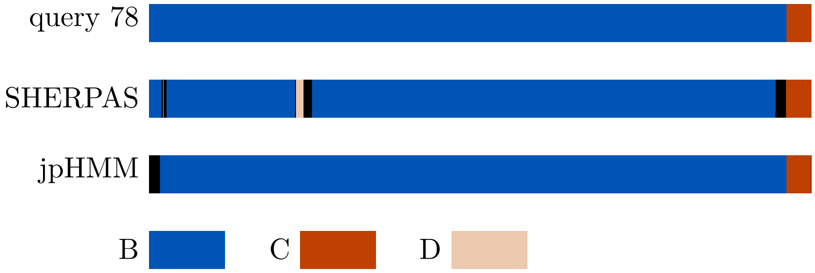
query 78
SHERPAS
jpHMM
B
C
D

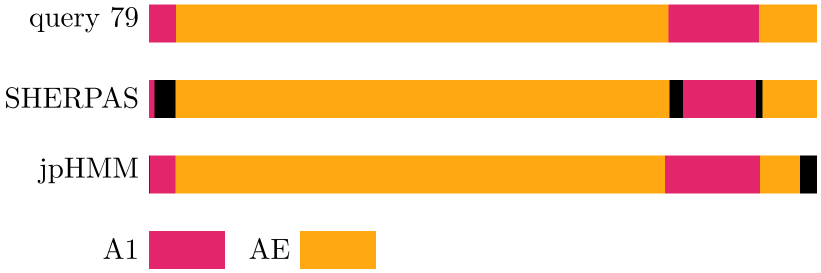
query 79
SHERPAS
jpHMM
A1
AE

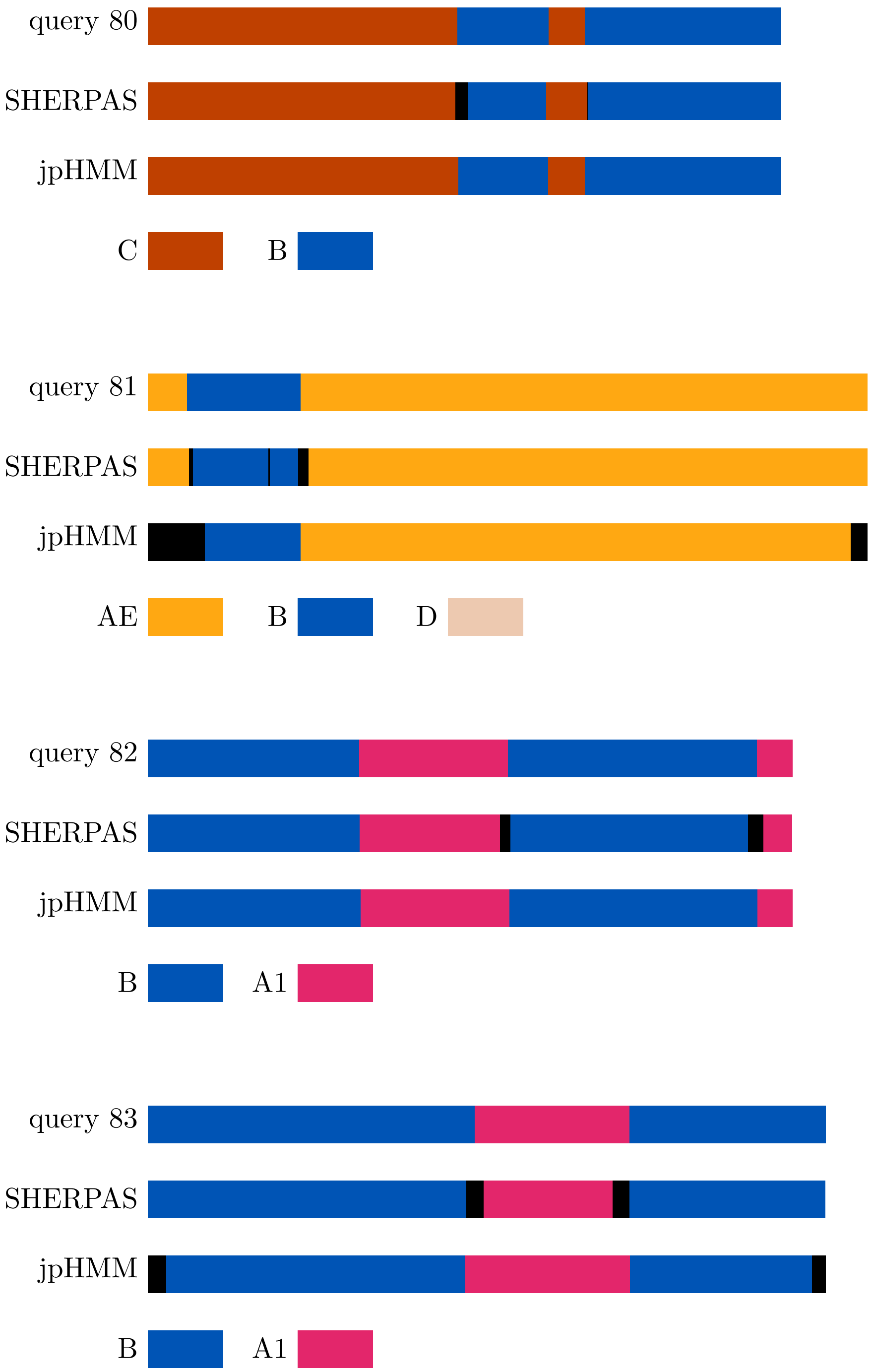
query 80
SHERPAS
jpHMM
C
B
query 81
SHERPAS
jpHMM
AE
B
D
query 82
SHERPAS
jpHMM
B
A1
query 83
SHERPAS
jpHMM
B
A1

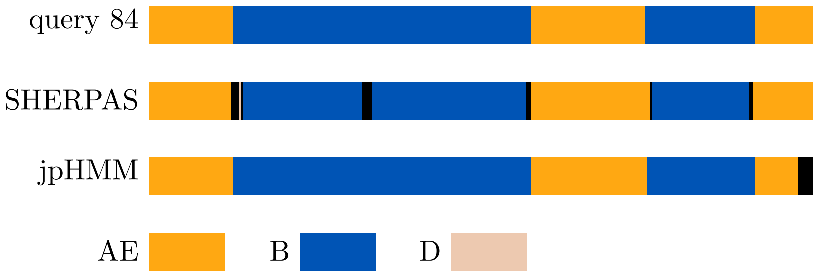
query 84
SHERPAS
jpHMM
AE
B
D

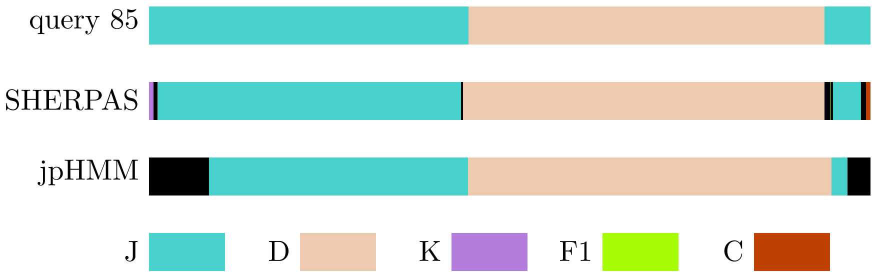
query 85
SHERPAS
jpHMM
J
D
K
F1
C

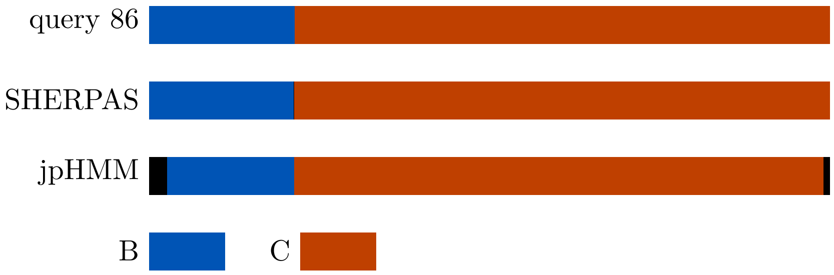
query 86
SHERPAS
jpHMM
B
C

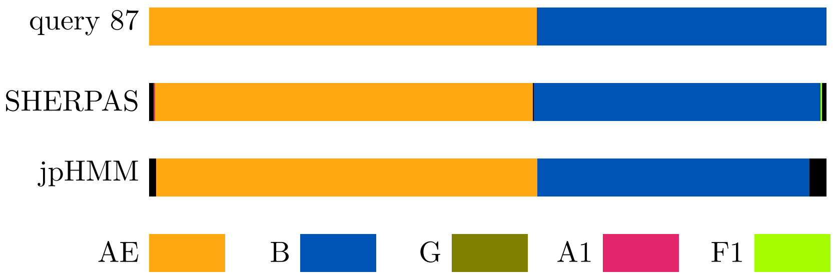
query 87
SHERPAS
jpHMM
AE
B
G
A1
F1

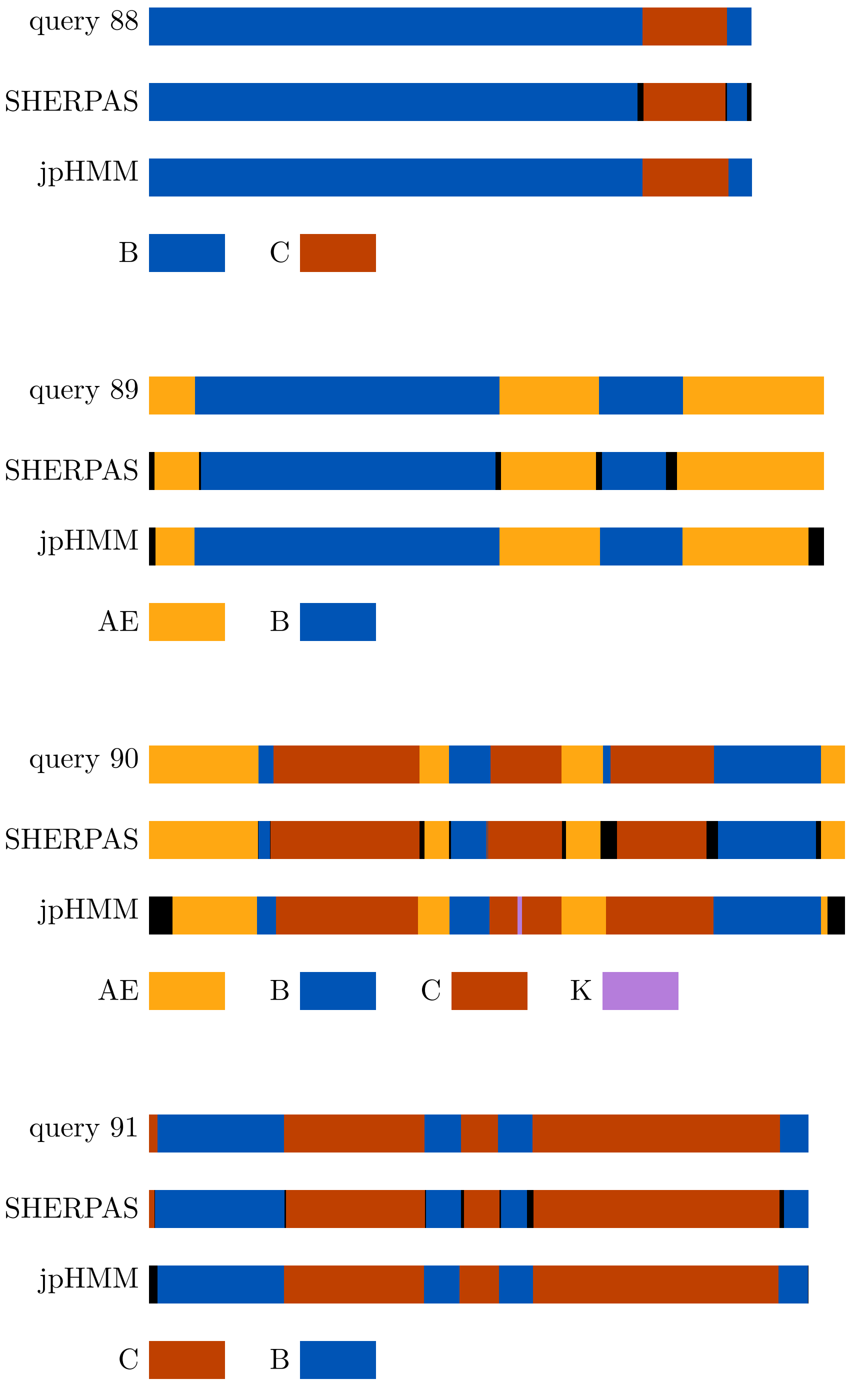
query 88
SHERPAS
jpHMM
B
C
query 89
SHERPAS
jpHMM
AE
B
query 90
SHERPAS
jpHMM
AE
B
C
K
query 91
SHERPAS
jpHMM
C
B

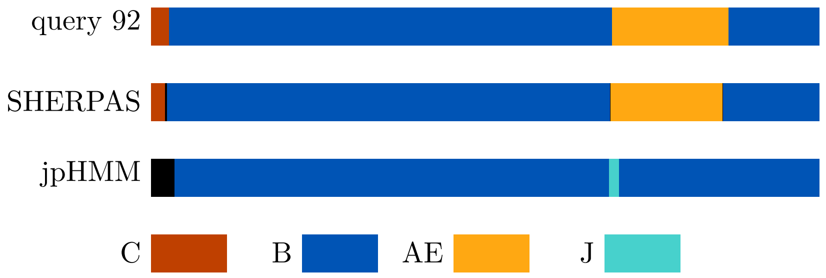
query 92
SHERPAS
jpHMM
C
B
AE
J

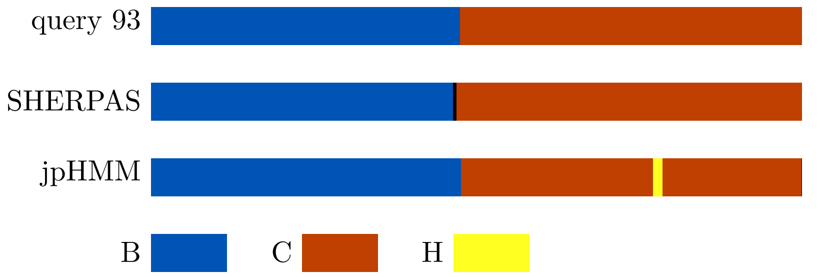
query 93
SHERPAS
jpHMM
B
C
H

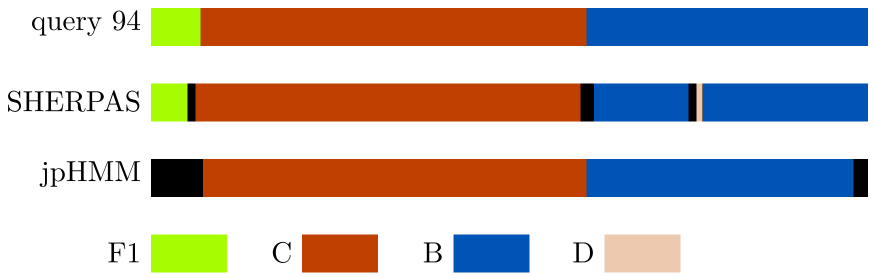
query 94
SHERPAS
jpHMM
F1
C
B
D

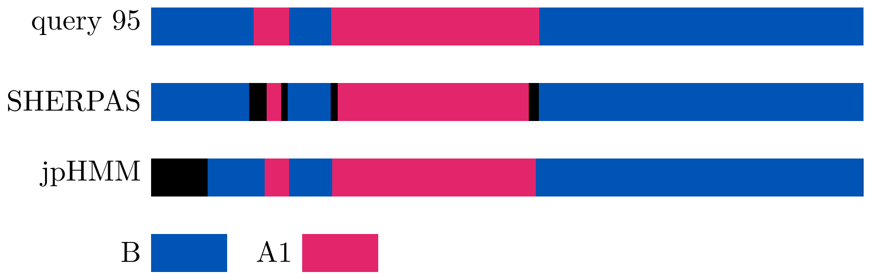
query 95
SHERPAS
jpHMM
B
A1

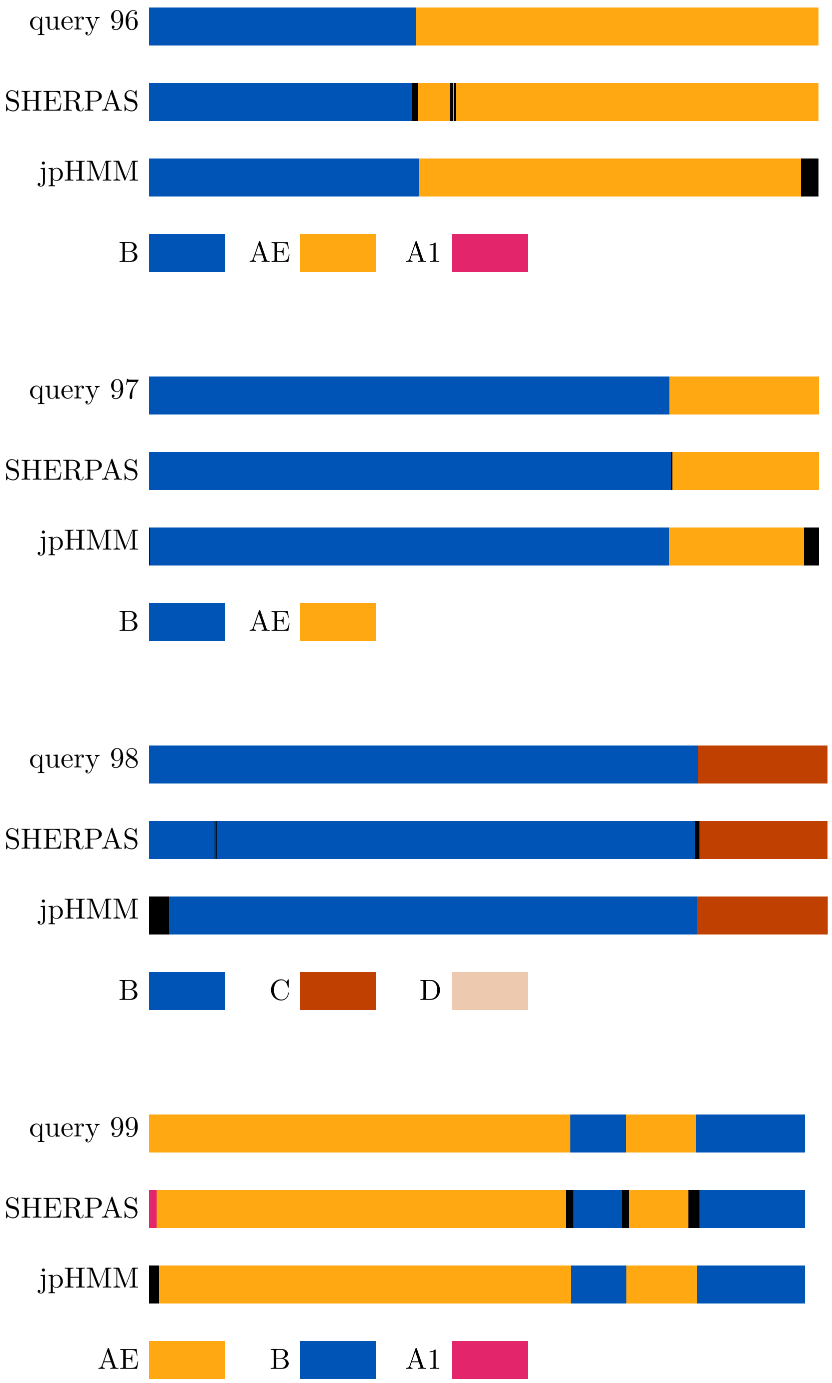
query 96
SHERPAS
jpHMM
B
AE
A1
query 97
SHERPAS
jpHMM
B
AE
query 98
SHERPAS
jpHMM
B
C
D
query 99
SHERPAS
jpHMM
AE
B
A1

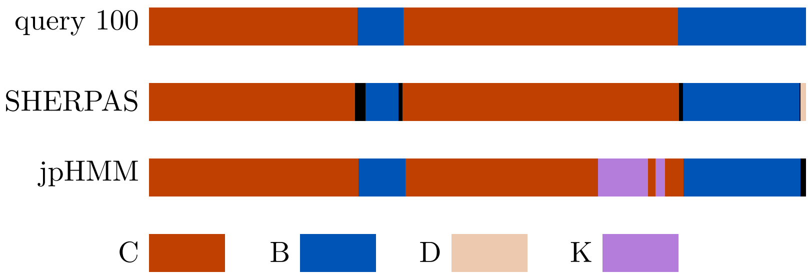
query 100
SHERPAS
jpHMM
C
B
D
K
